# Harmonized-Multinational qEEG Norms (HarMNqEEG)

**DOI:** 10.1101/2022.01.12.476128

**Authors:** Min Li, Ying Wang, Carlos Lopez-Naranjo, Aini Ismafairus Abd Hamid, Alan C. Evans, Alexander N. Savostyanov, Ana Calzada-Reyes, Ariosky Areces-Gonzalez, Arno Villringer, Carlos A. Tobon-Quintero, Daysi Garcia-Agustin, Deirel Paz-Linares, Dezhong Yao, Li Dong, Eduardo Aubert-Vazquez, Faruque Reza, Hazim Omar, Jafri Malin Abdullah, Janina R. Galler, John F. Ochoa-Gomez, Leslie S. Prichep, Lidice Galan-Garcia, Lilia Morales-Chacon, Mitchell J. Valdes-Sosa, Marius Tröndle, Mohd Faizal Bin Mohd Zulkifly, Muhammad Riddha Bin Abdul Rahman, Natalya S. Milakhina, Nicolas Langer, Pavel Rudych, Shiang Hu, Thomas Koenig, Trinidad A. Virues-Alba, Xu Lei, Maria L. Bringas-Vega, Jorge F. Bosch-Bayard, Pedro Antonio Valdes-Sosa

## Abstract

This paper extends our frequency domain quantitative electroencephalography (qEEG) methods pursuing higher sensitivity to detect Brain Developmental Disorders. Prior qEEG work lacked integration of cross-spectral information omitting important functional connectivity descriptors. Lack of geographical diversity precluded accounting for site-specific variance, increasing qEEG nuisance variance. We ameliorate these weaknesses. i) Create lifespan Hermitian Riemannian multinational qEEG norms for cross-spectral tensors. These norms result from the HarMNqEEG project fostered by the Global Brain Consortium. We calculate the norms with data from 9 countries, 12 devices, and 14 studies, including 1564 subjects. Instead of raw data, only anonymized metadata and EEG cross-spectral tensors were shared. After visual and automatic quality control developmental equations for the mean and standard deviation of qEEG traditional and Hermitian Riemannian descriptive parameters were calculated using additive mixed-effects models. We demonstrate qEEG “batch effects” and provide methods to calculate harmonized z-scores. ii) We also show that the multinational harmonized Hermitian Riemannian norms produce z-scores with increased diagnostic accuracy to predict brain dysfunction at school-age produced by malnutrition only in the first year of life. We provide data and software for constructing norms. iii) We offer open code and data to calculate different individual z-scores from the HarMNqEEG dataset. These results contribute to developing bias-free, low-cost neuroimaging technologies applicable in various health settings.

**Highlights:**

- We create lifespan Hermitian Riemannian qEEG norms for cross-spectral tensors.
- The norms are based on 9 countries, 12 devices, and 14 studies, with 1564 subjects.
- We demonstrate qEEG “batch effects”, providing harmonization methods to remove them.
- Multinational harmonized z-scores increase diagnostic accuracy of brain dysfunction.
- Data and software are available for norm and individual z-scores calculation.

## Introduction

Characterizing the age-dependent developmental trajectories of the brains of healthy and dSiseased individuals is essential for precision medicine. As Verdi et al.(2021) recently emphasized, when using Neuroimaging for this purpose, we must move beyond the “average patient”; we must instead understand individual differences in brain aging processes to allow early detection of functional deterioration and neurodegenerative disease. Quantifying these individual developmental trajectories hinges on choosing the proper “Descriptive Parameters” (DPs) that summarize brain anatomy or physiology features and distinguishing their normal or abnormal evolution during the lifespan. These DPs depend strongly on age.

Central to the definition of healthy developmental brain trajectories is the creation of age-dependent developmental “norms” (“charts” or “atlases”) comprising both measures of central population tendency as well as dispersion (Ahn et al., 1980). The norms enable quantifying the age-adjusted statistical distance of a subject’s DP from the healthy population. Examples of such distances are the z-score or Mahalanobis distance(John et al., 1988). Armed with these age-dependent statistical distances, we can quantify Brain Developmental Deviation (BDD) and even use them to cluster and stage disease progressions(Harmony, 2021).

Large multinational projects to develop norms are now underway. These projects aim to increase genotypic and phenotypic diversity, and importantly, to achieve sample sizes to provide adequate statistical power (Bethlehem et al., 2021). These endeavors have a fundamental limitation due to the costly nature and sparse geographical distribution of the technologies used to probe the brain. Large parts of the world population, even in high-income countries, are difficult or impossible to sample. Thus issues of fairness and racial bias cannot be ignored.

In contrast with other neurotechnologies, electroencephalography (EEG) is affordable, portable, and deployable in all health system levels--whatever the economic setting. Thus EEG is a potential tool for detecting BDD in a Global Health context (Valdés-Sosa et al., 2021). Quantitative EEG (qEEG) facilitates this use by using EEG-based DPs to compare individuals with qEEG norms. The most common embodiment of qEEG uses the EEG log-power spectrum as DPs. The seminal work of Matousek and Petersen (1973a) pioneered this work 50 years ago, with the visionary introduction of the “age-dependent EEG quotients” to measure brain age—antedating by 4 decades current interest in this topic! This line of work John et al.(1977), Harmony et al. (1988), Bosch-Bayard et al. (2020), Hernandez-Gonzalez et al. (2011) subsequently systematized this initiative. Consequently, developmental norms were constructed in several countries (Gordon et al., 2005; Lorensen and Dickson, 2003; Thatcher et al., 2003). Other projects recently vigorously launched are being repurposed for normative work (Pavlov et al., 2021).

An instance of norm construction and evaluation of BDD in a lower or middle-income country (LMIC) has been the Cuban Human Brain Mapping Project (CHBMP). Its first wave provided norms (means and standard deviations (SDs)) for the narrowband log-spectral DP based on 211 subjects from age 5 to 97. Despite being based on a single country database, CHBMP norms describe BDD consistently in other countries (Bosch-Bayard et al., 2001; Bringas Vega et al., 2019; Taboada-Crispi et al., 2018). Nevertheless, the relatively small sample sizes make country comparisons relatively underpowered compared to neuroimaging efforts such as ENIGMA (Thompson et al., 2014).

The lack of global inclusiveness and small sample sizes in qEEG norms is a situation that this paper attempts to ameliorate by constructing a multinational norm based on 1586 EEGs, 9 countries, and 12 EEG devices. We used the novel collaboration strategy described in (Gazula et al., 2020) to facilitate data-sharing. Each site did not share raw data but rather processed it with standard software. The only information shared for collaboration was anonymized data and the EEG cross-spectrum of each subject.

The diversity of countries and sites brought the problem of harmonization to the forefront. Harmonization is the elimination of “batch effects.” A Batch effect is a nuisance variance due to cross-site, equipment differences, and changes over time of parameters of experiments that purport to measure the same underlying biological mechanisms. Genomics was the first to identify and minimize batch effects with statistical techniques. One such well-known technique is COMBAT, described in (Johnson et al., 2007). Subsequently, MRI multisite studies identified a similar problem where batch effects may be due to different acquisition systems, variations in protocols. Recent efforts have been to address batch effects have gained traction in neuroimaging (Fortin et al., 2018; Pomponio et al., 2020; Rutherford et al., 2021a). To our knowledge, there have been no systematic studies of batch effects in qEEG.

A multinational EEG norm faces an even greater need for harmonization than MRI, the variability of recording systems from different vendors compounded by the lack of standards. Different amplifier transfer functions, electrode placement systems, preprocessing protocols beg the question of EEG batch effects. Despite the apparent need for harmonization, to the best of our knowledge, there are no statistical studies of batch effects in qEEG. Therefore, we propose new statistical techniques adapted to the nature of EEG spectra for this purpose and try to identify what variables effectively define qEEG batches, testing their effect on the resulting batch harmonized norms and the final practical impact on measuring BDD.

We shall also remedy a current difficulty with qEEG norms. As mentioned before, these are predominantly either for broad-band (BB) sensor space log-spectra DPs (Ahn et al., 1980; John, 1987) or their narrowband (NB) version (Valdés et al., 1992). This preference for log-spectra ignores the that all second-order (linear) properties of quasi-stationary EEGs are encoded in the full cross-spectrum, a tensor of Hermitian frequency-dependent matrices. The diagonal of each matrix contains the spectra for that frequency. Using only that information for qEEG ignores the off-diagonal elements, which are the cross-spectra

To explain the geometric relation of spectra and cross-spectra, we remind the reader of recent statistical developments for data laying on Riemannian spaces. This theory provides the unified framework to deal with cross-spectrum in a principled way. Proper distances to consider are not Euclidean as in usual multivariate statistics cross-spectrum, but somewhat related concepts for Riemann manifolds (Karcher, 1977; Pennec, 2004; Sabbagh et al., 2019).

Why is Riemannian geometry important for qEEG? Cross-spectra are positive-definite Hermitian (HPD) matrices that live in a nonlinear manifold which is a analog of Riemannian geometry. Previous efforts to construct qEEG norms used the logarithm as a transformation towards Gaussianity and then followed up with ordinary univariate statistics cross-spectrum to construct norms. In the Riemannian framework, the matrix logarithm transforms the whole cross-spectral matrix towards multivariate Gaussianity (Riemannian vectorization). The approximate multivariate gaussian distribution of log matrix covariance matrices has a long history (Leonard and Hsu, 2001).

Riemannian geometry has become popular in the Brain-Computer Interface literature, with significant advantages for classification precision (Yger et al., 2017). Sabbagh et al. (2020) recently showed Riemannian methods to improve brain age estimation with MEG. However, to our knowledge, the construction of Riemannian geometry-based developmental norms for either EEG or MEG has not been attempted. We remedy this situation in this paper, constructing harmonized norms for the Riemann vectorized cross-spectra and investigating the existence of batch effects for this type of multinational data. We also gauge the practical effect of these technical refinements on discriminant equations between out-of-sample controls and pathological subjects.

We alert the reader that our previous norms comprised scalp and source space log spectra (Bosch-Bayard et al., 2001). The excessive computational complexity of a Riemannian source cross-spectral analysis is out of the present paper’s scope that we postpone to future work.

This paper is organized as follows: Sections 1–4 deal with the methods used to construct the norms, with the treatment of 1) the theoretical basis of traditional and Hermitian Riemannian qEEG Descriptive Parameters; 2) Gathering data and preprocessing for the multinational qEEG norm project; 3) Construction of and presentation of the harmonized norms; 4) Quality control of the project; 5) Validation of the use of the norms in detection BDD.

## 1 The theoretical basis of qEEG descriptive parameters

The basic mathematical symbols used in this paper are shown in table I and the summary of operators in Table II. The summary of notation for independent variables is shown in Table A-I.

**Table I:**
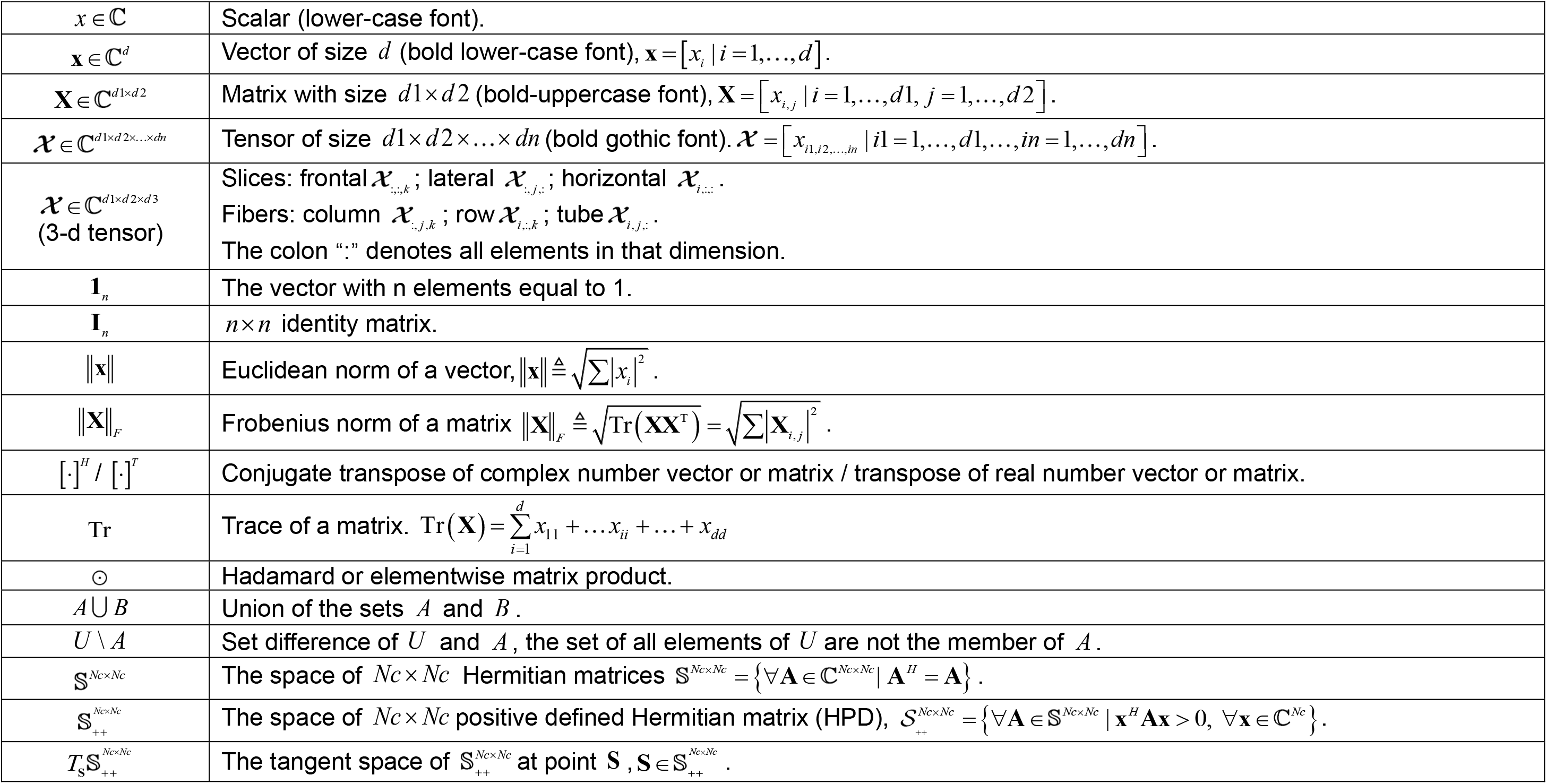
Basic mathematical symbols.

**Table II:**
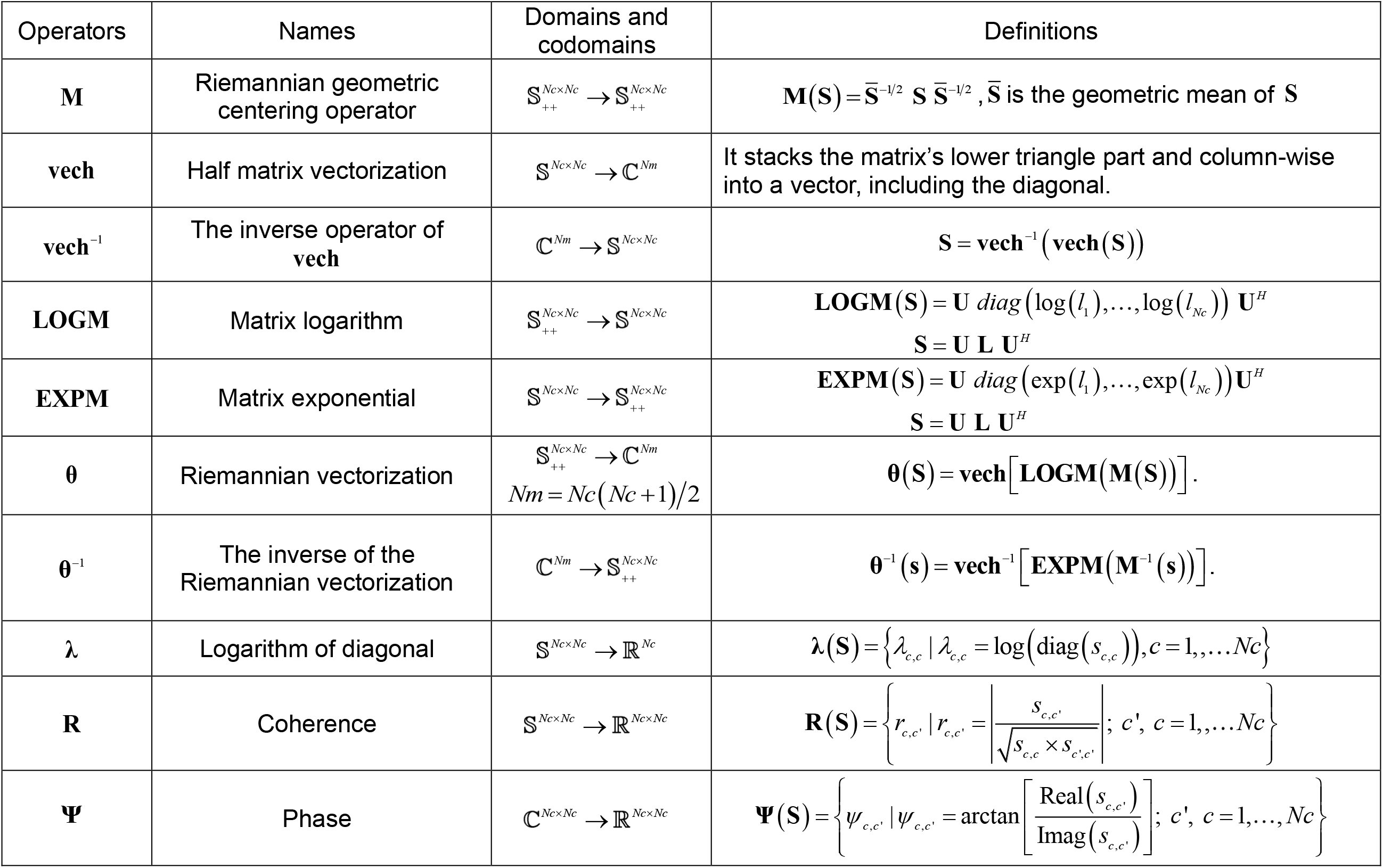
Operator definitions

### 1.1 Descriptive Parameters based on EEG cross-spectrum

Quantitative electroencephalography (qEEG) studies descriptive parameters (DPs) obtained from the electroencephalogram (EEG). These DPs encode physiologically relevant information. Here we explain the frequency domain DPs defined with the EEG cross-spectrum. Nonlinear frequency-domain (Billings and Tsang, 1989a, 1989b), time-domain (Koenig et al., 2002a), or time/frequency domain (Makeig et al., 2004) DPs are also essential. In the later stages of the multinational qEEG normative project, we will include these types of DPs. Table III contains a summary of EEG frequency domain DPs, which we now describe formally.

**Table III:**
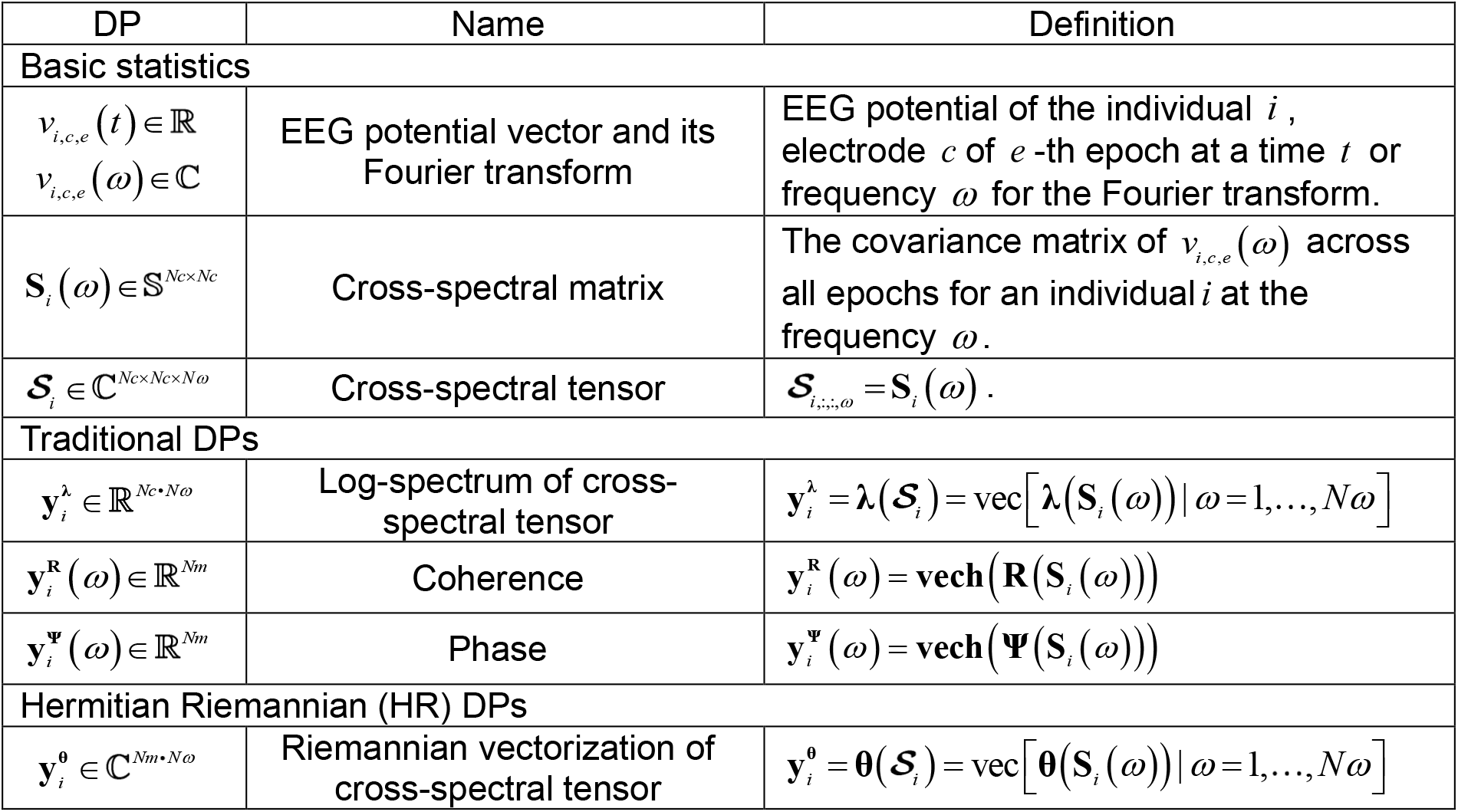
EEG frequency domain descriptive parameters (DPs)

| DP | Name | Definition |
| --- | --- | --- |
| **Basic statistics** |  |  |
| $v_{i,c,e}(t) \in \mathbb{R}$<br>$v_{i,c,e}(\omega) \in \mathbb{C}$ | EEG potential vector and its Fourier transform | EEG potential of the individual $i$, electrode $c$ of $e$-th epoch at a time $t$ or frequency $\omega$ for the Fourier transform. |
| $\mathbf{S}_i(\omega) \in \mathbb{S}^{N_c \times N_c}$ | Cross-spectral matrix | The covariance matrix of $v_{i,c,e}(\omega)$ across all epochs for an individual $i$ at the frequency $\omega$. |
| $\mathcal{S}_i \in \mathbb{C}^{N_c \times N_c \times N\omega}$ | Cross-spectral tensor | $\mathcal{S}_{i,\dots,\omega} = \mathbf{S}_i(\omega)$. |
| **Traditional DPs** |  |  |
| $\mathbf{y}_i^\lambda \in \mathbb{R}^{N_c \times N\omega}$ | Log-spectrum of cross-spectral tensor | $\mathbf{y}_i^\lambda = \boldsymbol{\lambda}(\mathcal{S}_i) = \text{vec}[\boldsymbol{\lambda}(\mathbf{S}_i(\omega)) \vert \omega = 1, \dots, N\omega]$ |
| $\mathbf{y}_i^{\mathbf{R}}(\omega) \in \mathbb{R}^{Nm}$ | Coherence | $\mathbf{y}_i^{\mathbf{R}}(\omega) = \text{vech}(\mathbf{R}(\mathbf{S}_i(\omega)))$ |
| $\mathbf{y}_i^{\Psi}(\omega) \in \mathbb{R}^{Nm}$ | Phase | $\mathbf{y}_i^{\Psi}(\omega) = \text{vech}(\boldsymbol{\Psi}(\mathbf{S}_i(\omega)))$ |
| **Hermitian Riemannian (HR) DPs** |  |  |
| $\mathbf{y}_i^{\theta} \in \mathbb{C}^{Nm \times N\omega}$ | Riemannian vectorization of cross-spectral tensor | $\mathbf{y}_i^{\theta} = \boldsymbol{\theta}(\mathcal{S}_i) = \text{vec}[\boldsymbol{\theta}(\mathbf{S}_i(\omega)) \vert \omega = 1, \dots, N\omega]$ |

The multi-channel EEG **v**_*i,e*_(*t*) is a vector time series recorded for the participant *i*, at the time *t*, for the e-th epoch (Figure 1-a). where, *i* = 1,…, *Ni* (*Ni* is the number of subjects), *e* = 1,…,*Ne* (*Ne* is the number of epochs which is an uninterrupted fixed-length sequence of time points). *t* = 1,…,*Nt* (*Nt* is the number of time points in each epoch). Note that *t* is an integer index. When referring to actual time (in seconds or milliseconds), *t* is multiplied by Δ*t*. This conversion (Δ*t* · *t*) allows physiological interpretation.

**Figure 1:**
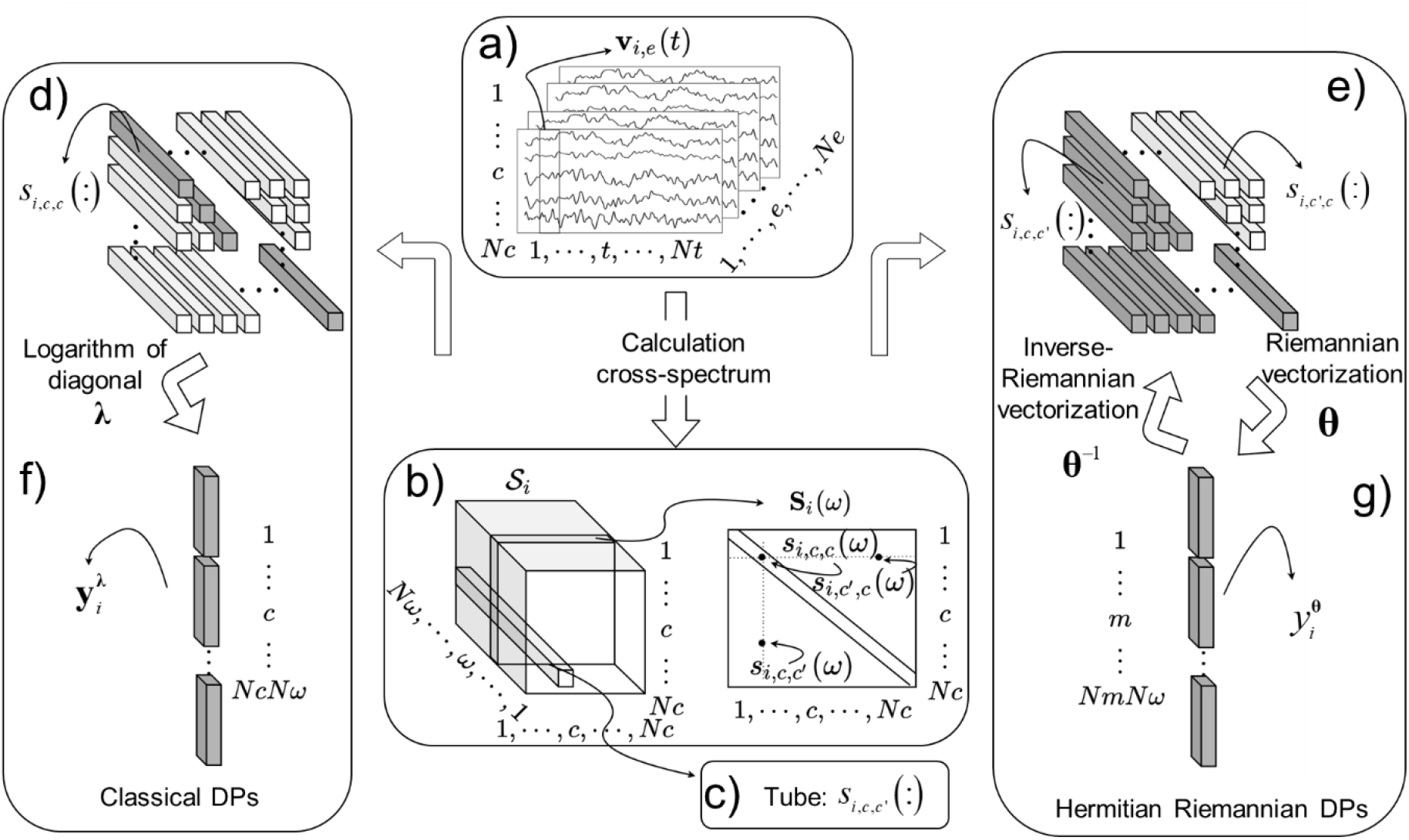
Traditional and Hermitian Riemannian EEG Frequency Domain Descriptive Parameters (DPs). a) Frequency domain DPs result from processing multi-channel EEG data *v_i,e_*(*t*) epochs (individual *i*, time *t*. e-th epoch).b) The cross-spectral matrices **S**_*i*_(*ω*) are the covariance matrices of the Fourier transform of the **v**_*i,e*_(*t*). These **S**_*i*_(*ω*) are the frontal slices of the tensor 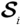; c) Besides columns and rows, the tensor has tubes *s_i,c,c′_*(:) which are the set of elements for two fixed channels and all frequencies. d) diagonal fibers. e) off-diagonal fibers. f)Vertically stacking the diagonal tubes’ logarithm generates the traditional DPs vector 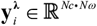 for fixed channels. g) Hermitian Riemannian DPs result from the half-vectorization of the matrix logarithm of the geometrically centered **S**_*i*_(*ω*), producing the vectors 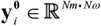. The inverse process is **θ**^-1^ which recovers 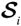.

The vector **v**_*i,e*_(*t*) has, as components, the set of scaler potentials *v_i,e c_*(*t*), measured with a common reference, for each channel *c*, where *c* = 1,…,*Nc*(*Nc* is the number of channels). To streamline our presentation, we indistinctively use the channel number of a particular montage or its channel label. An example is *v*_*i,e*,9_(*t*) which corresponds to the recording at the left occipital channel in the 10-20 system *v*_*i,e,O*1_(*t*).

Each EEG segment *v_i,e,c_*(*t*) is transformed to the frequency domain via the (discrete) Fourier Transform to yield the complex-valued coefficients *v_i,e,c_*(*ω*), where *ω*(*Nc* is the number of electrodes) and the symbol *ω* denotes frequency. Note that *ω* = 1,…,*Nω* (*Nω* is the number of the frequencies sampled). The frequency resolution multiped by Δ*ω* refers to actual frequency *ω*·Δ*ω* (in Hz.). Careful selection of epochs ensures that a) they are approximately stationary and b) lack long-range memory correlations in time. The Fourier coefficients are then asymptotically (as *Nt* → ∞) independent for each frequency. They are sampled from a Circular Multivariate Complex distribution 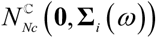 with mean **0** and population covariance **∑**_*i*_(*ω*) (different for each participant *i*). (Brillinger, 1981, chapter. 4.3). The set of **∑**_*i*_(*ω*); *ω*=1,…,*Nω* is the “cross-spectra” are our frequency domain DPs. These DPs encode linear and stationary properties of the EEG and are sufficient statistics cross-spectrum for second-order stationary. It is important to note that the **∑**_*i*_(*ω*) are not symmetric as in the real case but Hermitian.

Since the **∑**_*i*_(*ω*) are unobservable population quantities, we must use the maximum-likelihood estimate, the covariance matrix **S**_*i*_(*ω*) of the complex-valued coefficients of the Fourier transform, pooled over all epochs. We assemble these sample covariances **S**_*i*_(*ω*);*ω*=1,…,*Nω* into a cross-spectral tensor 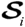 3-mode dimensional array with dimensions *Nc* ×*Nc* × *Nω* (Figure 1-b). In the parlance of tensors, fixing two of the dimensions and varying over the others produces “fibers”. Channels correspond to “column” and “row” fibers and frequencies to “tube” fibers. Also, in tensor terminology, “frontal slices” are just the usual cross-spectral matrices **S**_*i*_(*ω*). The tube fibers correspond to the “spectra” of time-series analysis. Let *s_i,c,c′_*(*ω*) denote the individual elements of **S**_*i*_(*ω*) (Figure 1-b), where *c* and *c′* denote channels *c,c′* = 1,…,*Nc*. Then we have two types of fibers *S_i,c,c_*(:) = {*s_i,c,c′_*(*ω*)|*ω* = 1,…,*Nω*} (Figure 1-c):

- When *c* = *c′* the fiber *s_i,c,c_* (:) sits on a diagonal element of the cross-spectral matrices over all frequencies, it is a real scalar tube known as the power-spectrum for the channel *c* (Figure 1-d). We shall refer to this type of tube or its element as “diagonal”.
- When *c* ≠ *c′* the fiber*s_i,c,c′_*(:) sits on an off-diagonal element of the cross-spectral matrices over all frequencies, this complex-valued tube is the cross-spectrum between channels *c* and *c′* (Figure 1-e). We refer to this type of tube or its elements as “off-diagonal”.
- Instead of the usual property *s_i,c,c′_*(*ω*) = *s_i,c′,c_*(*ω*) (valid only for real covariance matrices), we now have for the Hermitian case *s_i,c,c′_*(*ω*) = conj(*s_i,c′,c_*(*ω*)): symmetric elements are complex conjugates of each other.

For some calculations, using the complex value of an off-diagonal fiber *s_i,c,c′_*(*ω*) is inconvenient, and instead, we decompose these quantities into their real and imaginary parts (or absolute values and phases). We use the obvious notation, for example, Real(*s_i,c,c′_*(*ω*))-Imag(*s_i,c,c′_*(*ω*)) in the case of the cross-spectrum. Defined the coherence operator is 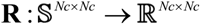, is the absolute magnitude of the complex correlation coefficient that,

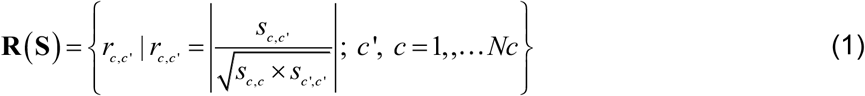

And the phase operator is defined as 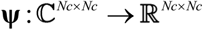,

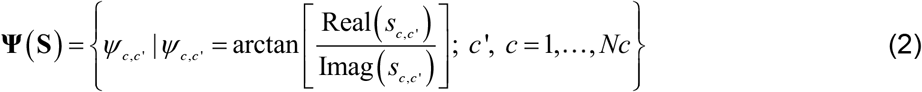

The earliest DPs used for qEEG norms were “Broad-Band” (BB) parameters that summarized the EEG spectrum by averaging over large segments of the frequency values (Duffy 1986; Burgess 1990). Instead of defining each *s_i,c,c_*(*ω*) as a DP for each *ω*, DPs aggregates measures: 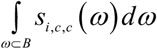. *B* is an a priori defined frequency band. BB DPs are still widely used, attempting to reduce the number of DPs. The bands try to reflect somewhat ambiguously defined clinically observed EEG rhythms. In practice, activity overlaps with several bands making BB DPs less sensitive.

In contrast, our group introduced “Narrow Band” (NB) or high-resolution resolution DPs (Szava, 1994). These are the complete set of *s_i,c,c_*(*m*) for each *ω*. We thus leverage the highest possible frequency resolution obtainable with the discrete Fourier transform. Henceforth the same symbol *ω* denotes either a discrete frequency or a band. We limit our attention to narrowband measures.

### 1.2 Traditional and Riemannian approaches to qEEG norms

DPs are highly age-dependent and are usually adjusted by the “z transform” (John, 1987) to provide age-independent measures of BDD. Formally, the z-score is based on the following model, defined for any type of frequency domain DP

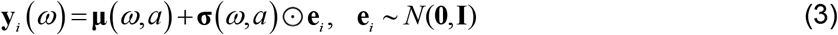

for any frequency *ω* and age *a*.

Thus the z score for any DP *y_i,c,c′_*(*ω*) of an individual, for electrode pairs (*c,c′*) and frequency *ω* is expressed in scalar form as:

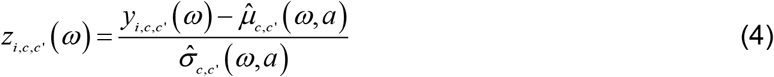

where *μ_c,c′_*(*ω, a*) and *σ_c,c′_*)(*ω, a*) are the frequency and age-dependent mean and SD, respectively. These regressions are known as “EEG development Equations” (Matoušek and Petersén, 1973b; John et al., 1977; Ahn et al., 1980) or “qEEG norms”. This concept of age-dependent norms also generalizes to include dependence on other independent variables such as sex, or as in the case of this paper site/country/study (batches). We shall call the new equations described in this paper “HarMNqEEG norm”.

The z-score is a probabilistic measure of the individual’s deviation from the normative population or BDD. Probabilistic statements about a z-score are most straightforward when the distribution of the DP for the normative sample is approximately Gaussian. Thus, it is convenient to transform DPs with a function that ensures this Gaussianity—a step carried out before calculating regression equations and z-scores. The most commonly used transformations are the logarithm of cross-spectrum diagonal, 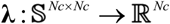 of 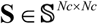,

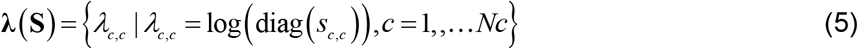

and its application on the cross-spectral tensor 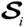 produce “traditional DP”, log-spectral vector 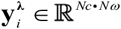 (Figure 1-f),

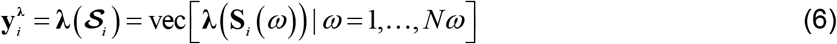

Our previous work created norms for the log-spectrum 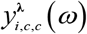 and introduced the term “Developmental surfaces” (Szava et al., 1994) for the Narrow Band (NB) developmental equations since both mean and SDs are bivariate functions of frequency and age. Developmental Surfaces describe EEG scalp channels and sources (Bosch-Bayard et al., 2001). These norms are now extensively used, with both the data and code now being open-source(Bosch-Bayard et al., 2020; Valdes-Sosa et al., 2021). These are the “traditional norms”.

Despite their widespread use, these traditional norms have several shortcomings:

i. The Developmental Surfaces are limited to the 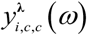 and have not dealt with the cross-spectra **S**_*i*_(*ω*)—thus sidelining functional connectivity measures. To our knowledge, there has been no attempt to create developmental surfaces for the complete cross-spectral tensor 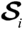.
ii. The log-transformations applied to the spectrum *s_i,c,c_*(*m*) are traditional univariate transformations that fail to ensure that all cross-spectral DPs are multivariate Gaussian. Empirical multivariate transformations can achieve this objective (Biscay Lirio et al., 1989), but they are time-consuming and not natural.
iii. Importantly the frontal slices of the cross-spectral tensor 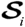, namely the cross-spectral matrices **S**_*i*_(*ω*) are Hermitian matrix positive-defined (HPD) matrices. They do not belong to a Euclidean space but rather live in a Hermitian manifold 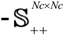 (Deza and Deza, 2013). The curved nature of this manifold precludes the use of simple Euclidean distances to quantify distances between cross-spectra, impeding both the calculation of norms and measuring distances of individuals to those norms.

Riemannian Geometry allows novel multivariate cross-spectral norms to solve the “traditional norms” three mentioned difficulties.(Deza and Deza, 2013) defined, “Hermitian Geometry studies complex manifolds equipped with Hermitian metric, it is a complex analog of Riemannian Geometry” Consequentially, we are working with a complex analog of the usual real-valued Riemannian Geometry typically used in BCI applications (Barachant et al., 2012). Riemannian Geometry is now finding wide application in Brain-Computer-Interface studies (Barachant et al., 2013). Sabagh Engelmann used real-valued Symmetric Positive-definite geometry to define brain age. Here we invert the dependent and independent variables and Hermitian geometry to obtain “Hermitian Riemannian norms “ (HR norms). For this, we first need to obtain Hermitian Riemannian DPs (HR DPs).

We define HR DPs by leveraging a consequence of Hermitian Geometric theory that provided a natural way to transform **S**_*i*_(*ω*) from its original curved HPD manifold to tangent space which lives in a flat Euclidean space where Gaussianity holds. We achieve this dual objective by applying the “Riemannian vectorization” operator (Pennec, 1999, 2004; Sabbagh et al., 2019). The Riemannian vectorization operator is defined as **θ**, 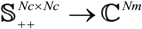,.

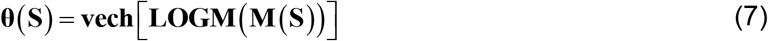

where, 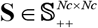. The operator **M**, **LOGM** and **vech** defined in Table II. The operation **M** is the Riemannian equivalent of the centering operation from the usual Euclidean multivariate statistics, which places the origin of the data to zero. Here the centering is to the identity matrix. We thus center {**S**_*i*_(*ω*)|*i* = 1,…, *Ni*} using 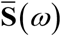 at frequency *ω*, the geometric mean (which minimizes the sum of square Riemannian distances of the set {**S**_*i*_(*ω*)|*i* = 1,…, *Ni*} at the common vector tangent space 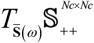) (Pennec, 2004; Bhatia and Holbrook, 2006).

The application of Riemannian vectorization of cross-spectral tensor produce 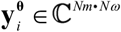 (Figure 1-g),

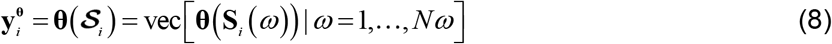

This transformation guarantees that the 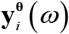 more closely satisfy the Multivariate Gaussianity assumption (Pennec, 2004). Note that we scale the off-diagonal 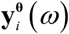 by 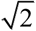 preserving the equality of norms 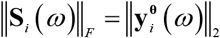.

The inverse operator of **θ** is 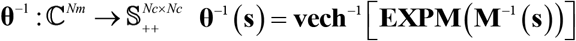 which returns the vector to the HPD manifold.

We have defined two types of DP vectors, traditional 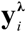 and Hermitian Riemannian 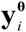. For ease of interpretation, when referring to elements of these vectors, instead of linear indices, we indicate the row and column channel indices (or labels) of a matrix with the dimensions as the cross-spectrum.

## 2 The harmonized multinational qEEG project

### 2.1 Data acquisition

The HarMNqEEG collaboration is creating multinational norms for the cross-spectral tensors 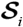. It has two main components:

1. Preparation of the data by the collaborators at each site sharing information (Figure 2-b)
2. Processing of the shared data to achieve harmonized norms (Figure 2-a).

**Figure 2:**
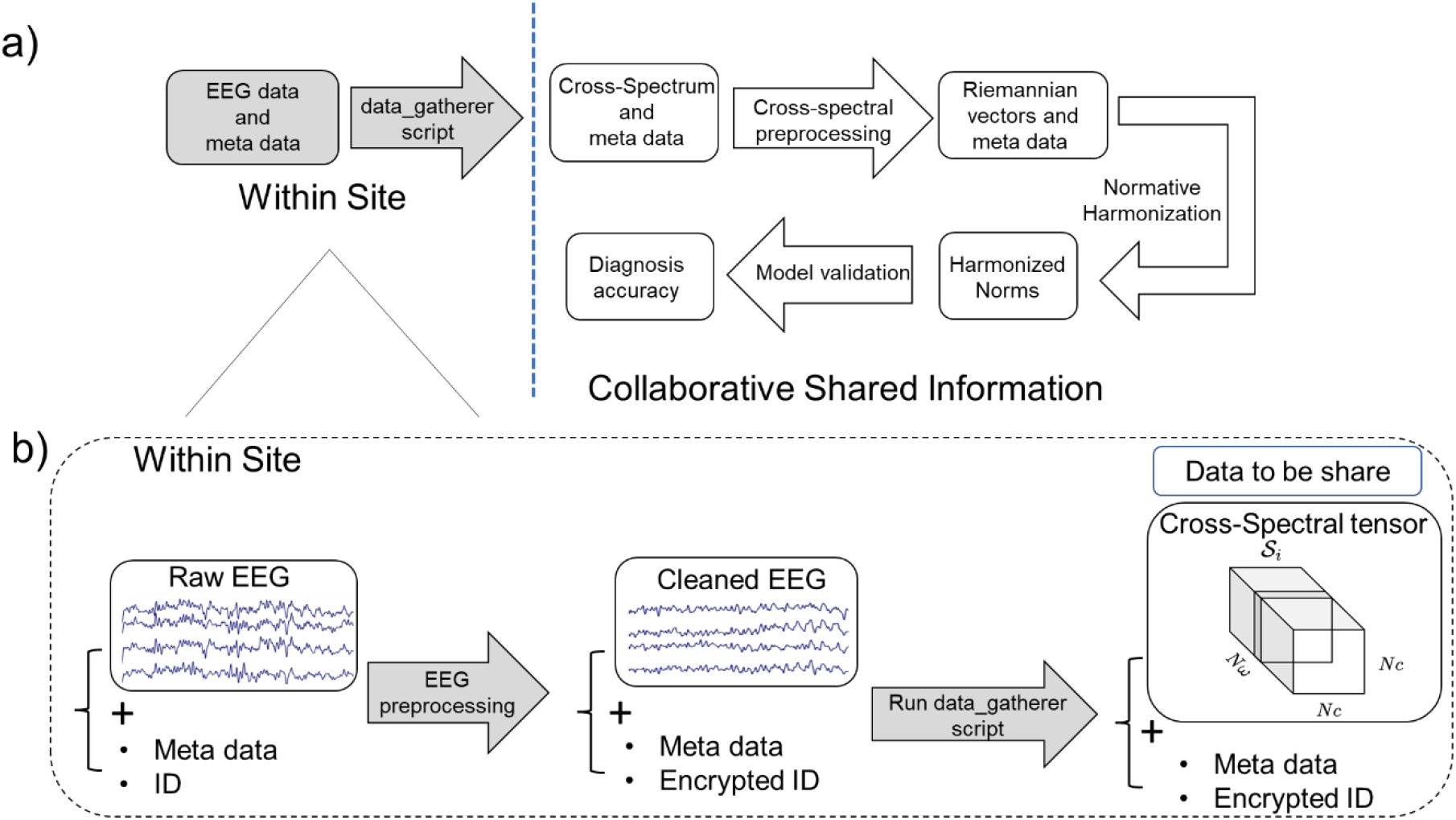
Dataflow of the collaboration for the Harmonized qEEG norms. a) General overview. Each collaborative site collected resting-state EEGs from healthy control subjects in the eyes closed condition (obtained from a normative study). After within-site quality control and artifact rejection, the MATLAB “data_gatherer” script was used to obtain the EEG cross-spectrum for each participant, which, together with anonymized meta-data, constituted a “sample”. These samples were shared with the central processing site, and after further calculations, yielded qEEG Descriptive Parameters (DPs), traditional and HR. In turn, these DPs were used to construct the harmonized qEEG norms. An independent data set of healthy subjects and those with Brain Developmental Disorders allowed the comparison of the diagnostic accuracy of different DPs. Boxes shaded in gray indicate data and process private to each collaborative site; b) Further datils of within-site processing. Each site carried out initial quality control of raw EEG recordings and metadata (sample), using procedures for artifact rejection. Samples with clean EEG and encrypted ID were processed with the MATLAB script “data_gatherer”, which computes the cross-spectrum and bundles it with the encrypted meta for sharing.

We now summarized the descriptive parameters and workflow to process them, explaining details separately in the subsequent sub-sections.

The project was launched by an open international call for participation via the Global Brain Consortium (https://globalbrainconsortium.org/; https://3design.github.io/GlobalBrainConsortium.org/project-norms.html). The collaborative model only shared processed data, EEG cross-spectral tensors 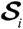 and anonymized metadata. This data was obtained using the within-site software data gatherer MATLAB program, as shown in Figure 2-b.

A prerequisite for any site to join the study was to have ethical approval by the corresponding authorities to share processed data (EEG cross-spectra tensors 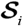) and anonymized metadata. The inclusion/exclusion criteria for the normal subjects have been described respectively in the reference, the last column of Table A-II in Appendix A-II. These criteria are sufficiently and equally stringent to guarantee a sample of functionally healthy subjects.

Each site submitted a “batch” of samples. A sample contains a cross-spectral tensor 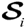 and the metadata: sex, *s_i_* age *a_i_*, and batch *b_i_* Here *i* tags each individual. See the dimensions of 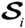 in the last block of Figure 2-b.

To be accepted into the study, the batch had to fulfill the following requirements:

1. It had to be part of a normative study or control group with explicit inclusion and exclusion criteria (see below).
2. The 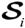 had to be obtained with the MATLAB script (data_gatherer) from at least one minute of artifact-free, eyes closed, quasi-stationary, resting-state EEG epochs **V**_*i,e*_(*t*).
3. Finally, the batch had to pass numerical quality control with partial or total rejection of the batch or samples being possible (detail see section 4).

We explain some design decisions of this project, which were necessary given the significant difference in recording protocols for the different batches. We homogenized a minimalistic set of specifications applicable to all sites and devices:

- Recordings were from the 19 channels *Nc* = 19 of the 10/20 International Electrodes Positioning System: Fp1, Fp2, F3, F4, C3, C4, P3, P4, O1, O2, F7, F8, T3/T7, T4/T8, T5/P7, T6/P8, Fz, Cz, Pz) ([Standards and Best Practices organization for open and FAIR neuroscience | INCF](https://www.incf.org/)).
- The amplifiers included a minimal frequency response range from 0.5 to 35 Hz.
- On-site artifact rejection did not include the use of ICA techniques.
- Each EEG was organized as a sequence of artifact-free 2. 56 seconds. This format allows a frequency resolution of 0.39 Hz. Frequency analysis of the EEG (via the discrete Fourier transform) was limited to the range 1.17 to 19.14 Hz, obtained by sampling in Fourier coefficients for *ω*∈(3,…,49) a total of 47 frequency points with frequency resolution Δ*ω* = 0.39. The gatherer code guaranteed this frequency resolution. If a higher frequency resolution is detected, it is downsampled to match the standard one. IF a lower and lower frequency was detected, the spectra were calculated using the original sampling frequency and then interpolated using a Piecewise Cubic Hermite Interpolating Polynomial (PCHIP) (Fritsch and Carlson, 2006; Kahaner et al., 1989).
- The scalp EEG cross-spectrum was calculated using Bartlett’s method (Møller, 1986) by averaging the periodograms of more than 20 consecutive and non-overlapping epochs of 512 samples, regardless of possible discontinuity in the recovered records.
- Finally, the cross-spectrum was restricted to the standard frequency range of 1.17 Hz to 19.15 Hz.
- Following the principles of open science but respecting the researchers’ rights to retain control of their raw data, all these functionalities were encapsulated in a script programmed in MATLAB and distributed among the researchers in each recording site. (Github location of the script: https://github.com/CCC-members/BC-V_group_stat/blob/master/data_gatherer.m). Each site ran the script on their data and only shared the processing results without sharing their raw data. Thus, they only sent shared the EEG spectra, basic subjects’ information as an anonymized code, age, and sex, as well as technical parameters like recording conditions, montage, recording reference, EEG device used, laboratory, and country.

We define the term “batch” as a specific combination of country and device and year of recordings (Table A-III, Appendix A-III). Three different definitions of batches reflect different hypotheses, namely:

a. (Country) Batches are the countries from which data comes.
b. (Device) Batches are the specific type of equipment from which data come.
c. (Study) Batches are the specific projects which generate a data set, a combination of country, device, and year of study.

We test whether batches, defined in these different ways, need to be accounted for when calculating qEEG normative equations. If there are systematic differences between the normative equations of each batch, one must add a batch-specific additive correction to these equations.

The multinational call for the multinational qEEG norms and subsequent batch selection produced 1792 samples. After quality control, samples diminished to 1564, with 783 females and 781 males. A further breakdown of the samples by country, device, study, and age range is in Appendix A-II.

Figure 3 gives an overview of the age range of the samples is 5-97 years. The Age distribution of the Multinational qEEG norms sample is skewed towards younger ages, with relatively fewer samples older than 65 years. In addition, there is an almost balanced gender distribution for all the samples.

**Figure 3:**
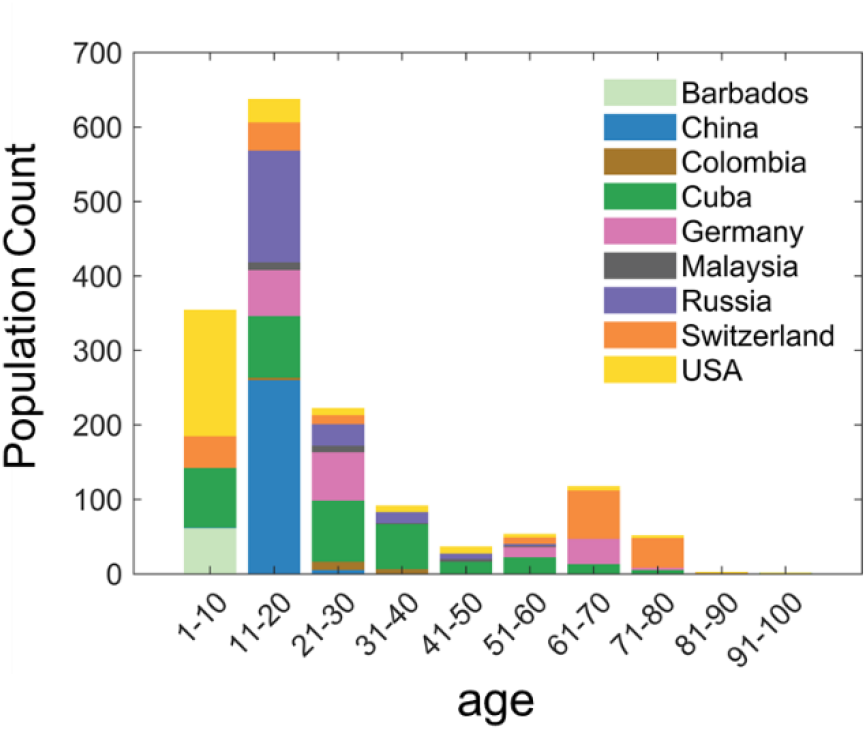
The dataset was collected from 9 countries, including Barbados, China, Colombia, Cuba, Germany, Malaysia, Russia, Switzerland, and the USA. The ages of samples for multinational EEG norms span the whole life span (5-95 years). Sampling is skewed towards younger participants, reflecting sampling of normative projects involved.

### 2.2 Preprocessing procedures and transformation of DPs towards Gaussianity

To be able to pool all cross-spectrum under the same framework, at the same time controlling for irrelevant nuisance variables, we implemented the following preprocessing steps:

#### 2.2.1 Average Reference

We center all cross-spectrum matrices **S**_*i*_(*ω*) from their original recording montages to the Average Reference montage (Hu et al., 2019):

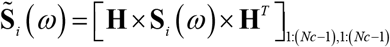

with the operator **H** = **I**_*Nc*_ –**1**_*Nc*_/*Nc*

The final electrode (Pz) was eliminated from 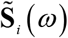 to avoid the trivial singularity of matrices.

#### 2.2.2 Regularization of symmetric semi-positive defined matrices

We regularize sample cross-spectrum matrices to ensure them to be of full rank. For this, we use the Maximum likelihood shrinkage factor described in (Schneider-Luftman and Walden, 2016). The 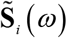 are Hermitian matrices, and frequently are rank reduced. To guarantee subsequent Riemannian operations (especially the matrix logarithm operator), we regularize 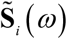 to ensure positive definiteness. Regularization was achieved with the method of (Schneider-Luftman and Walden, 2016):

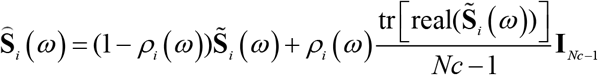

Where the shrinkage coefficient *ρ_i_*(*ω*)∈(0,1) is obtained by minimizing the Hilbert-Schmidt loss function, which results in:

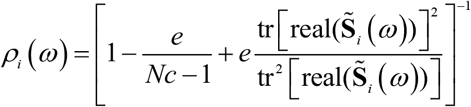

Here, 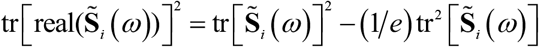.

#### 2.2.3 Global-scale correction

Though two EEG recordings may show a similar appearance, they may differ significantly in overall amplitude. This observation is modeled as a general gain factor is randomly varying for similar EEG data. These sources of variance arise from different EEG devices, recording conditions, amplifiers, and other subject characteristics (skull thickness, hair thickness, skin impedance, and other non-physiological factors). A solution to this nuisance source of variability is to rescale the cross-spectra. A random general scale factor (GSF) as described in J. L. Hernández et al., (1994). These authors showed that the maximum likelihood estimate of the GSF for an individual is the harmonic average of all log power values across all derivations

In more detail, suppose for an individual *i*, the EEG potential recorded at the electrode *c* is *v_i,e,c_*(*t*) = *γ_i_β_i,e,c_*(*t*), the global scale factor *γ_i_* is independent of time epoch and electrode. Taking the Eigen decomposition 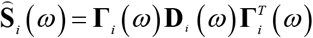, where **Γ**_*i*_(*m*) is the eigenvector matrix and **D**_*i*_(*ω*) is the diagonal matrix of eigenvalues. Rescaling the **Ŝ**_*i*_(*ω*) we have:

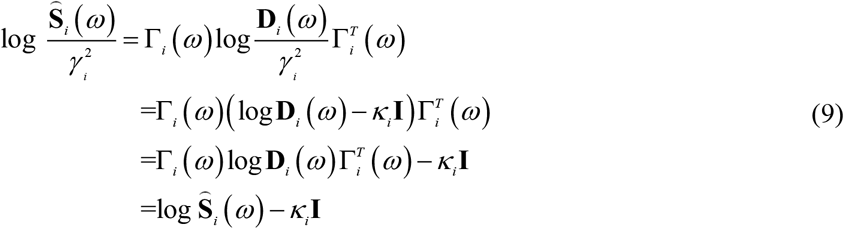

Thus, the GSF affects the diagonal of the cross-spectrum (the log spectra). The estimator of *κ_i_* is the geometric mean of log spectrum, *κ_i_* = 2log(*γ_i_*),

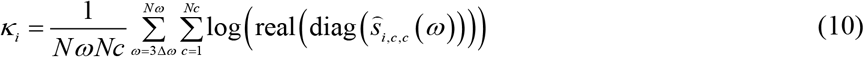

Then the GSF-corrected cross-spectrum can be represented as,

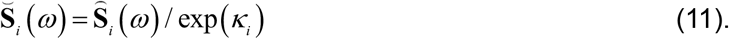

#### 2.2.4 The logarithm of spectra and Riemannian vectorization of cross-spectra The final step is to obtain gaussianity DPs

For the traditional log-spectrum DPs, get 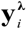 with diagonal logarithm operator as in the equation (6). To obtain 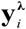., we only apply steps 2.2.1 and 2.2.3 for preprocessing.

For the HR DPs, we transform **S**_*i*_(*ω*) to the Euclidean tangent space, 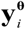 employing the Riemannian vectorization operator as in the equation (8).

## 3 Construction of multinational harmonized qEEG norms

### 3.1 Possible normative models

Here, for the HarMNqEEG modeling, we work on the two types of DPs, 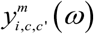, *m* = ***λ*** or **θ**, where *m* are types of DPs shown in Table III, which we assume satisfy the gaussian distribution. In Table IV, we summarize normative equations and related z-scores.

**Table IV:**
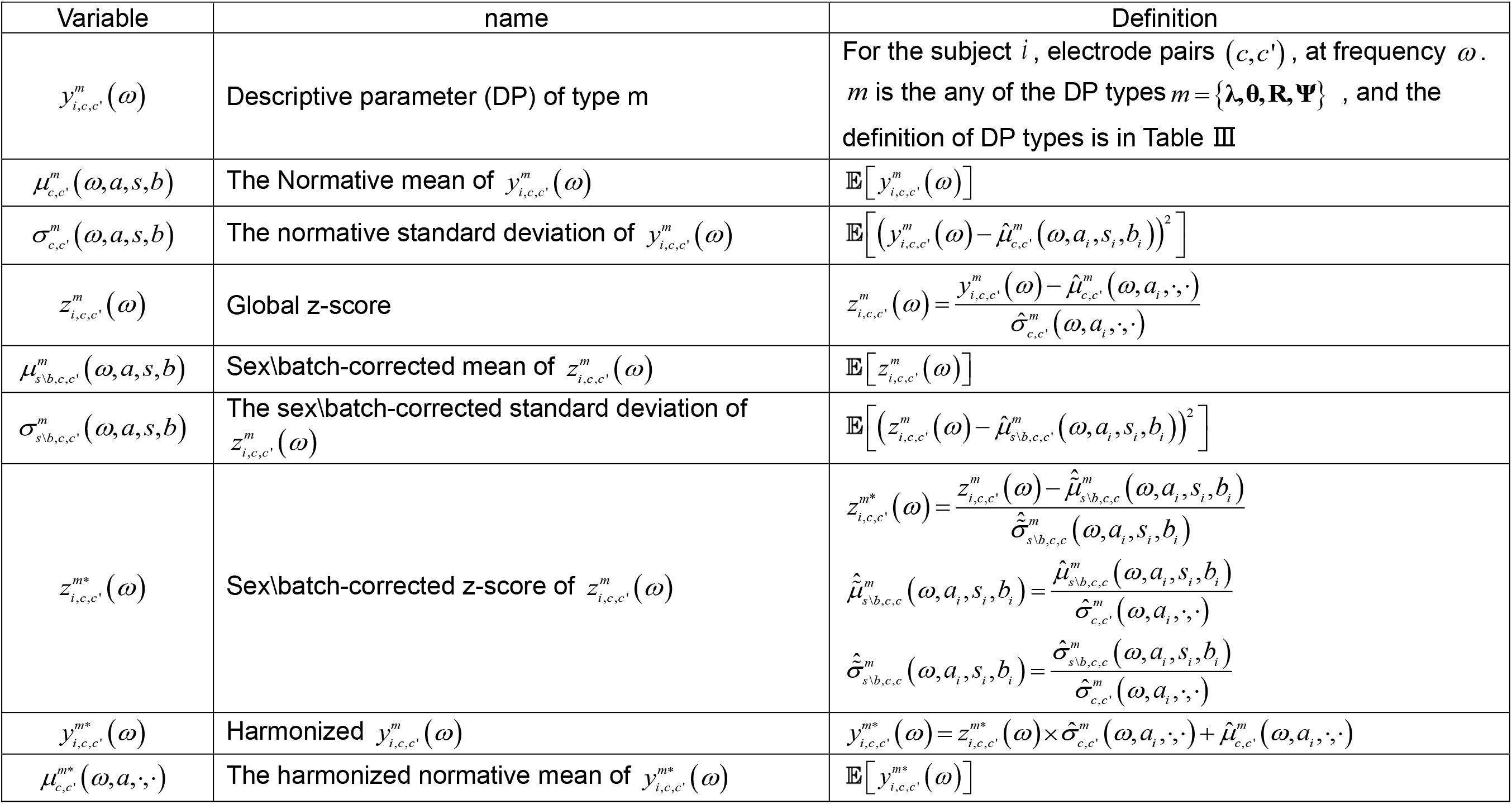
Normative equations and related z-scores

Each 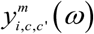 can be expressed as a general linear model (GLM):

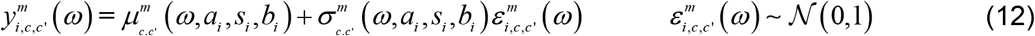

where 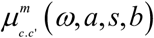 and 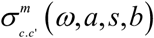 are the population means and SD, respectively. Henceforth, the value of a variable for an individual is denoted with a subscript, e.g. *a_i_*. Also, the symbol instead of a variable indicates that it is pooled over all individuals. These conventions for indices are summarized in Table A-1.

Unfortunately, the general HarMNqEEG model (12) is very “data greedy”, being too complex. For that reason, we explore more parsimonious additive models, each depending on a smaller subset of variables. Instead of the general 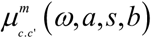, we consider the additive models described in Table V. The most trivial model assumes that 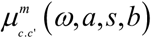 is a constant 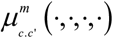. Given our prior work, we chose as fixed effects only frequency and age, with possible models 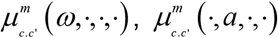, and 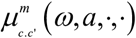. We then consider additional constant additive random effects that depend on batch: 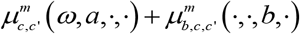, sex 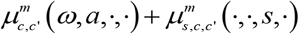 or batch and sex 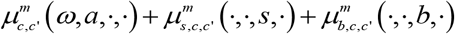. We finally look at the last level of complexity for the population mean: the additive random effects (for example, the random effects is a batch effect), instead of being constant, are now functions of frequency and age, 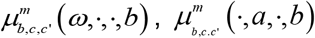 and 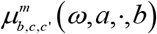.

**Table V:**
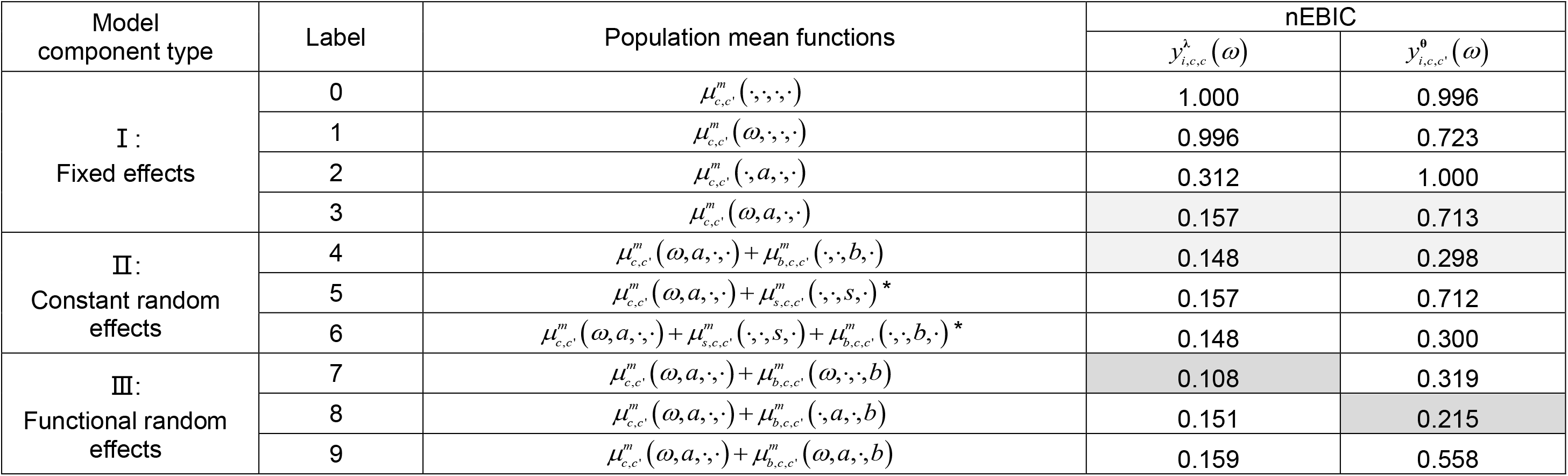
Extended Bayesian Information Criterion of models for DPs

Similarly to the mean, we model the SD shown in Table VI with fixed effects, for example, 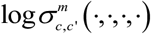 or 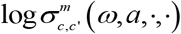. Additive batch random effects considered were either constant 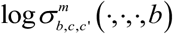 or functions 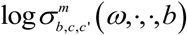 and 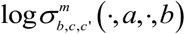 dependent on frequency and age, respectively.

**Table VI:**
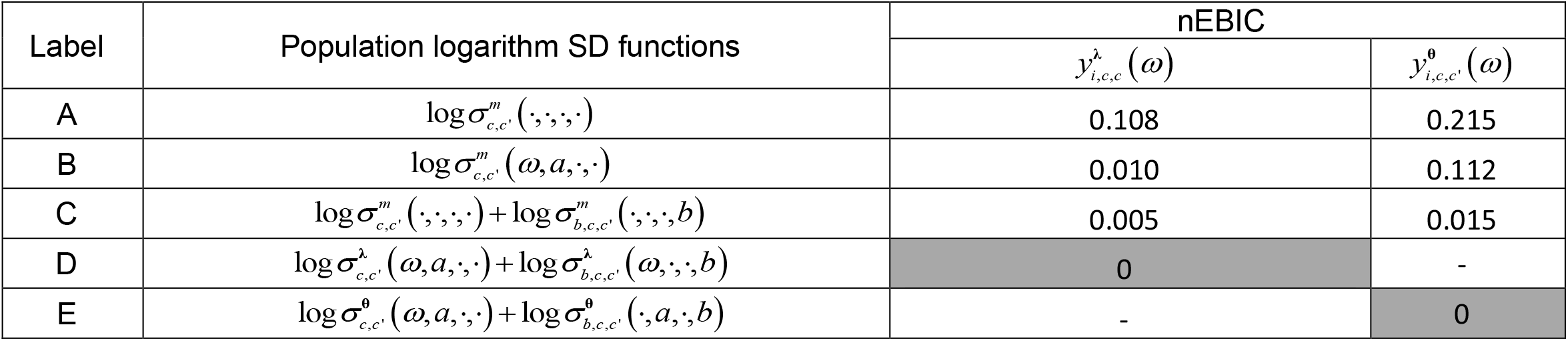
Models for the population logarithm SD

When fitting these models, we follow a sequential strategy, fitting first the fixed effects and then, based on the residuals or z score of this fit, we fit the random effects if required. A concrete example of this is deferred to section 3.3. All estimates of the mean function and SD functions (including fixed and random effects) are obtained with the Nadaraya-Watson (NW) kernel regression (Nadaraya, 1964). This nonparametric smoothing method depends on the bandwidth hyper-parameter, which controls smoothing. The more smoothed the data, the less complex the model. The complexity of the model is reflected in the “equivalent” degrees of freedom (Fisher, 1922). Special consideration was given to estimating the population variance as it does not have a Gaussian distribution as assumed by NW regression. Instead, for the variance, we used the modified NW regression to estimate the variance function for heavy-tailed innovation (Chen et al.,2009).

To stabilize sub-sample (batch or sex) estimators, we fixed their smooths to the same bandwidth as the global smooth, thus using the bandwidths obtained with all the data for the smaller samples. The shared bandwidth chosen for each model is optimal for a given DP set 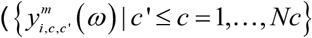 for model 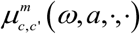 and 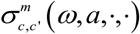. In this paper, the bandwidths for the mean smooth were 0.4 and 048 for frequency and age, respectively. The corresponding bandwidths for the variance smooth are 0.6 and 0.72.

Due to the large amount of computation needed for our models, instead of applying ordinary NW regression, we used an in-house procedure Fast multivariate kernel regression with nufft (nufft-mkreg). Our nufft-mkreg is a nonparametric multiple multivariate kernel regression based on a fast binning algorithm (Wand, 1994) that executes in *o*(*m*log*m*) operations instead of *o*(*n*^2^), and usually, the gridding points *m* are smaller than the number of variables *n*. About the nufft-mkreg in-house code, we mention that it includes the capability for complex-valued DPs regression (The nufft-mkreg paper is under preparing). We, therefore, use the FBR algorithm to estimate real and imaginary smooths of DPs simultaneously with a common variance. To our knowledge, this is the first instance of complex-valued qEEG harmonized norms. We also used a fast bandwidth selection for multiple multivariate smooth. The speed-up was possible by the use of the randomized generalized-cross-validation (rgcv) to approximate the best range of degrees of freedom (df) (Girard, 1989). Subsequently, we fine-tune the df with the more accurate (but more time-consuming) method of Turlach and Wand (1996).

The COMBAT model (Johnson et al., 2007) is our model 4-C for 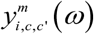, GAMLSS (Rigby and Stasinopoulos, 2005) is our model 7-D and 8-E. For Gaussian noise, both these models are just particular instances of our formulation.

### 3.2 Normative model selection

As mentioned in 3.1, the equation (12) can have different specifications. To find the optimal HarMNqEEG model, we minimize information-theoretic measures that are a tradeoff between model fit and model complexity. For standard statistical scenarios, examples of these criteria are Akaike’s information criterion (AIC) (Akaike, 1973), cross-validation (CV)(Stone, 1974), generalized cross-validation (GCV)(Craven and Wahba, 1978). The Bayes information criterion (BIC) (Schwarz, 1978) is of particular interest due to its good properties and adopted in this paper.

Unfortunately, in our data, the number of DPs is large compared to that of samples. This excess of variables is the “small n large p” problem, common in bioinformatics and neuroimaging, making most model comparison criteria (including BIC) perform poorly. BIC performs too “liberally,” usually picking excessively complex models. Chen and Chen (2008) propose a correction for BIC in the “small n large p” scenario. They diagnosed that the uniform prior on the model space is the cause of BIC’s liberality in the small-n-large-p situation. They correct this problem with a family of Extended Bayes information criteria (EBIC)(Chen and Chen, 2012). The EBIC value for a model is,

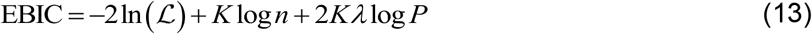

Where, *K* is the model’s degree of freedom, 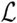 is the model likelihood. *P* = *O*(*n^k^*), and *λ* > 1 – 1/2*k*. We set *λ* = 0.3 based on independent simulations and to remove the scalar, we normalized the EBIC (nEBIC) values to [0,1] of all possible models.

Table V and Table VI show the nEBIC for the sequence of tested models, containing results for both 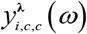 and 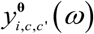. We thus ranked models based on the nEBIC criterion and selected the optimal HarMNqEEG model with the lowest nEBIC.

We first examined the optimal structure for the mean function in the equation, assuming a homoscedastic variance model that 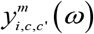 was a constant. The trivial model 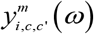, labeled as “0” in Table V, assumes no dependence on any covariates and is only included as a baseline null model. Models with frequency and age fixed effects, labeled 1-3 in Table V, were checked next. Note that a model depending on frequency alone did not lower the nEBIC noticeably. On the other hand, modeling age substantially decreased the criterion. The combination of age and frequency achieved the minimum (marked with light gray in the level I). Thus 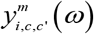 was kept as a fixed effect for all subsequent models tested.

Next, we turned attention to the models that add constant random effects related to sex or batch, labeled 4 to 6. You can see that the best model depends only on the batch effect for 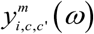. Surprisingly, including gender as a covariate does not improve the criterion. Gender also deteriorates model performance when batch effects are included (models with an asterisk). We, therefore, discounted gender from further exploration. Finally, we focused on functional random additive effects for the homoscedastic model family where batch interacts with age and frequency (models 7 to 8). For 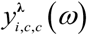, the optimal model includes both variables 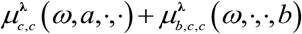. Contrary to our initial expectations, the random functional effect chosen for 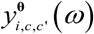 was 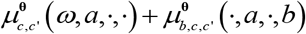, which depended on age but not frequency.

At this stage, an inspection of the models’ residuals suggested that the homoscedastic assumption is not realistic. We, therefore, searched for the best model for the SD, as we show in Table VI. All heteroscedastic models improved substantially on the homoscedastic one (model 7-A and 8-A for 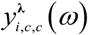 and 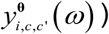. Note that we assumed that the model for the SD should have a similar form as for the mean. This choice is plausible, reducing the number of models to examine. The model with overall lower nEBIC is 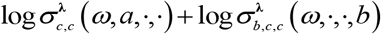 for 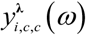, and 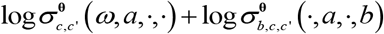 for 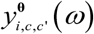.

To summarize, the optimal normative model selected for 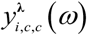 was,

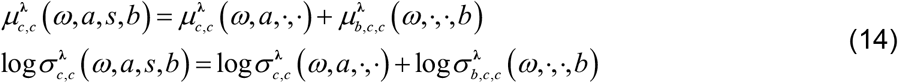

The optimal model for 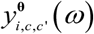 was:

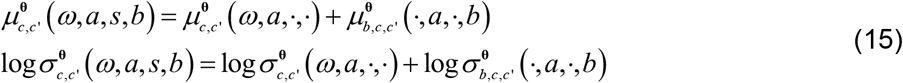

### 3.3 The HarMNqEEG norms

The HarMNqEEG norms are data and procedures that calculate global and harmonized z-scores. It does this by using the model (14) for the 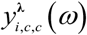 and the model(15) for the 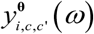. The norms thus contain information for calculating the global developmental surface of means and SDs, with additional batch corrections for the models described in section 3.2.

For the 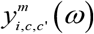, *m* = ***λ*** or **θ**, the basic model is:

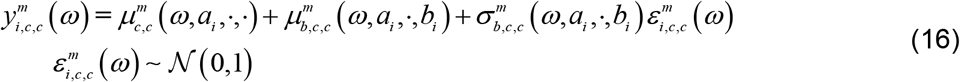

For computational expediency, to carry out sequential z-scores for the fixed and random effects, we rescale the random effects mean and the SD by dividing them with the fixed effect SD that 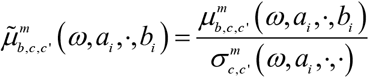 and 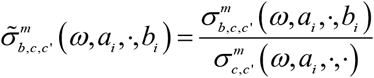. The modified model now reads:

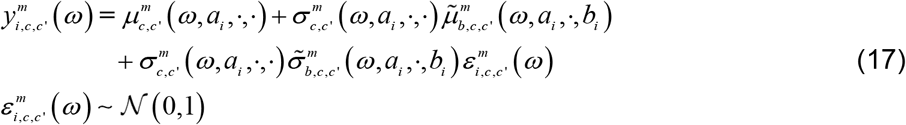

The steps to obtain the z-scores for 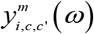 are: 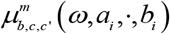 and 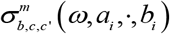

1. We first ignore the possible batch effects and fit global estimates for 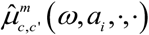 and 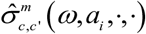
2. We then calculate the “batch-free” z-score value as

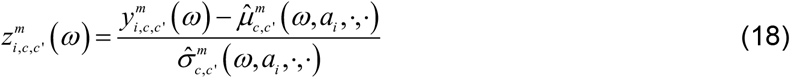
3. We obtain the batch-specific mean estimators 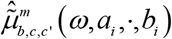 and SD estimators 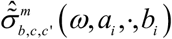. The batch-harmonized z-score is:

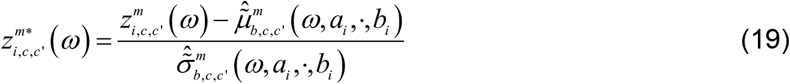

The HarMNqEEG norms 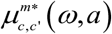 for 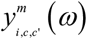 are obtained by smoothing the batch harmonized 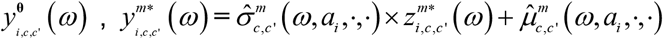.

Note that since we removed all random effects, we can omit the “.” symbols and use the notation 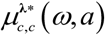 as in section 1.2.

To see how well the HR norms approximate the classical norms, we first calculate the “surrogate” cross-spectral norms:

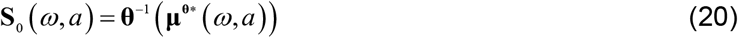

We then obtain surrogates of the classical norms from this cross-spectral norm by applying the operators **λ**, **R**, and **Ψ** on **S**_0_(*ω, a*). The process is summarized in Table VII.

**Table VII:**
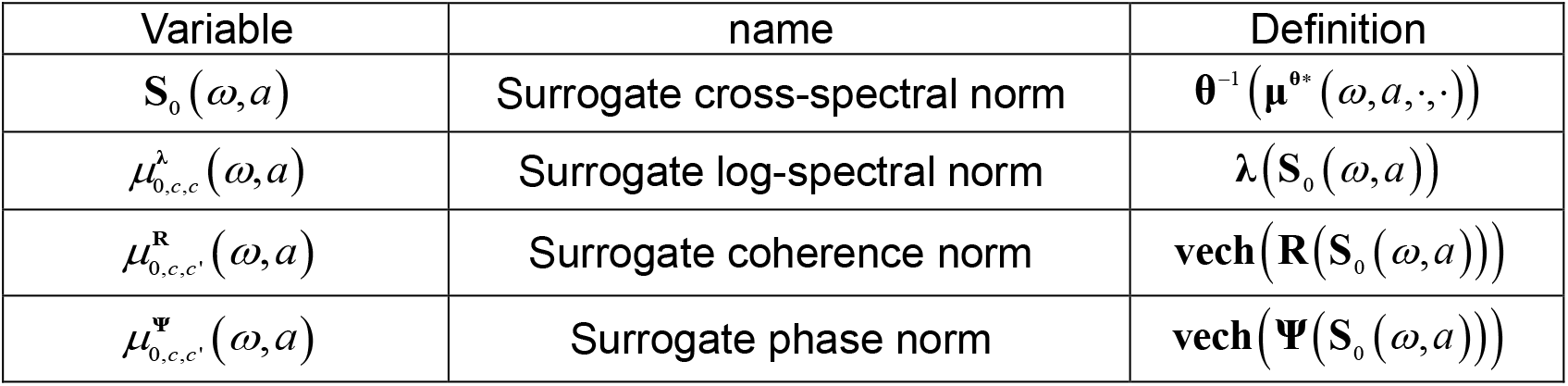
Surrogate traditional normative equations

We first show in Figure 4 examples of harmonized developmental surfaces for the diagonal elements Fp1, O1, and O2. Parts a)-to b) of that figure corresponds to the measures 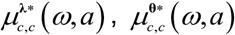, and 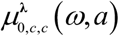 that it the traditional DPs, the HR DPs, and the surrogate Riemannian log-spectrum. To be noted is the similarity of Figure 4-a (the detail in Figure S2) to the developmental surfaces reported in Szava et al. (1994). Our current, more extensive, multinational dataset produces very similar results to the previous, smaller, single-country study. Figure 4-b shows the surfaces for 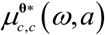 (the detail in Figure S3), which are quite different from those in Figure 4-a 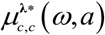. These differences are not surprising considering the highly nonlinear nature of the transformations involved in passing to the manifold tangent space—involving centering with the geometric mean covariances and a matrix-logarithmic transformation. The consistency of the norm construction procedure is illustrated by the concordance between the traditional log-spectra 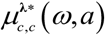 (Figure 4-a).and the surrogate log-spectra 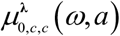 (Figure4-c), confirming the results of Szava et al. (1994). Thus, when analyzing traditional log-spectral measures, 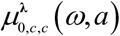 we base our subsequent analyses on **S**_0_ (*ω, a*).

**Figure 4:**
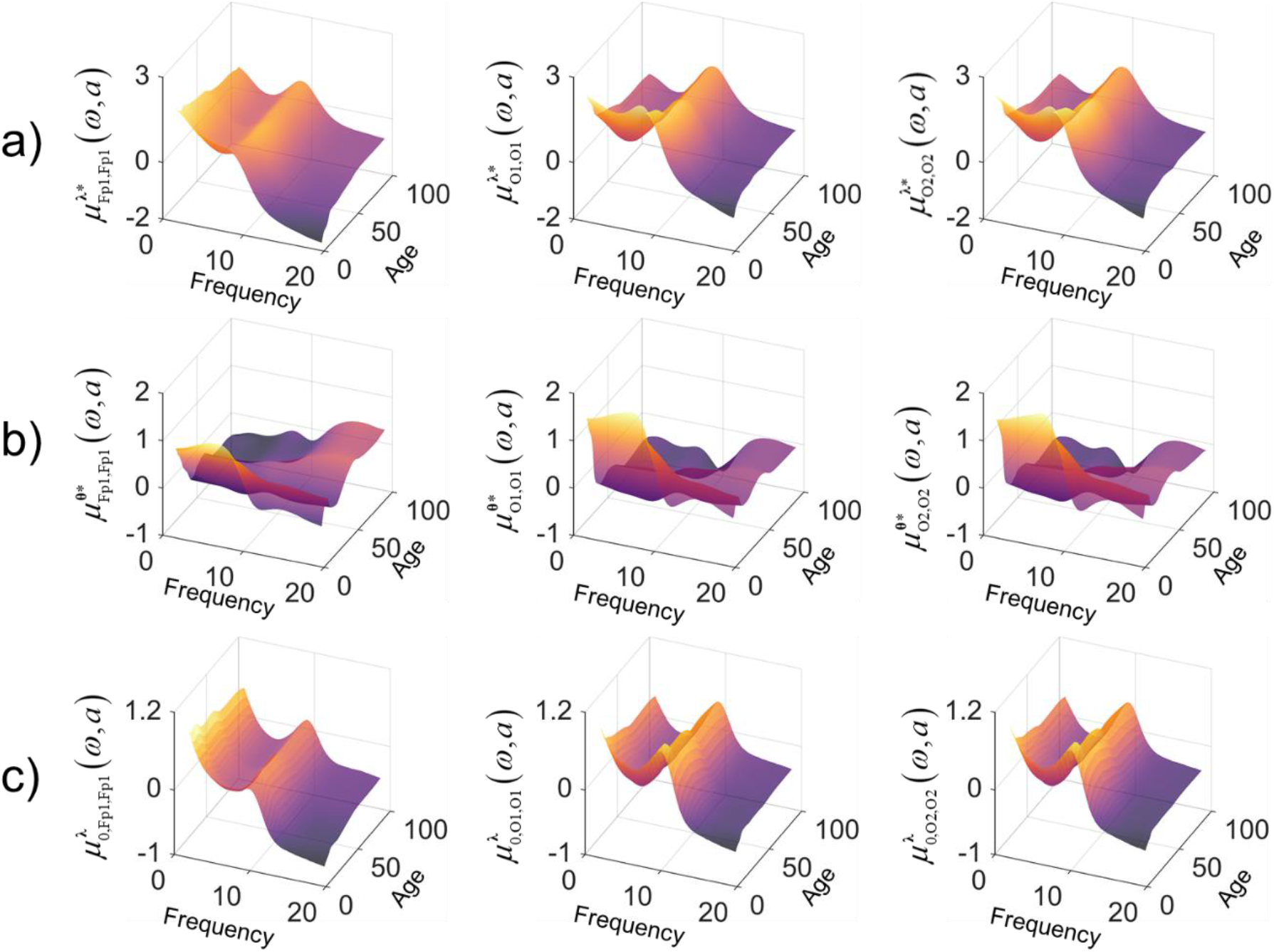
Examples of harmonized normative means for channel pairs *c* = *c′* as a function of frequency and age. Normative means (Developmental surfaces) with examples *c* = *c′* = Fp1 or O1 or O2 .a) Traditional log-spectra- 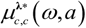; b) The HR norm 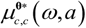; c) Surrogate log-spectrum norm 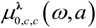, reconstructed from the normative mean of **S**_0_ (*ω, a*)

Figure 5 provides a more detailed view of the surrogate log-spectral developmental surface depicted in Figure 4-c. Fig. 5-a) plots surface cuts of 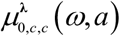 for all frequencies at fixed younger ages (5, 15, 25, and 40). Fig. 5-b shows similar plots for older ages (45, 60, 80, and 95). Orthogonal cuts of 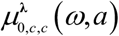 are shown in Figure 5-c, documenting changes with age at a few specific frequencies (2Hz, 8Hz, 10Hz, 15Hz)—the vertical lines on this figure mark 16 and 50 years.

**Figure 5:**
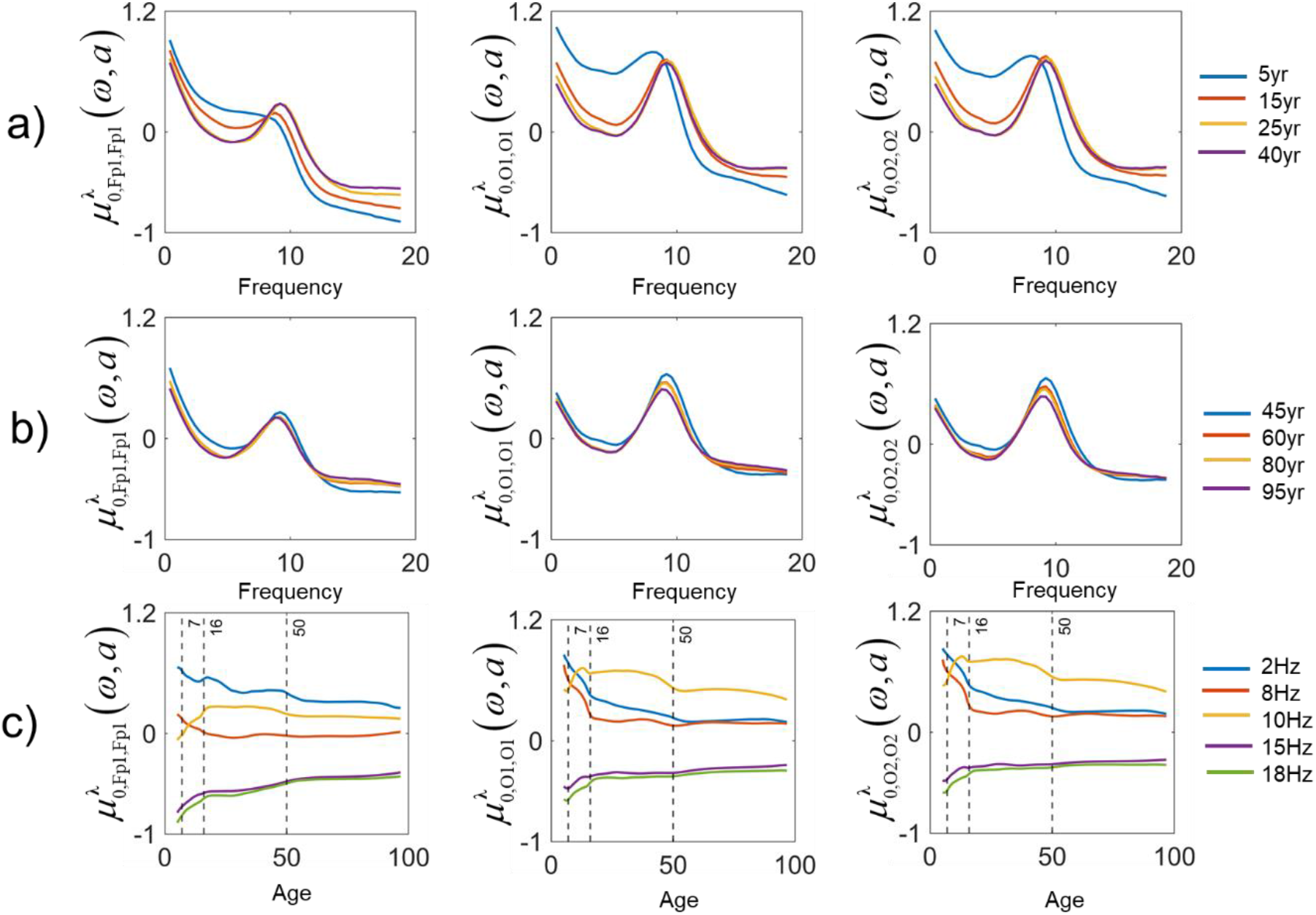
Details of the surrogate log-spectral normative mean. The data in this figure are from the same normative means as in Figure 4-c. The variation of the norm for fixed ages: a) at younger ages (5 yr, 15 yr, 25 yr, and 40 yr). b) at elder ages (45yr, 60yr, 80yr, and 95yr age). c) Changes of the norm with age at specific frequencies (2Hz,8Hz,10Hz, 15Hz, with vertical lines at 7yr, 16yr, and 50yr.

These plots of 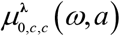 show that children have higher values than other ages at the lower frequency bands (delta and theta). These frequencies decrease dramatically with aging. By contrast, alpha activity increases in magnitude, and its peak moves towards higher frequencies. The alpha peak stabilizes around 25 years until around 40 years. After that age, the alpha peak moves back towards lower magnitudes and frequencies, albeit slightly. For figure 5, detailed illustrations are in the supplement (S4).

Next, we give the norms for the off-diagonal part *c* ≠ *c′*, (Fp1, O1) or (O1,O2) with surrogate coherence and phase developmental surfaces. We show the same types of plots as in Figures 4 and 5. Figures 6, 7, and 8 show the developmental surface of 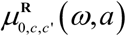 and 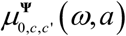. From Figures 6-a and 7, we can see that the coherences 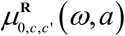 increase from early childhood to around 40 years and after 40 years slightly decrease until around 50, subsequently increasing again after 50. Coherences are maximal at around 10 Hz for all ages. Figures 6-b and 8 show the developmental surface of 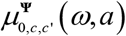. With age, the phase increases from childhood to age 20, later decreasing until 50, to increase afterward until 95 (Figure 8-a to 8-b). The highest phases are at 10 Hz for all ages (Figure 8-c). Here, to better express the range of phases, we define 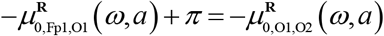

**Figure 6:**
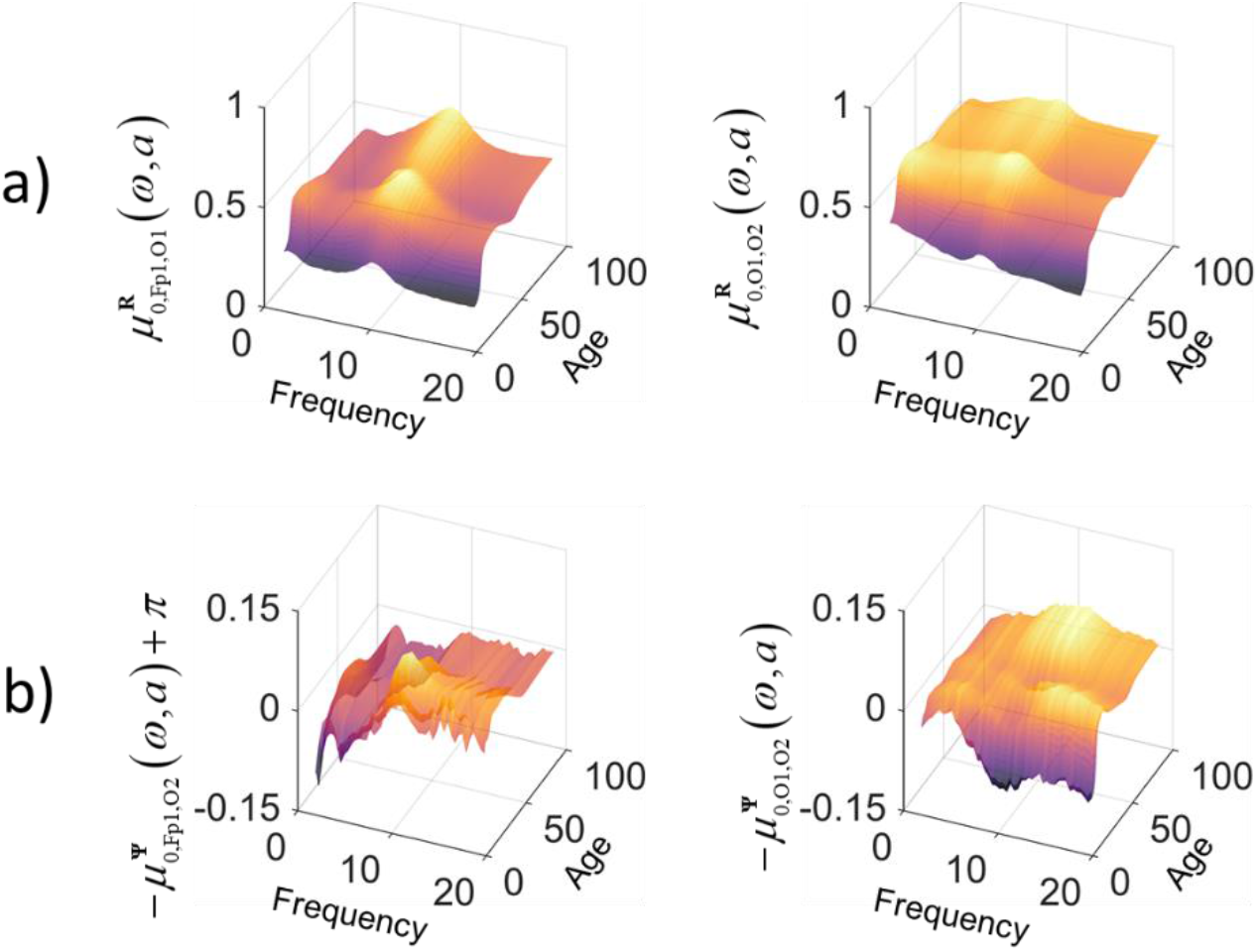
Examples of surrogate coherence and phase normative means. Surrogate normative means of 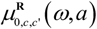 and 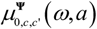 for (Fp1-O1) and (O1-O2). a) Surrogate coherence normative mean for 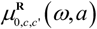; b) Surrogate phase normative mean for 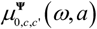.

**Figure 7:**
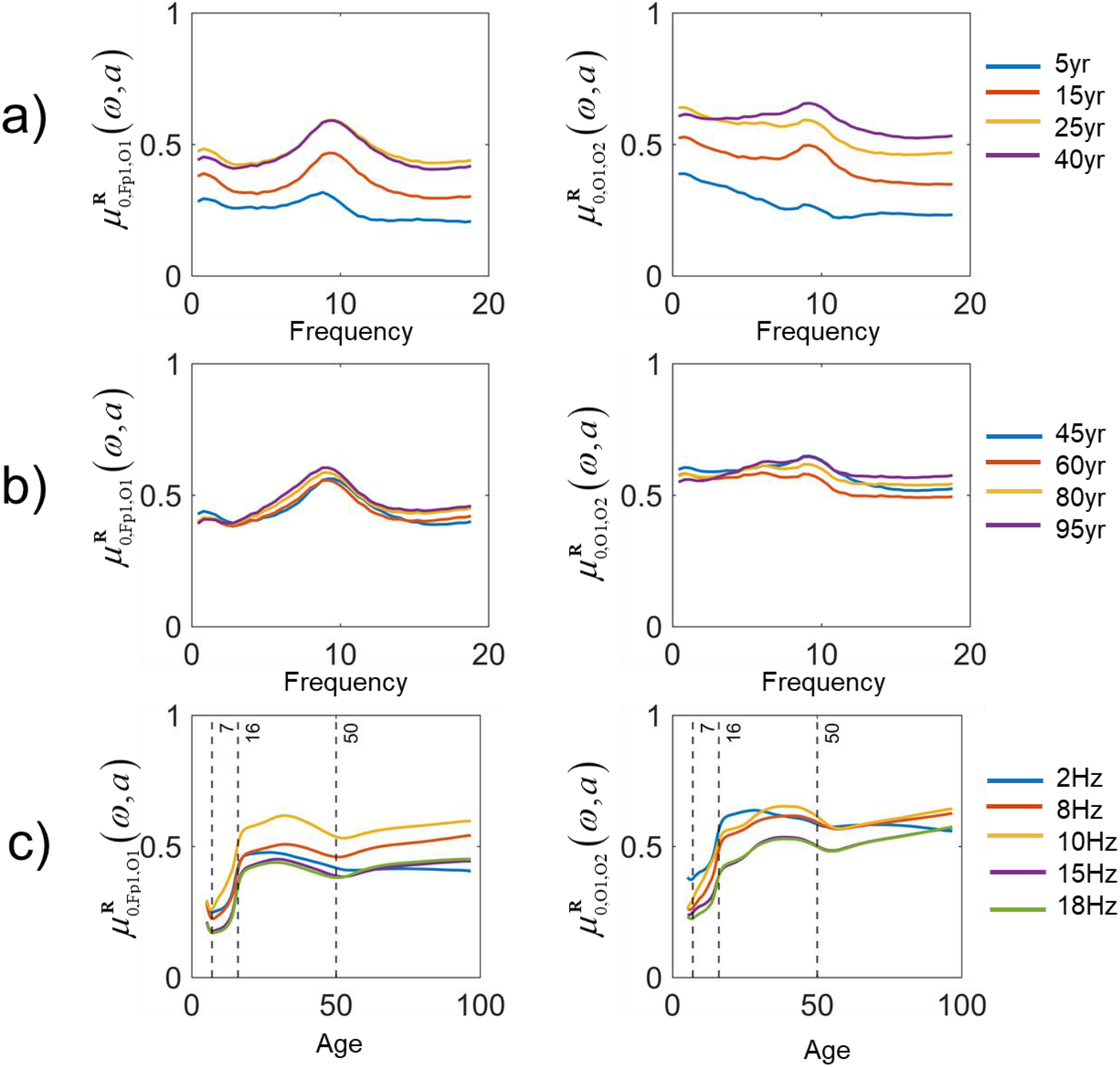
Details of surrogate coherence 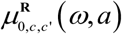 normative mean. The data in this figure are from the same normative means as in Figure 6-a. The variation of the norm for fixed ages: a) at younger ages (5 yr, 15 yr, 25 yr, and 40 yr). b) at elder ages (45yr, 60yr, 80yr, and 95yr age). c) Changes of the norm with age at specific frequencies (2Hz,8Hz,10Hz, 15Hz), with vertical lines at 7yr, 16yr, and 50yr.

**Figure 8:**
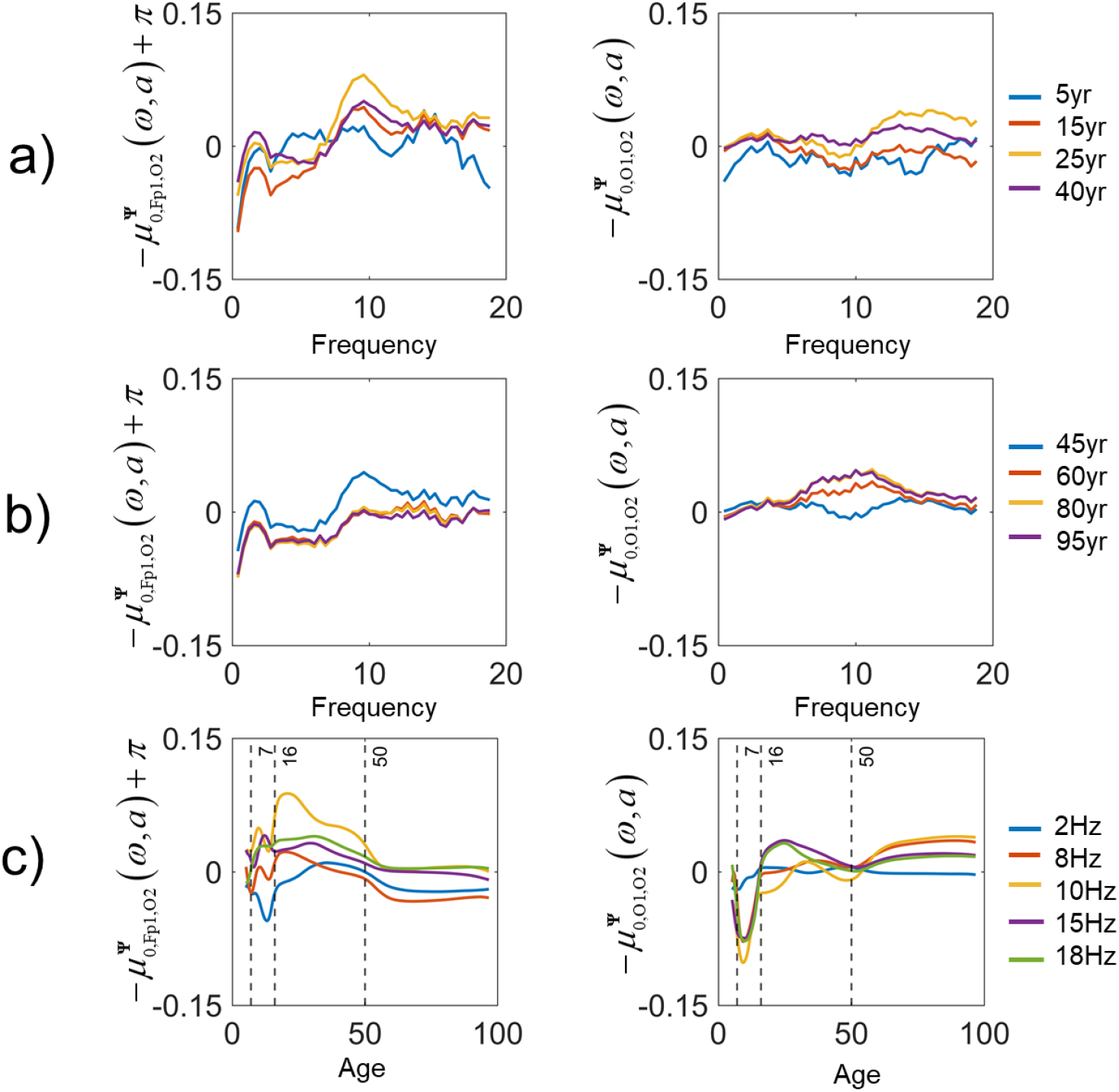
Details of surrogate phase 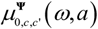 normative mean. The data in this figure are from the same normative means as in Figure 6-b. The variation of the norm for fixed ages: a) at younger ages (5 yr, 15 yr, 25 yr, and 40 yr). b) at elder ages (45yr, 60yr, 80yr, and 95yr age). c) Changes of the norm with age at specific frequencies (2Hz,8Hz,10Hz, 15Hz), with vertical lines at 7yr, 16yr, and 50yr.

### 3.4 Batch Harmonization results

Section 3.2 suggests that qEEG norm models must include batch effects. It remains to see if this effect has practical consequences. A possible effect of ignoring these batch effects is that 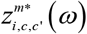 may not distribute as standard Gaussian variables. In this case, harmonizing samples by batch correction using 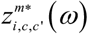 should remedy this.

Indeed, Figure 9-a compares the histograms of 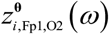 (Figure 9-a) and 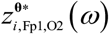 (Figure 9-b) at different batches. With the histogram of the 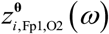 for each batch, suggests they are not standard Gaussian. Note that this effect is not evident in the histogram of the aggregated global z scores pooling all batches (Figure 9-c). The histograms for each batch are more closely gaussian for the harmonized 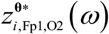 (Figure 9-b), now in correspondence with the appearance of the aggregate for all batches in Figure 9-d.

**Figure 9:**
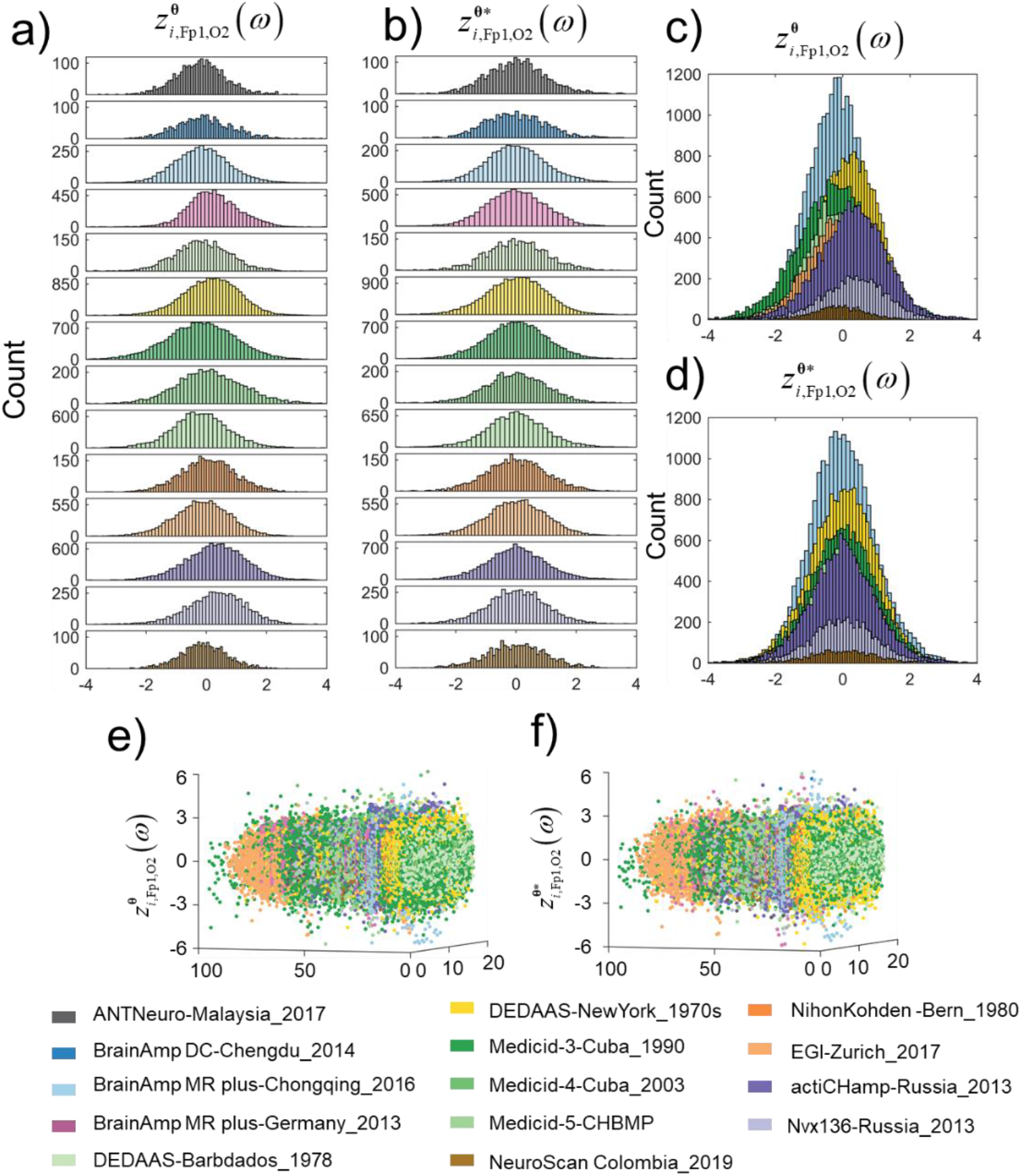
Histograms and scatter plots for 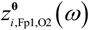 before and after harmonization. Each batch (study) is coded with a different color. a) Histograms of z-score 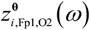 for each batch separately, and c) for all batches superimposed; b) Histograms of batch harmonized HR DP z-score - 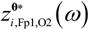 for each batch separately, and d) with all batches superimposed. e) Scatter plot of 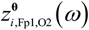 as a function of frequency and age; f) Scatter plot of 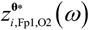 as a function of frequency and age, after harmonization

The harmonization effect is also evident in the scatter plots of the 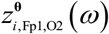 as a function of frequency and age. There are slight but detectable deviations of the z-scores of each batch from a symmetric distribution around the zero plane (Figure 9-e). However, for the 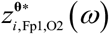, these deviations are removed (Figure 9-f). The corresponding results of 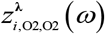 and 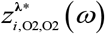 are shown in Figure S4

Additional insight about the effect of harmonization follows from the manifold learning method t-SNE. Let all the subjects as observers, we project the 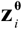 to a common two-dimensional space and see that the data points form clusters, each corresponding to a different batch (Figure 10-a). After harmonization, in the t-SNE plot for the 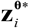, these clusters disappear (Figure 10-b). This batch correction also occurs for multivariate spectral measures. Clustering of 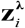 and their disappearances for 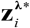 are shown in Figure S5.

**Figure 10:**
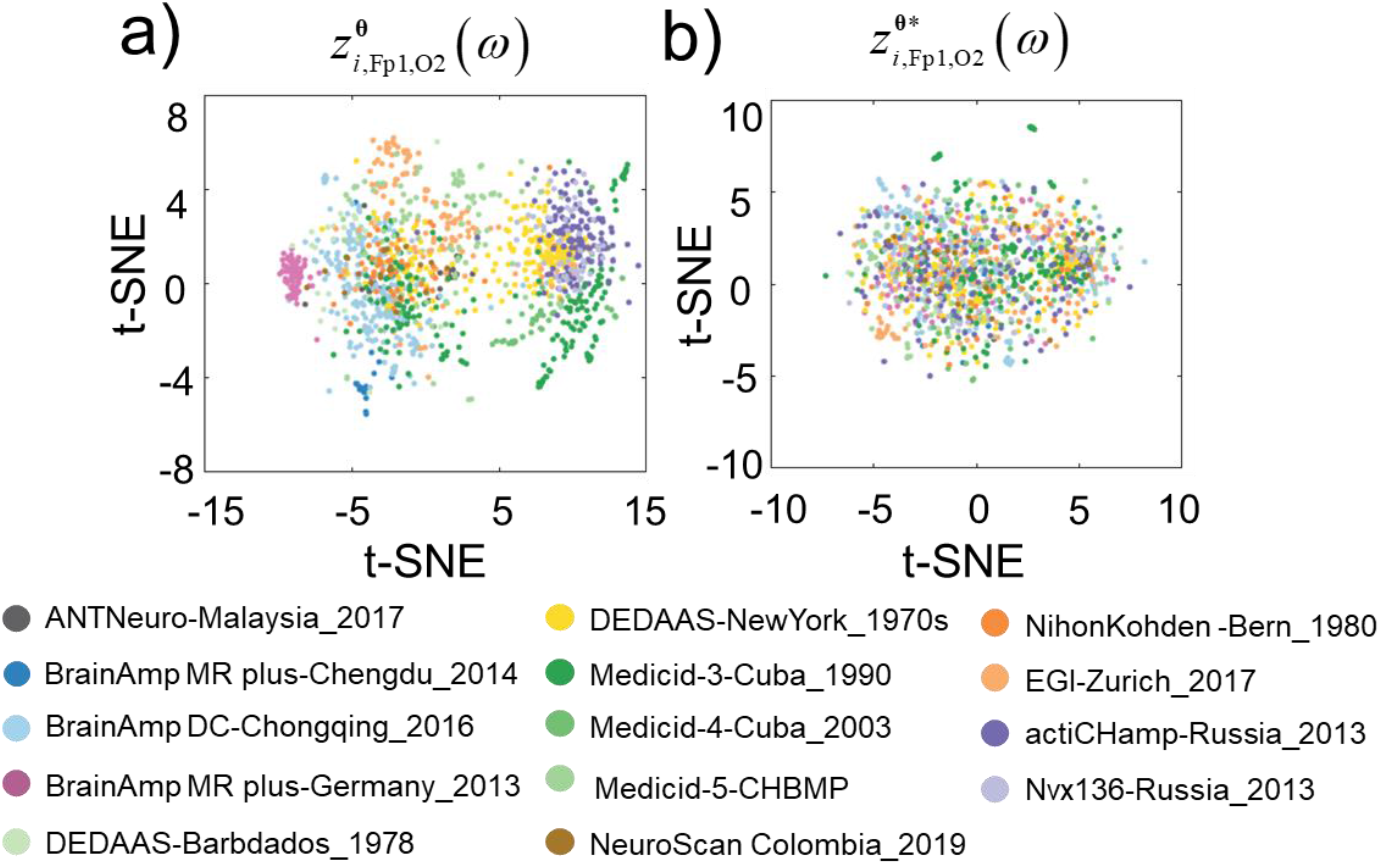
Low dimensional scatter plots of HR DP z-scores 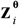, before and after harmonization. The low dimensional representation is a nonlinear mapping (via t-SNE) of z-scores of all HR DPs onto two dimensions. Each axis is log-transformed. Points represent subjects, colored-coded by batch (study). a) Low dimensional unharmonized 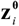 form clusters; b) Low dimensional harmonized 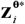 lack cluster structure

These graphical demonstrations of batch effect can also be shown by testing whether 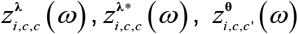, and 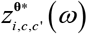 have mean 0, and variance 1—which would follow from these variables having a standard Gaussian distribution. We used MATLAB functions ttest for the mean and vartest for the variance, choosing a significance level of p=0.001 and a Bonferroni correction for multiple comparisons. Table VIII shows there are 73% null hypotheses rejected for the mean values and variance for 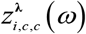. 66% null hypotheses rejected for the mean and 75% null hypotheses rejected for the variance for 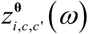. After harmonization, the statistic test for 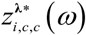 and 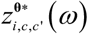 is never rejected, providing further assurance that batch effects are corrected.

**Table VIII:**
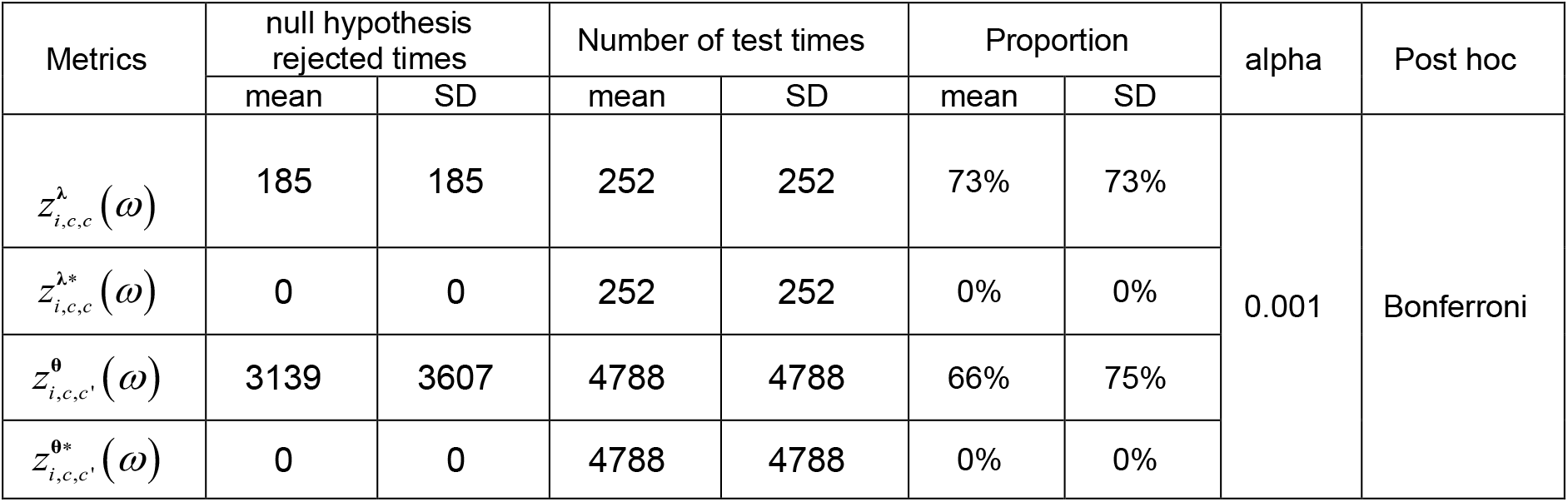
The effect of harmonization on the mean and SD of z-scores

## 4 Quality control

We implemented three distinct stages of quality control:

1. The first filter for quality control and correction or elimination of outliers was at each recording site. At each site, the batches submitted had to be part of a normative study or control group with explicit inclusion and exclusion criteria Appendix A-II.).
2. As mentioned before, the centralized processing team did not have access to the raw EEG data but rather only to the cross-spectra. Preliminary quality control for each sample consisted of visual inspection of a) the topographic map of 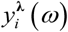 at representative frequencies and b) the 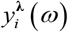 from 1.1718Hz to 19.1394Hz. At least two experienced experts carried out the independent evaluation to avoid subjectivity. The criteria for rejection of cases was overall extreme deformation from expected patterns. There was no attempt to enforce a “very typical pattern” such as the presence of an alpha peak since this may be absent in many normal subjects. Examples of accepted and rejected cross-spectra are shown in Figure S1, where a) is an accepted sample, b) is a rejected sample whose highest power was not at the occipital lobe. Additionally, the spectra are almost flat, indicating the prevalence of noise.
3. Once all batches were gathered, we conducted a machine-learning check for outliers. This centralized second stage quality control step is explained below in detail.

In qEEG research, as in all forms of data science, outliers are a problem that violates this assumption and can derail model estimation and comparison, and inference. Outliers are a common problem that can arise at any step in the whole qEEG pipeline due to: recording artifacts, mistakes in electrode ordering, missteps in data processing, and other factors. we wish to ensure the simple probabilistic interpretation of the DP z-scores 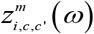 by guaranteeing (as much as possible) that they conform to a multivariate Gaussian distribution. we leveraged the fact that the maximum likelihood covariance estimate (MLE) estimator is sensitive to outliers that spill over to the derived Mahalanobis Distance (MD).

Iterative robust methods for robust MD (Leroy and Rousseeuw, 1987) calculate a corrected MD and make it easy to diagnose multivariate normality(Olive, 2004) and identify outliers samples. Specifically, before attempting the construction of harmonized norms, we detected and eliminated outlier subjects for 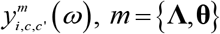 values separately. We first calculated each subject’s 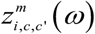 according to the model equation (4). We then create a data matrix where each subject is an observation (n=1564), measured on the 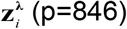 (p=846) and 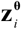 (p=8037).

Because of the large p/ n ratio, we first carry out a nonlinear data mapping to a low dimensional component space using the t-distributed stochastic neighborhood embedding (t-SNE) (Van der Maaten and Hinton, 2008). This reduction allows us to compute both the Traditional and Robust Mahalanobis distances (CMD, RMD) of each sample. We employ the robust estimates Minimum Covariance Determinant (FAST-MCD) method (Rousseeuw and Driessen, 1998) for the normative sample’s mean and covariance matrix.

It is then convenient to detect outliers from the D-D plot (Scatter plot of CMD versus RMD). In the absence of outliers, the data points cluster around the line y=x. By contrast, outliers deviate from this pattern with a practical threshold being 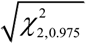, the value of inverse chi-square cumulative distribution with two degrees of freedom for probability=0.975 After this analysis, we found that most data samples are closely grouped in the low dimensional space for 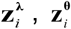 (Figure 11-a,c). The exceptions are some outlier points, as shown in Figure11-d. We can numerically identify these outliers quickly by inspection of the DD plot (Figure 9-b and d). Non-outlier data points cluster tightly around the y=x line since their RMD and CMD should be similar. Outliers have a larger than expected RMD and CMD, as quantified by the Chi-square criteria (vertical and horizontal red lines of coordinate axes) (Figure 11-b and d).

**Figure 11:**
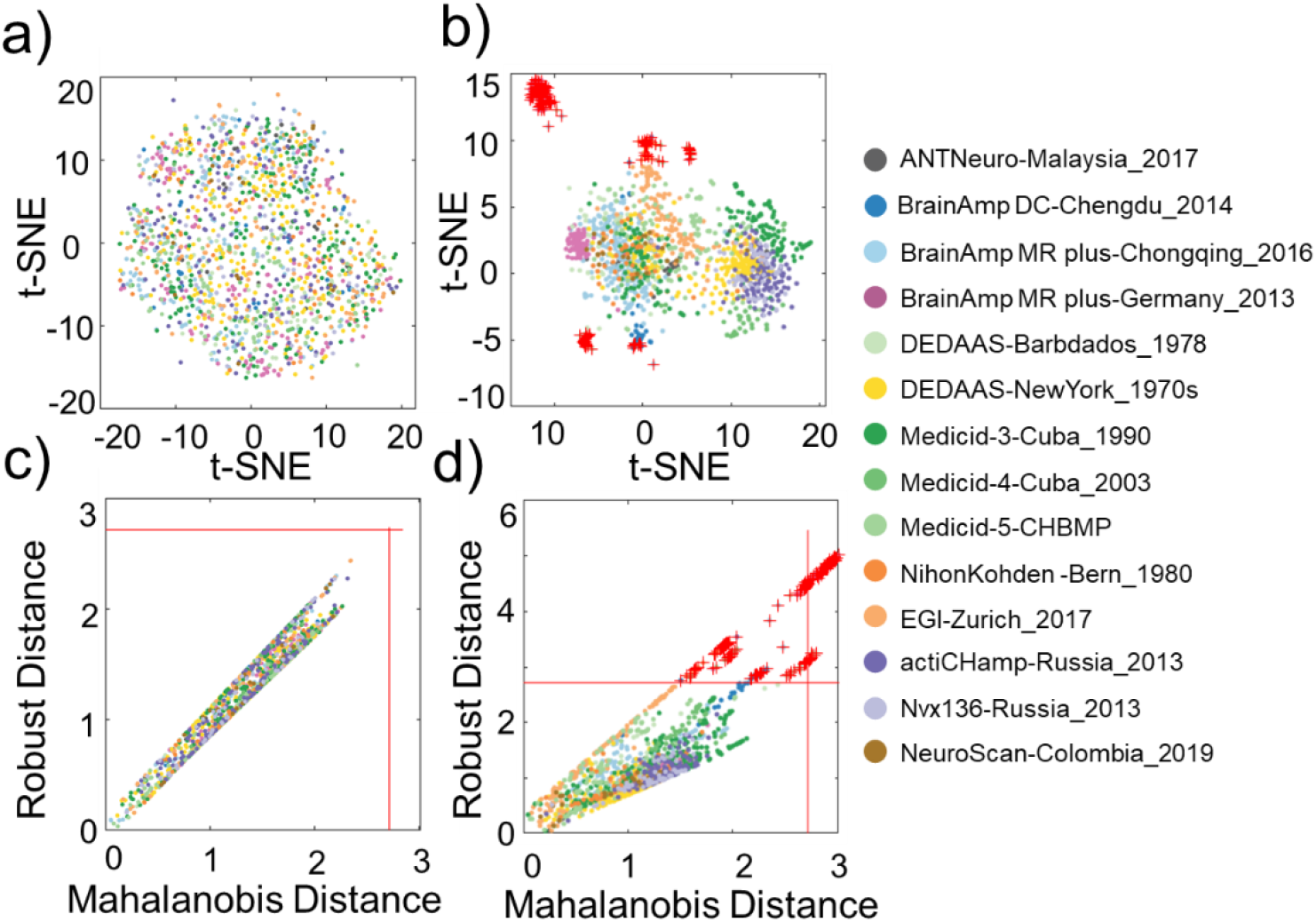
Outlier detection for traditional log-sepctrum and HR z-scores. Each batch (study) is coded with a different color. Subplots a) and c) are two-dimensional representations of DPs via t-SNE, for traditional logspectrum 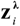 and HR z-scores, 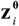. Subplots b) and d) are the corresponding D-D (Robust-Mahalanobis distance versus Mahalanobis distance) plots with limits for outlier detection set at the conventional level, confirming the existence of outliers for HR z-scores. Dots indicate accepted sample points and red crosses outliers. There are no 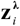. Outliers.

Centralized quality control was carried out iteratively, with feedback about each batch at each site. On several occasions, we detected mistakes in using the gatherer program or in selecting the normative sample—evidenced by a complete batch consisting of outliers. An example of this type of error was a batch with the EEG channels ordered incorrectly— correction of this order eliminated the majority of outliers. With these types of errors corrected, the number of outliers diminished considerably.

As a consequence of this outlier detection step, we identified and eliminated 191 subjects from the samples used for the norm calculation of 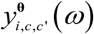 which reduced the final samples number to 1373. By contrast, there are no outliers for 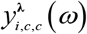.

## 5 Validation of the HarmMNqEEG norms for classification

To check the validity of our new harmonized qEEG norms, revisited the problem of using the qEEG to classify school-children who suffered from Protein Energy Malnutrition limited to the first year of life (BMal) and to distinguish them from healthy classmate controls (BCtrl), who were matched by age, sex and handedness. Prior work is described in Bringas Vega et al. (2019) Taboada-Crispi et al. (2018). This work is part of the Barbados Nutrition Study-- a project that is still ongoing for nearly half a century (Galler et al., 1983a, 1983b).

With the 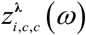, we previously achieved excellent robust elastic-net discrimination between BCtrl and BMal. Those z-transforms were obtained using the CU1990 qEEG norms (Bosch-Bayard et al., 2020)

With the new tools developed in this paper, we gauge the effect on the discrimination between the two groups of three enhancements a) Use of multinational instead of a national norm, b) Use of 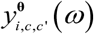 instead of 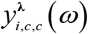, and c) Use harmonized multinational norms.

To answer these questions, we constructed several datasets as shown in the Venn diagram (Figure 12-a):

- Barbados 1978 malnutrition (BMal) comprising *Ni* = 44 samples;
- Barbados 1978 Control (BCtrl) comprising *Ni* = 62 samples.
- In this case, the norms used to calculate z-scores were the complete HarMNqEEG dataset (MN) but excluding the Barbados controls (MN\BCtrl). This modification of the normative data set avoided validation bias,

**Figure 12:**
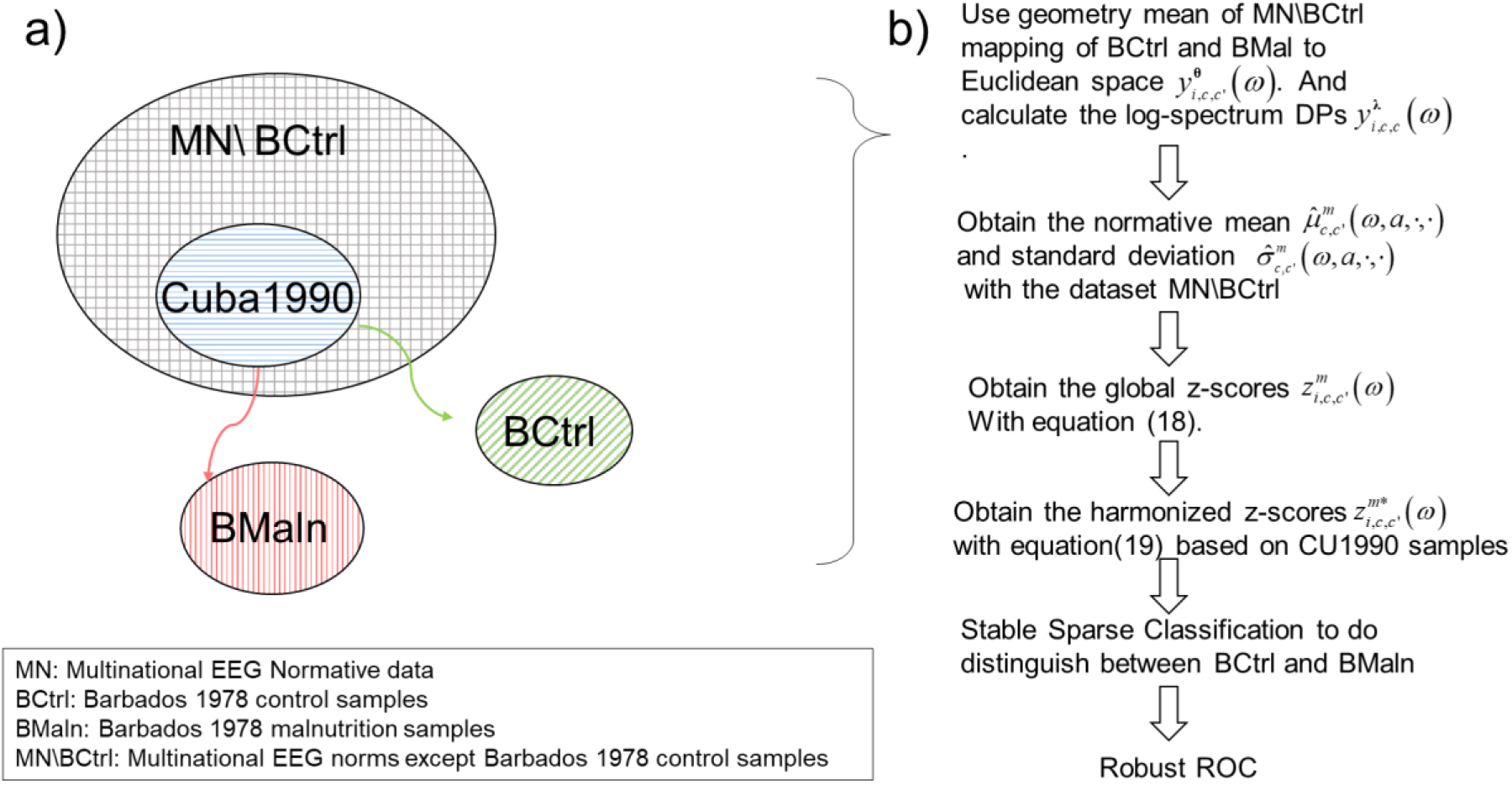
Structure of training and test sets for evaluating the accuracy of different qEEG DPs to detect early protein-energy malnutrition. The different datasets are shown in a Venn diagram in a). The normative dataset used to calculate the 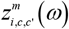 is the complete Multinational Normative dataset, excluding the Barbados Controls (MN\ BCtrl). Batch corrected 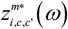 are obtained using the batch information in the Cuba1990 study. The steps for the evaluation of diagnostic accuracy are outlined in b).

After the formation of these sets and training tests, we processed two types of test DPs for the validation (Figure 12-b):

i. Obtain the 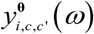 of BMal and BCtrl using the geometric mean of MN\BCtrl when applying the centering operator **M** as in the equation (7). And calculate 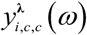 of BMal and BCtrl;
ii. Estimate the normative mean 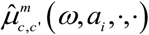 and SD 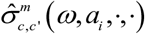 with the dataset MN\BCtrl;
iii. Obtain the global z-scores 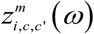 as in the formula (18);
iv. Obtain the batch harmonized z scores 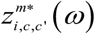 as in the formula (19).

To carry out the batch correction in step iv, and for lack of a larger sample of BCtrl, we plugged in the CU1990 random effect estimator. We base our choice of the CU1990 sample for the batch correction on the social, ethnic, and climatic similarity between Cuba and Barbados and prior studies that indicated that the Cuban norms describe the variability of the BCtrl group (Taboada-Crispi et al., 2018). Thus, we enter the statistical learning procedure with the following types of z-scores for BMal and BCtrl:

| Type of DPs | $y_{i,c,c}^{\lambda}(\omega)$ | $y_{i,c,c'}^{\theta}(\omega)$ |
| --- | --- | --- |
| qEEG norms | $z_{i,c,c}^{\lambda}(\omega)$ | $z_{i,c,c'}^{\theta}(\omega)$ |
| HarMNqEEG norms | $z_{i,c,c}^{\lambda*}(\omega)$ | $z_{i,c,c'}^{\theta*}(\omega)$ |

We evaluate the discriminatory power of z-score types with the elastic-net regression using a stable sparse classifier (Bosch-Bayard et al.,2018). The stable sparse classifier, also used for this same dataset in Bringas Vega et al. (2019), uses multiple resampling to select a stable set of predictors and then (with additional independent resampling) estimate the ROC curves for discrimination between both groups. We plot the ROC curve and report the Area under the ROC curve (AUC) for each type of z-score. Besides the AUC for the whole ROC curve, we also report the standardized partial AUC (spAUC) (McClish, 1989) for the discriminant scores that produce 10% and 20% false positives ratio (FPR). These partial AUC curves are of even greater interest in clinical settings, which usually require low false-positive rates. Accompanying these ROC curves are their probability distribution functions, also obtained via the resampling method.

In figure 13, we first show the ROC curves of these four metrics (Figure 13-a). Obviously, 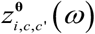 and 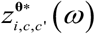 allow higher accuracy than 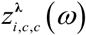 and 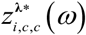. Harmonization boosts total accuracy from 0.828 to 0.87 when comparing 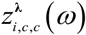 and 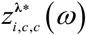. Likewise, there is an increase from 0.943 to 0.952 when comparing 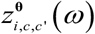 and 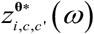. A valuable output of the stable robust classier of Bosch is the probability density functions of the AUC and spAUC for the different measures. From figure 13-B, we see that the AUC of 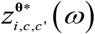 is significantly larger than that of 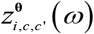, indicating that harmonization is beneficial for accuracy. However, the ROC curves become very close for lower False Positive Rates (FPR). The spAUC at 20% FPR still shows a modest edge for harmonization. However, the situation reverses for even lower FPR (10%), with harmonization lowering the spAUC. Table IX, we summarize the AUC for the different measures compared under different FPRs.

**Figure 13:**
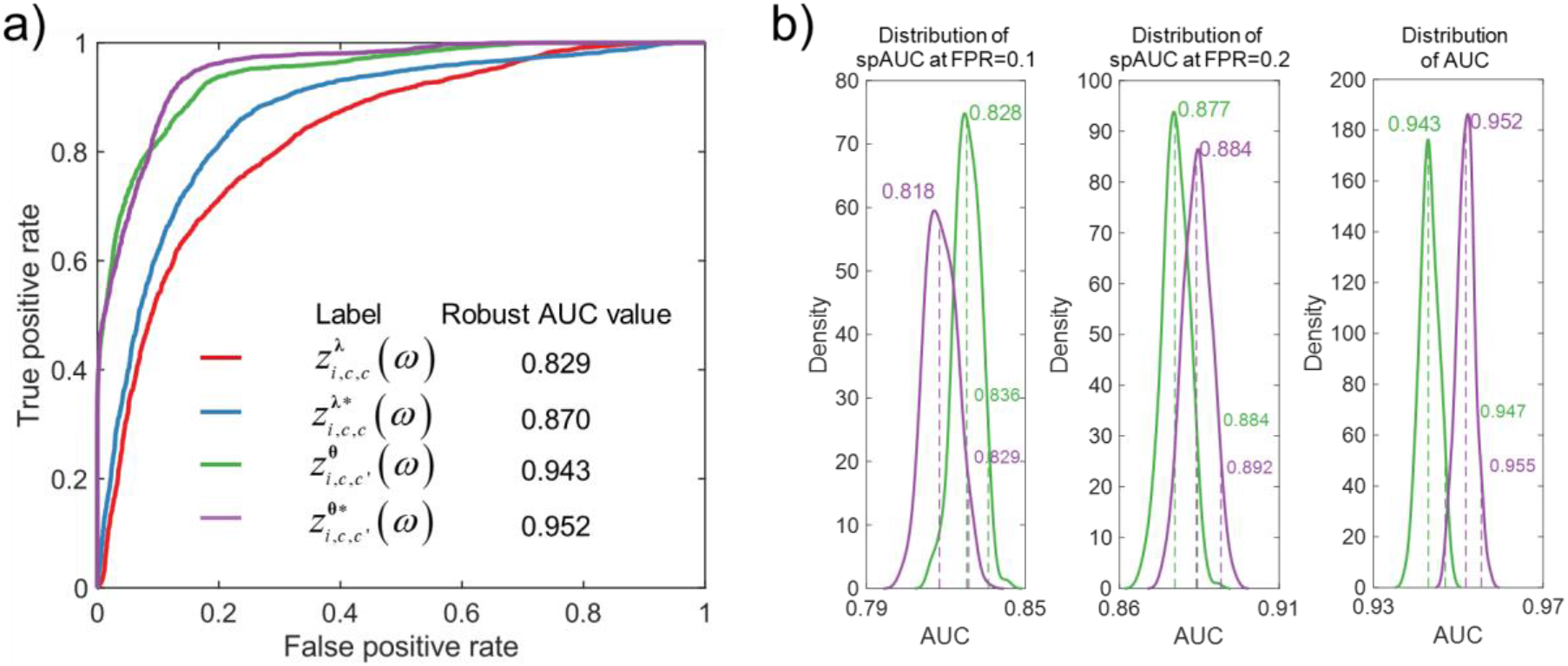
Diagnosis accuracy in detecting early protein-energy malnutrition based on different types of qEEG DPs. Shown in subplot a) are Receiver Operator Curves (ROC) for discriminant functions to distinguish children with protein-energy malnutrition, based on four types of DPs: red 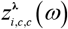, blue 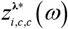, green 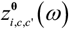, and purple 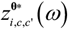. The inset shows the robust Area under the ROC curve (AUC) for these types of DPs. In subplot b), the distribution of spAUCs for HR DPs, before and after harmonization. These are spAUC values for 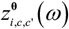 and 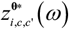 at 10%, 20%, and 100 % (full).

**Table IX:**
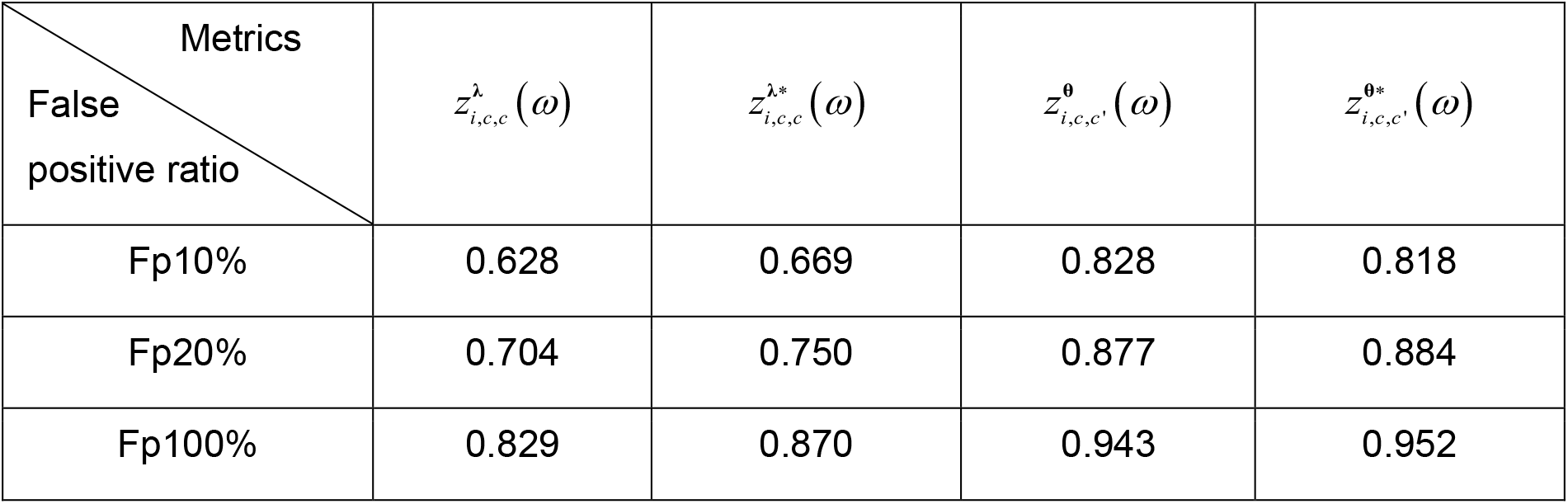
Standardized sparse AUC (spAUC) values of model validation

Moreover, the higher accuracy for 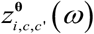 and 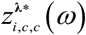 are achieved with a much smaller number of features. As shown in Figure 14, a and b show the selected features of 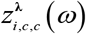 and 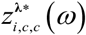 grouped in four frequency bands (1-4Hz, 4-8Hz, 8-13Hz, 13-20Hz). There are more frontal and occipital electrodes involved, like O2 and 8. Figures 14-c and 14-d show the selected features of 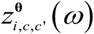 and 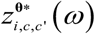 where, more connections between frontal with occipital, e.g., F8-O1.

**Figure 14:**
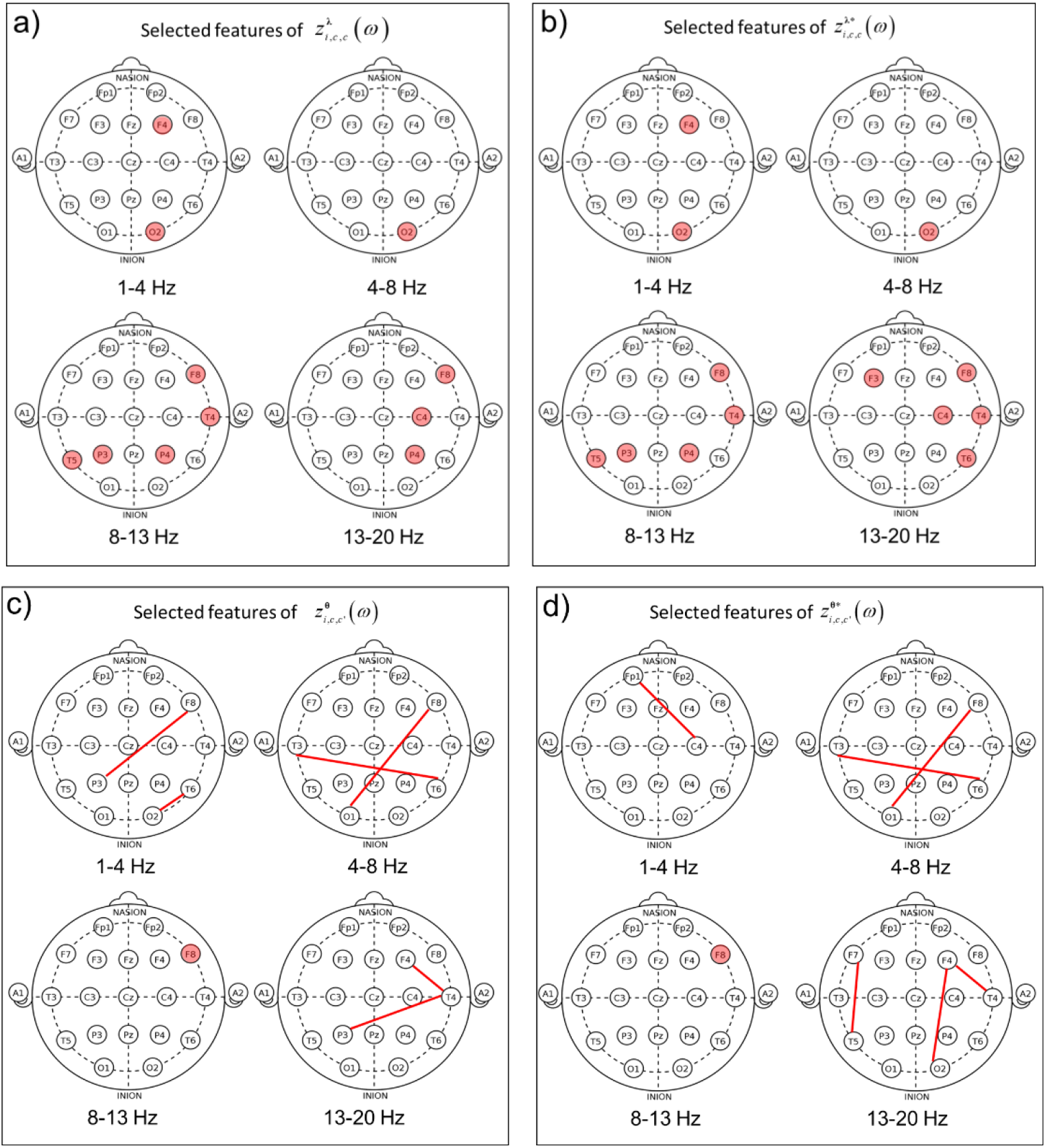
Topography of features selected for detection of malnutrition based on different types of qEEG DPs. The features selected by the robust classifier are indicated in a plot of the 10/20 channel system. They are classified into the four traditional EEG frequency broad bands (though narrowband). A red electrode indicates features related to a single channel. A red link between two electrodes indicates a feature selected for those two channels. a) Selected features for 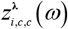; b) Selected features for 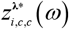. c) Selected features for 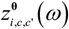. a) Selected features for 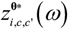.

## 6 Discussion

This paper presents the HarMNqEEG project, an international qEEG normative collaboration intended to avoid racial and socioeconomic bias. It is one of the main projects of the Global Brain Consortium (https://globalbrainconsortium.org/). The construction of unbiased qEEG norms and methods to quantify Brain Developmental Dysfunctions (BDD) is essential for providing easily accessible neuroimaging tools for use in public health settings worldwide. The model chosen for collecting data was offline processing at each site using the MATLAB program “data_gatherer,” which produced batches of samples, each sample consisting of an EEG cross-spectral tensor and anonymized meta-data. Thus samples from different sites were guaranteed to be fully compatible, save possible instruments and site variables. This compatibility was further enhanced by imposing a minimalistic set of recording requirements based on the IFCN 10/20 EEG recording montage and a limited range of EEG frequencies to analyze.

We decided to avoid sharing raw EEG data from recording sites. This choice facilitated the incorporation of diverse groups with varying ethical and administrative constraints. Data collection took less than 3 months to complete. It did, however, require intensive visual and numerical quality control at different stages of the processing pipeline. Other initiatives such as COINSTAC (Gazula et al., 2020) go to wholly and further decentralize processing, a vision that is the next logical step for our project. We are currently studying the outliers detected (in section 4) in the sample to propose methods to avoid future occurrences.

Central processing of the cross-spectral tensors produced two sets of qEEG Descriptive Parameters (DPs). The traditional narrowband log-spectra was included for backward compatibility and comparison purposes. Importantly we introduce in this paper a new type of EEG qEEG DP based on Hermitian Riemannian geometry. This development is essential since cross-spectral tenors occupy a highly nonlinear manifold. To this end, we define a Riemannian vectorization operator that transforms the cross-spectral tensor to a vector that closely follows a multivariate gaussian distribution in Euclidean space. An essential ingredient of the Riemannian vectorization is the matrix-logarithmic transform of the coss-spectral matrices. Leonard and Chiu (Leonard and Hsu, 2001) proposed this transformation to obtain an unconstrained multivariate gaussian vector from covariance matrices.

We note that the Riemannian Geometry of Symmetric Positive Definite matrices has been widely used for Brain-Computer Interface applications (Barachant et al., 2013; Yger et al., 2017). Recently, in a seminal series of papers, (Sabbagh et al., 2020; Engemann et al., 2021) showed the superiority of HR DPs to predict brain age with MEEG/EEG. Our work differs from these previous applications in two essential aspects. The first is the use of Hermitian (not Real) Riemannian Geometry since the frontal faces of the cross-spectral tensor are cross-spectral matrices, Hermitian covariance matrices of EEG Fourier coefficients. A second difference is that rather than predating age by the HR DPs, we consider age as an independent variable to predict Riemannaian DPs.

The added conceptual and computational machinery of Riemannian Geometry seems to be beneficial. With this framework, we can now norm not only the frequency-resolved EEG activities at each channel (via the spectra) but also the functional connectivity between electrodes (reflected in the cross-spectra). The matrix-logarithmic transform allows an integrated transformation towards normality of activity and connectivity that was impossible with isolated transformations of the elements of the cross-spectral matrix.

The multinational character of the datasets collected in the project allowed us a first look at qEEG harmonization. We believe the HarMNqEEG project is one of the first efforts to check for “batch” effects” in qEEG multisite datasets and propose computing subject z-scores from batch-free qEEG norms. To our knowledge, this is the first attempt to create a statistically valid qEEG norm for cross-spectral tensors and their derived measures (coherence and phase).

Our work confirmed that a large part of the variance of traditional and HR DPs depends on frequency and age, underscoring the need for frequency and age-dependent norms to detect BDD. This fixed-effect dependency has been reported many times for traditional log-spectra, including several papers from our group (Taboada-Crispi et al., 2018; Bosch-Bayard et al., 2001; Bringas Vega et al., 2019). Indeed, the shape of the norms or “developmental surfaces” for traditional measures hold up with the larger multinational sample. Based on Riemannian geometry, we confirm this frequency and age dependency for the full cross-spectrum. The reconstruction of traditional log-spectral norms from the HR norms (surrogate norms) shows the consistency of the two procedures for the traditional DP set.

Regarding batch differences, note that using a global geometric mean to center cross-spectral tensors forces all data onto a unique normative tangent space. We studied whether batch or sex effects should be retained in the normative equations (Simeon et al., 2021).

These models were compared using the Extended Bayesian Information Criterion. Contrary to our expectation, sex was pruned from the independent variables for fixed and random effects. This lack of sex-dependency requires further study, given the numerous neuroimaging studies for other modalities that find such an effect.

On the other hand, batch effects, specifically site random effects, were evident. The modeling framework allowed the definition of batch-free z-scores. The need for this correction is particularly evident for HR DPs and easily observable in the t-sne plots where harmonization eliminates batch differences.

The readers familiar with the construction of norms may be acquainted with COMBAT (Johnson et al., 2007) and generalized additive model for local scale and shape (GAMLSS) (Rigby and Stasinopoulos, 2005)--harmonization methods widespread to eliminate batch effects. COMBAT captures the effect of biological variables with a linear model, then positing that batch effects have a constant mean and SD. The GAMLSS model, on the other hand, is more general than COMBAT. It allows nonparametric additive modeling of biological effects for the mean, variance, and higher statistical fixed and random effects parameters, but the use of GAMLSS for harmonization is usually restricted linear models (Bethlehem et al., 2021; Dinga et al., 2021). Subsequently, combining COMBAT with generalized additive model (GAM) made it possible to identify nonlinear relationships between age and brain structure (Pomponio et al., 2020). However, together with GAMLSS this class of models is restricted to univariate function.

Our model includes and generalizes both COMBAT and GAMLSS when the latter is restricted to Gaussian noise—which is our case. HarMNqEEG allows multivariate nonparametric functional forms for the biological variables and the random additive effect (sex or batch) for mean and SD. Due to the Gaussianity of our DPs, we did not include models of higher-order statistics as in GAMLSS, but our models are more general for Gaussian data since our code allows the mean and SD functions to be both complex-valued and multivariate, and the variance function can be log additive. Finally, compared with the generalized additive mixed-mode (GAMM) (Lin and Zhang, 1999), though the latter does allow multivariate functions, these are only for the mean. We emphasize that both COMBAT and GAMLSS models were tested in model selection and not retained due to having a higher nEBIC.

We additionally provide evidence that Hermitian Riemannian qEEG can provide higher diagnostic accuracy than traditional qEEG by evaluating the previously well studied Barbados Nutritional Study data set (Bringas Vega et al., 2019; Rutherford et al., 2021b; Taboada-Crispi et al., 2018). We are currently exploring other clinical datasets with this approach.

The present study has several limitations. The most important one is that, despite the substantial increase in sample size (compared to previous studies), it still may be underpowered to detect the effect of some biological variables such as gender. More importantly, the lack of combinations of types of equipment and country precludes separating the effect of these two variables. This issue needs to be the focus of further studies. Additionally, and as mentioned above, the age distribution of the sample is skewed towards younger ages. It is essential to remedy this unbalance of age distributions to improve the analysis of cognitive aging.

A normative study with denser electrode montages would also be desirable to improve localization accuracy, though the present study helps evaluate less intensive EEG examinations.

Finally, though our Narrow Band set of frequency-domain Descriptive Parameters (Szava) was proven to be more informative than the BroadBand variety, it is not entirely satisfactory. A set of DPs based on empirical decompositions of the power spectrum is preferable. Splitting the resting-state power spectrum into a /f background component and an alpha peak has been long studied (Zetterberg, 1969; Pascual-marqui et al., 1988; Donoghue, 2020) The most salient feature of the alpha peak is its alpha peak frequency (IAF), for which excellent developmental studies exist (Voytek et al., 2015; Tröndle et al., 2021) In our formulation (Pascual-marqui et al., 1988) this decomposition was termed the “Xi-alpha” model, and the IAF was identified as the Palpha parameter, which we showed early on to be the spectral parameter best correlated with age (Amador et al., 1989). However, we did not use the Xi-Alpha here approach since the generalization needed for cross-spectra requires completing the multivariate generalization(Pascual-marqui et al., 1988; Valdés et al., 1992) that has an adequate Rimeannian embedding. This project will be the subject of future work.

There are several future directions of this work already being developed. Some of these directions are:

- Extension of the multinational Hermitian Riemannian norms to source space as done already achieved for traditional qEEG (Bosch-Bayard et al., 2001; Bringas Vega et al., 2019)
- Creation of multinational norms for other sets of qEEG descriptive parameters such as microstates (Koenig et al., 2002a)
- Create norms for multivariate Xi-Alpha models of the EEG cross-spectrum (Pascual-marqui et al., 1988; Hu and Valdes-Sosa, 2019; Tröndle et al., 2020)

We have also made available the code to calculate different individual z-scores from the HarMNqEEG dataset.

The results presented in this paper contribute to developing bias-free, low-cost neuroimaging technologies applicable in various health settings, especially those in low resource areas that are at greatest risk for neurodevelopmental disabilities and other brain disorders (Galler et al., 2021).

## Ethics Statement

The studies involving human participants were reviewed and approved by the Ethics Committees of all involved institutions. In all cases, the participants and/or their legal guardians/next of kin provided written informed consent to participate in this study. All data were de-identified, and participants gave permission for their data to be used shared as part of the informed consent process.

## Author contribution

The central processing team comprised: ML, who integrated methodological and software developments, directed quality control and data Curation. Together with PAVS, she carried out the main contribution to paper writing. TK carried out the initial collation of datasets and continued involved in the project. YW contributed to theoretical and software development methods for creating norms with kernel regression. He was also involved in data curation and contributed to writing, reviewing, and editing. JFBB created, with PAVS, the traditional qEEG methods, wrote the gatherer program and supervised the curation of data supplied from all sites. He supervised theoretical developments and was responsible for statistical learning methods to check diagnostic accuracy. MLB was responsible for the conceptualization and organization of normative projects, the supervision of data collection, guarantee of essential resources. She also did write and editing.CLN was responsible with ML and PAVS for developing the essential toolbox for Riemannian geometry. PAVS conceived this line of research, stated the main methodological strategies, organized the Global Brain Consortium project, and carried out its administration (with ACE). He oversaw statistical analyses and conceived the paper’s structure and all stages of writing. The following co-authors (in alphabetical order) contributed data, essential discussions, and paper editing: AIAH, ACE, ANS, ACR, AAG, AV, CATQ, DGA, DPL, DY, LD, EAV, FR, HO, JMA, JRG,JFOG, LSP, LGG, LMC, MJVS, MT, MFBMZ, MRBAR, NSM, NL, PR, TK, TAVA, SH, XL. ML and YW have equal contribution to this paper.

## Acknowledgment

The research was funded by grants (to PAVS) from the National Project for Neurotechnology of the Ministry of Science Technology and Environment of Cuba, the National Nature Science Foundation of China NSFC grant No. 61871105, CNS Program of UESTC (No. Y0301902610100201) and (to MLBV and PAVS) from the Nestlé Foundation (Validation of a long-life neural fingerprint of early malnutrition, 2017). The National Science Foundation of China (to SH) grant with No. 62101003. JFBB was supported by the Fonds de recherche du Québec (FRQ) HBHL FRQ/CCC Axis (246117), the CFREF/HBHL, HIBALL, Helmholtz (252428), and Brain Canada (243030). The team of Malaysia was funded from the Translational Research Grant Scheme, Ministry of Higher Education (TRGS/1/2015/USM/01/6/3) and the Research University Grant (RUI), Universiti Sains Malaysia (1001/PPSP/8012307). The dataset from Russian was collected with the support of the Russian Foundation of Basic Research, grant No. 18-29-13027. The Barbados dataset was funded by the Grants R01 HD060986 (JRG).S

## Data and code available

The shared raw cross-spectrum with encrypted ID is hosted at Synapse.org (https://doi.org/10.7303/syn26712693) and complete access is possible after login in the system. The multinational harmonized norms (HarMNqEEG norms) of traditional log-spectrum and HR cross-spectrum open in Synapse.org (https://doi.org/10.7303/syn26712979). And the corresponding HarMNqEEG code for calculating the z-scores based on the HarMNqEEG norm opened in GitHub, see: https://github.com/LMNonlinear/HarMNqEEG.

## Appendix A

### Appendix A-I: Summary of indexes

**Table A-I:** Summary of indexes

|  |  |
| --- | --- |
| $i$ | Index of individual $i \in \{1, 2, \dots, N_i\}$. |
| $c$ | Index of the channel $c \in \{1, 2, \dots, N_c\}$, $N_c = 19$ including Fp1, Fp2, F3, F4, C3, C4, P3, P4, O1, O2, F7, F8, T3, T4, T5, T6, Fz, Cz, Pz electrodes. |
| $e$ | Index of EEG epochs $e \in \{1, 2, \dots, N_e\}$, the disjoint segments selected from the continuous EEG. |
| $\omega$ | Discrete frequency $\omega = 1, \dots, N_\omega$. When multiplied by $\Delta\omega$ gives the physical frequency. |
| $a$ | Age, which takes value from the subset of $\{a \in \mathbb{R}_+, \cdot\}$. Then $a_i$ denotes the individual's age $i$. The symbol "." indicates all ages. |
| $s$ | Sex, which takes value from the subset of $\{\text{Male, Femal}, \cdot\}$. Then $s_i$ indicates the sex of the individual $i$. The symbol "." indicates all sex. |
| $b$ | Batch, where $b_i$ means batch of individual $i$, the symbol "." indicates all batches. |
| $t$ | Time $t = 1, \dots, N_t$. Physical time is obtained by multiplication with $\Delta t$. |
| $c$ | Index of the channel $c \in \{1, 2, \dots, N_c\}$, here $N_c = 19$ including Fp1, Fp2, F3, F4, C3, C4, P3, P4, O1, O2, F7, F8, T3, T4, T5, T6, Fz, Cz, Pz electrodes. |
| $e$ | Index of EEG epochs $e \in \{1, 2, \dots, N_e\}$, the disjoint segments selected from the continuous EEG. |

### Appendix A-II: Multinational EEG norms dataset

**Table A-II:**
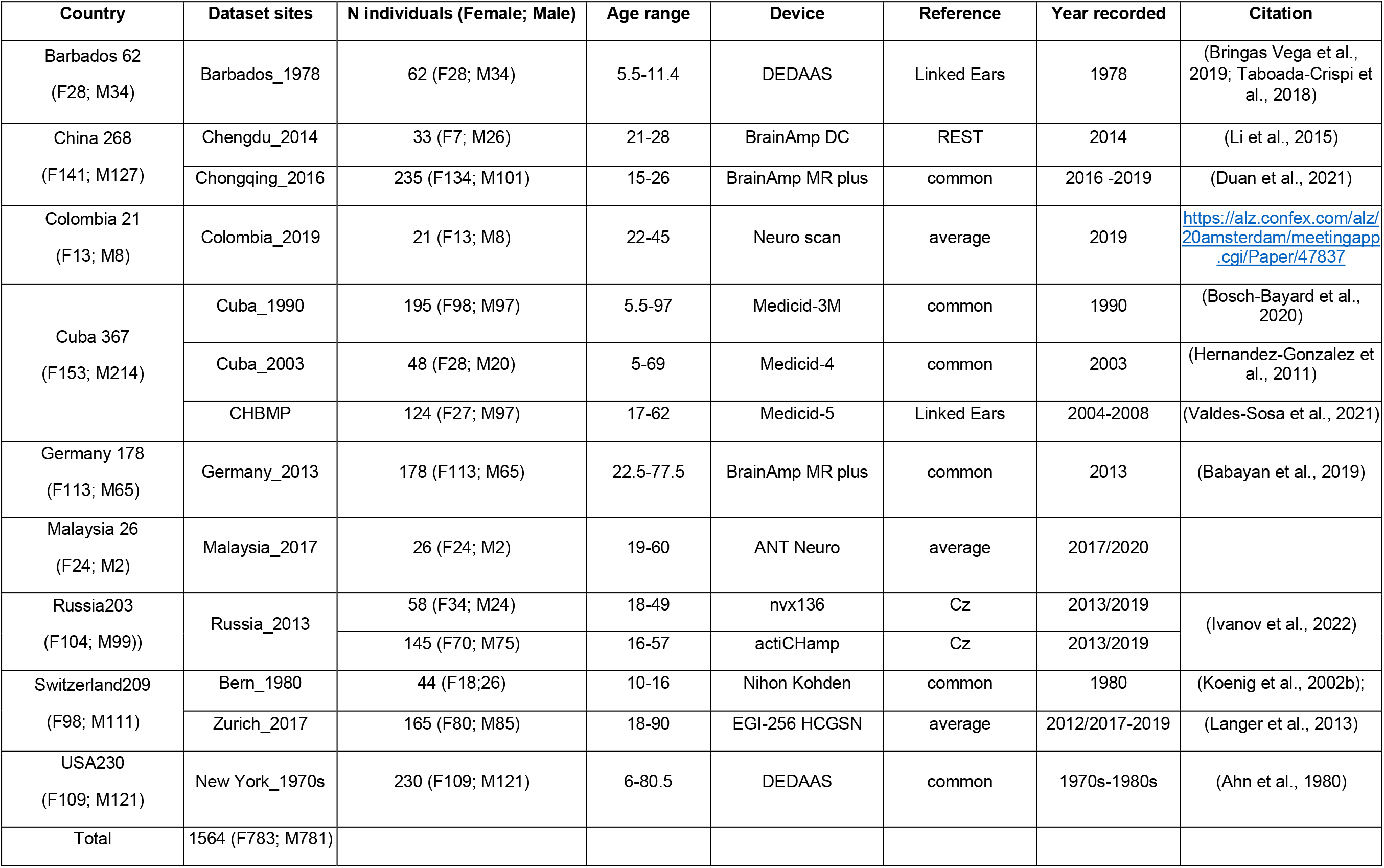
Multi-national EEG norms dataset: including 9 countries, 12 devices and 14 batches.

| Country | Dataset sites | N individuals (Female; Male) | Age range | Device | Reference | Year recorded | Citation |
| --- | --- | --- | --- | --- | --- | --- | --- |
| Barbados 62<br>(F28; M34) | Barbados_1978 | 62 (F28; M34) | 5.5-11.4 | DEDAAS | Linked Ears | 1978 | (Bringas Vega et al., 2019; Taboada-Crispi et al., 2018) |
| China 268<br>(F141; M127) | Chengdu_2014 | 33 (F7; M26) | 21-28 | BrainAmp DC | REST | 2014 | (Li et al., 2015) |
|  | Chongqing_2016 | 235 (F134; M101) | 15-26 | BrainAmp MR plus | common | 2016 -2019 | (Duan et al., 2021) |
| Colombia 21<br>(F13; M8) | Colombia_2019 | 21 (F13; M8) | 22-45 | Neuro scan | average | 2019 | https://alz.confex.com/alz/20amsterdam/meetingapp.cgi/Paper/47837 |
| Cuba 367<br>(F153; M214) | Cuba_1990 | 195 (F98; M97) | 5.5-97 | Medicid-3M | common | 1990 | (Bosch-Bayard et al., 2020) |
|  | Cuba_2003 | 48 (F28; M20) | 5-69 | Medicid-4 | common | 2003 | (Hernandez-Gonzalez et al., 2011) |
|  | CHBMP | 124 (F27; M97) | 17-62 | Medicid-5 | Linked Ears | 2004-2008 | (Valdes-Sosa et al., 2021) |
| Germany 178<br>(F113; M65) | Germany_2013 | 178 (F113; M65) | 22.5-77.5 | BrainAmp MR plus | common | 2013 | (Babayan et al., 2019) |
| Malaysia 26<br>(F24; M2) | Malaysia_2017 | 26 (F24; M2) | 19-60 | ANT Neuro | average | 2017/2020 |  |
| Russia203<br>(F104; M99)) | Russia_2013 | 58 (F34; M24) | 18-49 | nvx136 | Cz | 2013/2019 | (Ivanov et al., 2022) |
|  |  | 145 (F70; M75) | 16-57 | actiCHamp | Cz | 2013/2019 |  |
| Switzerland209<br>(F98; M111) | Bern_1980 | 44 (F18;26) | 10-16 | Nihon Kohden | common | 1980 | (Koenig et al., 2002b); |
|  | Zurich_2017 | 165 (F80; M85) | 18-90 | EGI-256 HCGSN | average | 2012/2017-2019 | (Langer et al., 2013) |
| USA230<br>(F109; M121) | New York_1970s | 230 (F109; M121) | 6-80.5 | DEDAAS | common | 1970s-1980s | (Ahn et al., 1980) |
| Total | 1564 (F783; M781) |  |  |  |  |  |  |

### Appendix A-III: Batches defined equipment, and country

**Table A-III:** Batches defined equipment, and country

| Device<br>Country | BrainAmp<br>MR plus | BrainAmp DC | actiCHamp | DEDAAS | EGI-256<br>HCGSN | Medicid-3M | Medicid-4 | Medicid<br>-5 | NihonKohd<br>en | nvx136 | ANT<br>Neuro | Neuro<br>scan |
| --- | --- | --- | --- | --- | --- | --- | --- | --- | --- | --- | --- | --- |
| Barbados |  |  |  | DEDAAS-Barbdados_<br>1978 |  |  |  |  |  |  |  |  |
| China | BrainAmp<br>MR plus-Chongqing_20<br>16 | BrainAmp DC-Chengdu_2014 |  |  |  |  |  |  |  |  |  |  |
| Cuba |  |  |  |  |  | Medicid-3M-Cuba_1990 | Medicid-4-Cuba_200<br>3 | Medicid-5-CHBMP |  |  |  |  |
| Colombia |  |  |  |  |  |  |  |  |  |  |  | NeuroScan-Colombia_2<br>019 |
| Malaysia |  |  |  |  |  |  |  |  |  |  | ANTNeuro-Malaysia_2<br>017 |  |
| Germany | BrainAmp<br>MR plus-Germany_201<br>3 |  |  |  |  |  |  |  |  |  |  |  |
| Russia |  |  | actiCHamp-Russia_2013 |  |  |  |  |  |  | nvx136-Russia_2<br>013 |  |  |
| Switzerland |  |  |  |  | EGI-Zurich_201<br>7 |  |  |  | NihonKohden<br>-Bern_1980 |  |  |  |
| USA |  |  |  | DEDAAS-NewYork_1<br>970s |  |  |  |  |  |  |  |  |

---

**Figure S1.**
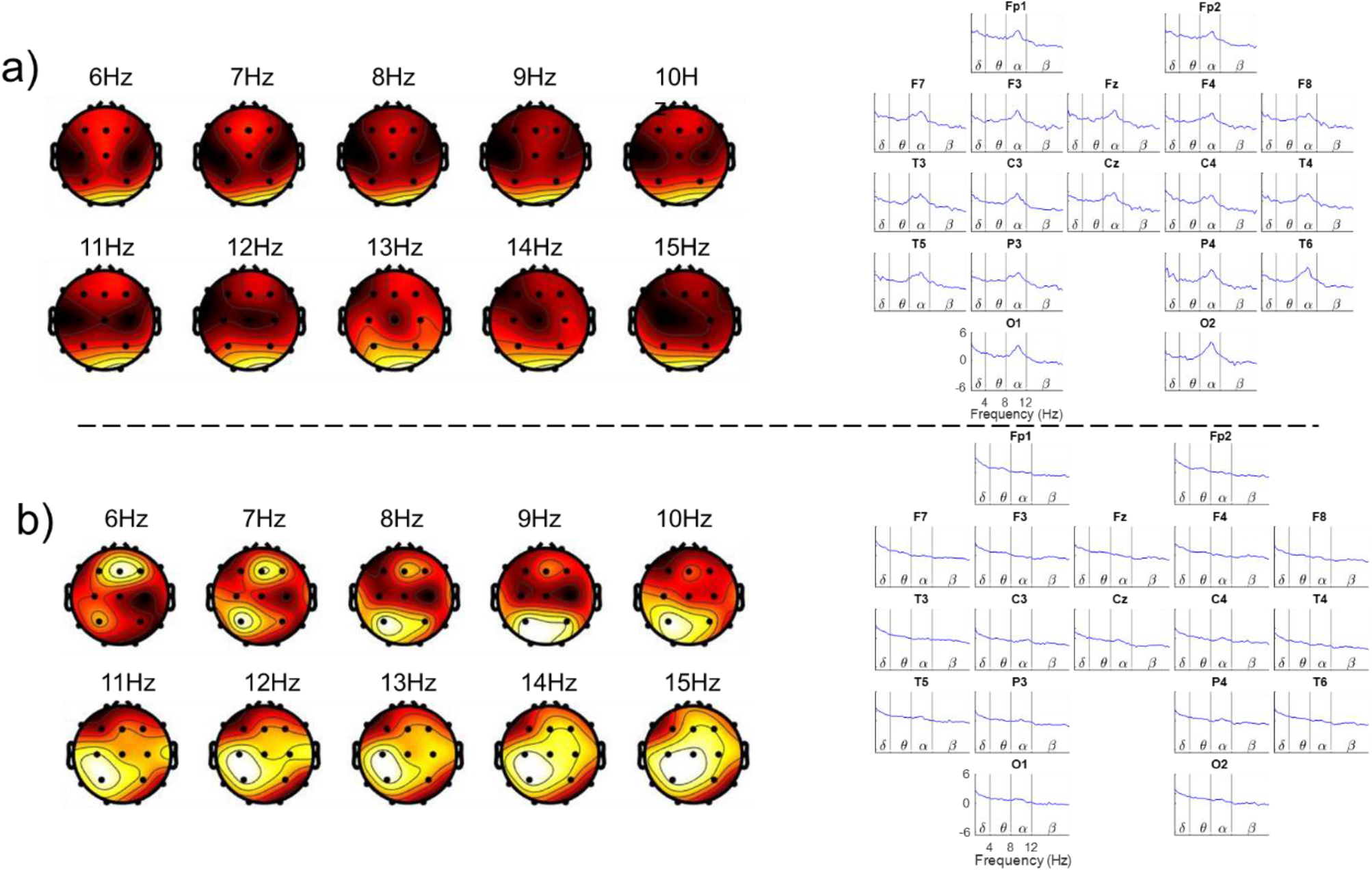
Initial visual quality based on log-spectra of 10/20 system. Initial visual quality based on log spectra of 10/20 system. Initial quality control was carried out before preprocessing, based on two types of log-spectral plots. The (interpolated) topographic maps of the log-spectrum at different specific frequencies are on the left. On the right is the plot of the log-spectra for each electrode. The figure shows two examples of cases, on the top an accepted case with topography and spectra conforming to most normative cases. On the bottom, a rejected case that was atypical on both plots. This case generated a request to the recording site to do additional quality control.

**Figure S2.**
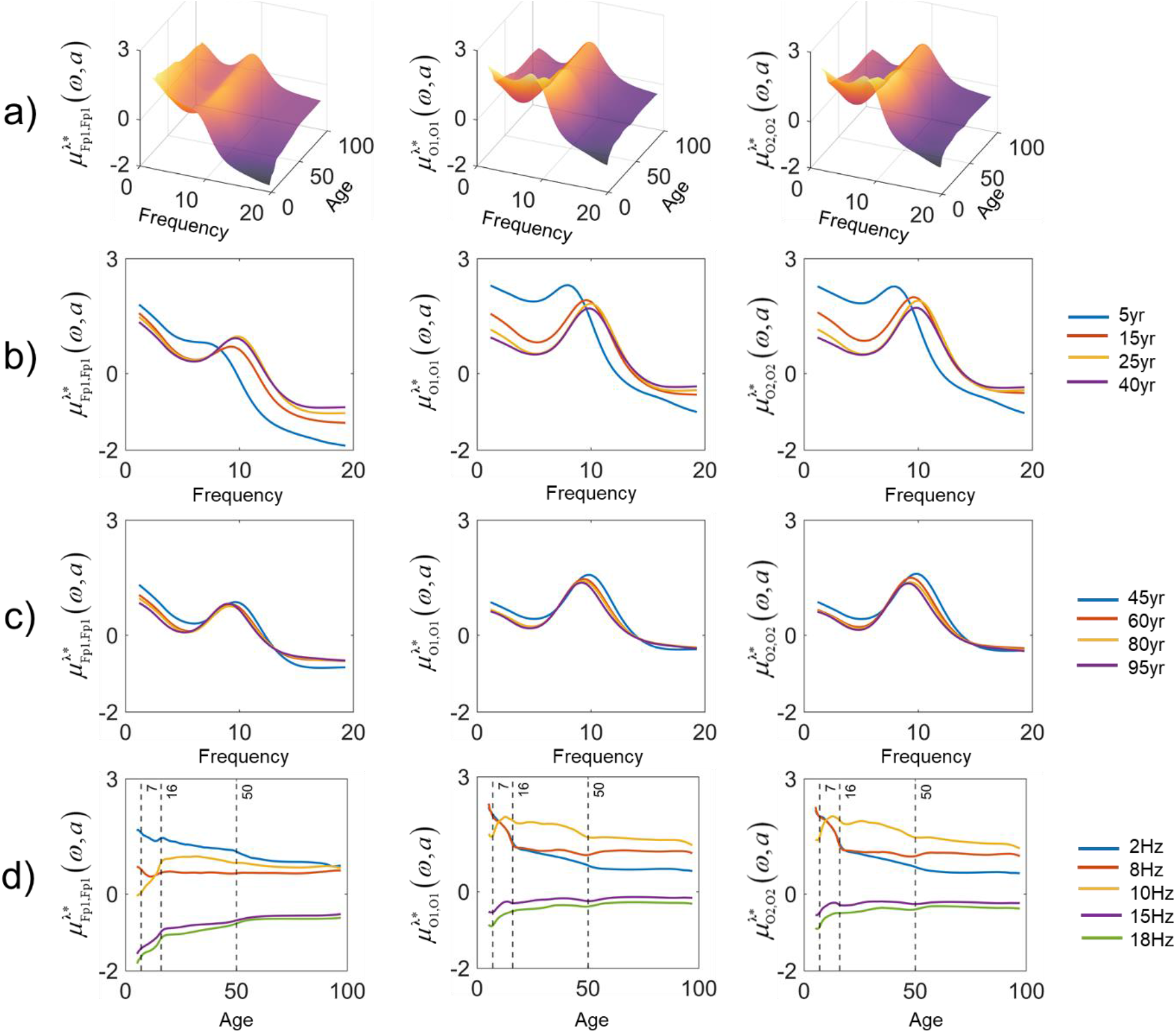
Examples of harmonized normative means and details for traditional log-spectral descriptive parameters as a function of frequency and age. Examples of harmonized normative means and details for traditional log-spectral descriptive parameters as a function of frequency and age. a) Normative means (Developmental surfaces) for channel-Fp1, O1, and O2 of 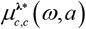 .b) The variation of the norm for fixed ages at younger ages (5 yr, 15 yr, 25 yr, and 40 yr). c) The variation of the norm for fixed ages at elder ages (45yr, 60yr, 80yr, and 95yr age). d) Changes of the norm with age at specific frequencies (2Hz,8Hz,10Hz, 15Hz), with vertical lines at 7yr, 16yr, and 50yr.

**Figure S3.**
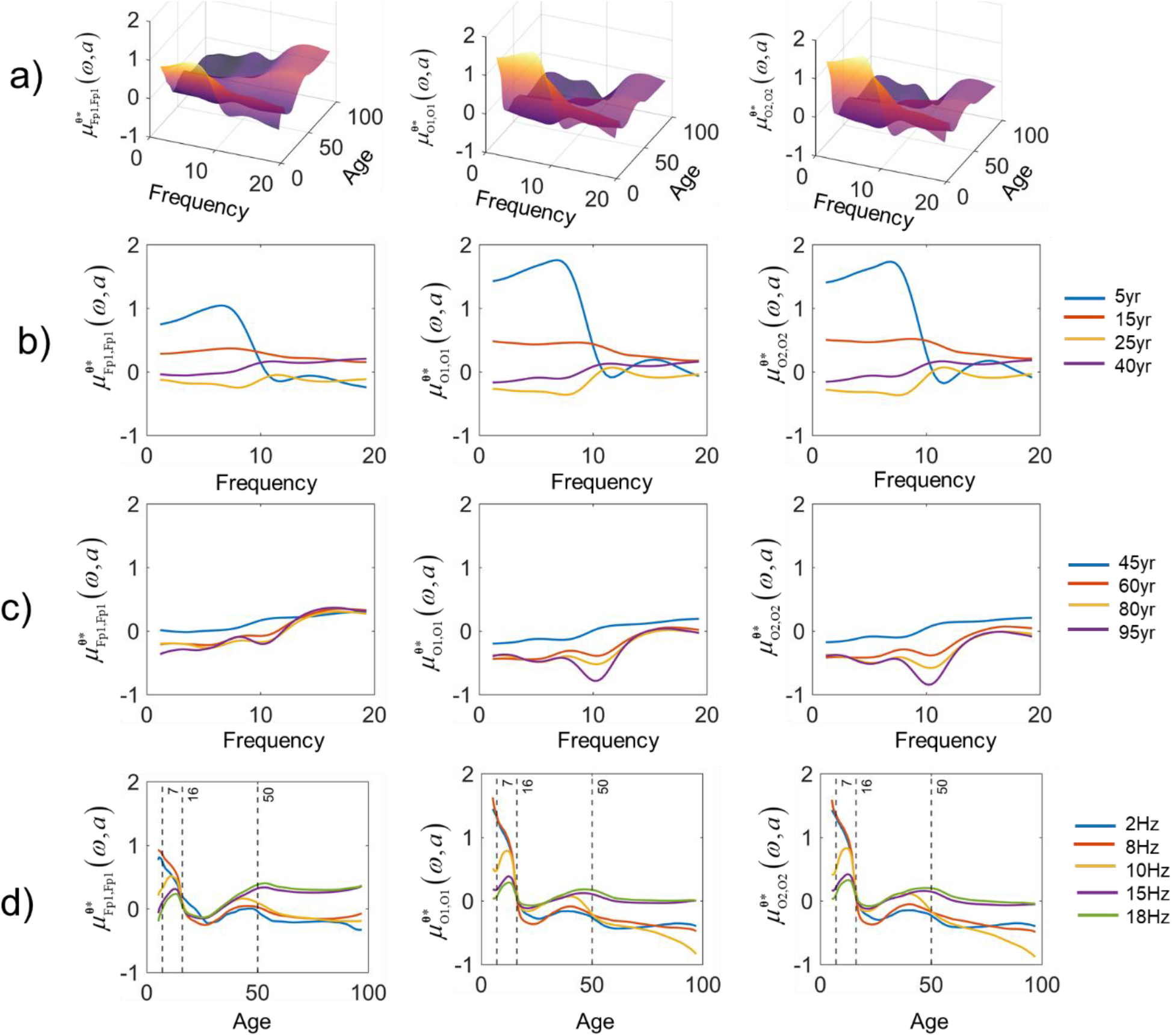
Examples of harmonized normative means and details for Hermitian Riemannian descriptive parameters as a function of frequency and age. Examples of harmonized normative means and details for Hermitian Riemannian descriptive parameters as a function of frequency and age. a) Normative means (Developmental surfaces) for channel-Fp1, O1, and O2 of 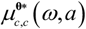. b) The variation of the norm for fixed ages at younger ages (5 yr, 15 yr, 25 yr, and 40 yr). c) The variation of the norm for fixed ages at elder ages (45yr, 60yr, 80yr, and 95yr age). d) Changes of the norm with age at specific frequencies (2Hz,8Hz,10Hz, 15Hz), with vertical lines at 7yr, 16yr, and 50yr.

**Figure S4.**
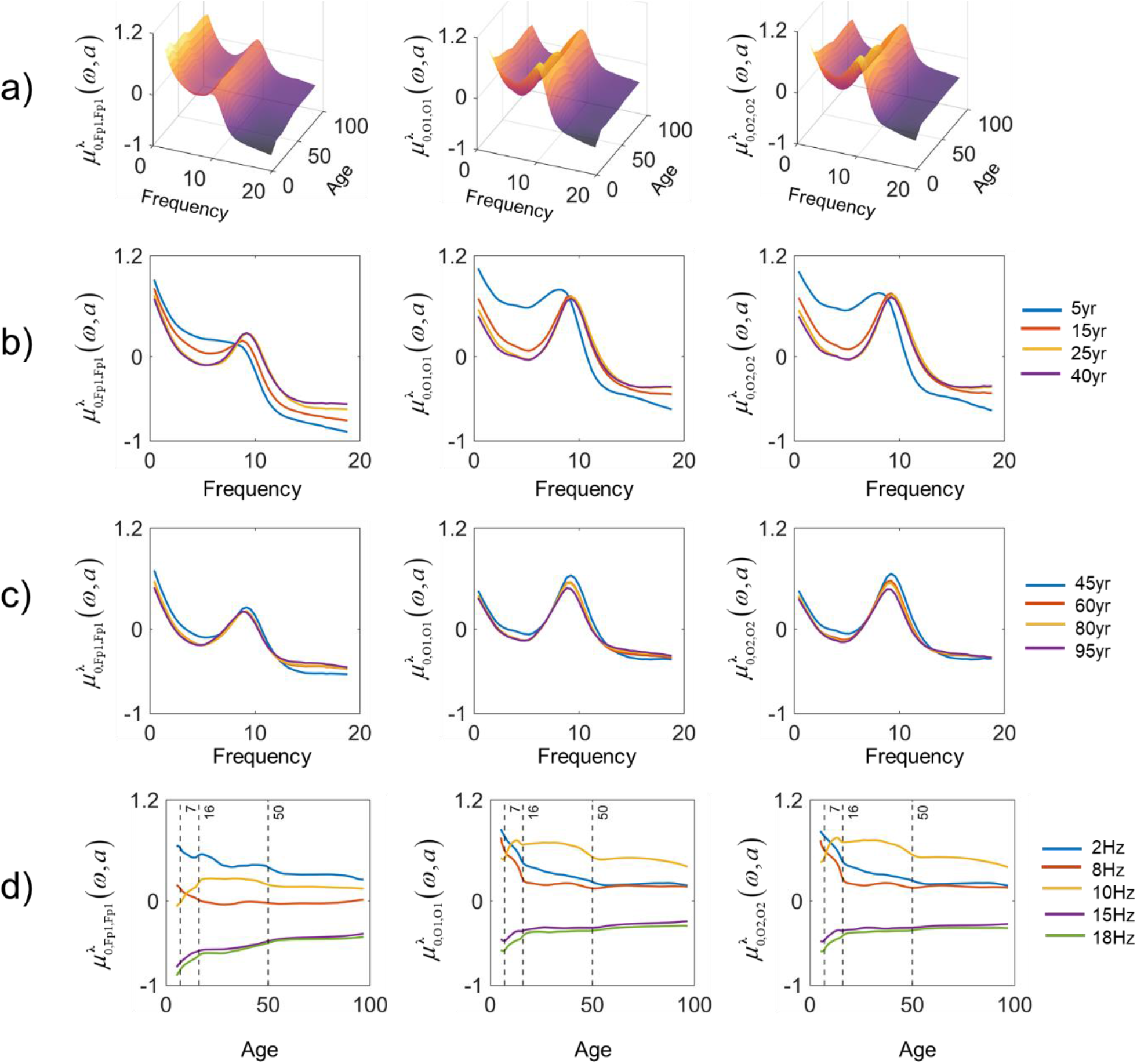
Examples of harmonized normative means and details for surrogate log-spectral norms as a function of frequency and age. Examples of harmonized normative means and details for surrogate log-spectrum norms as a function of frequency and age. a) Normative means (Developmental surfaces) for channel-Fp1, O1, and O2 of 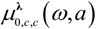. b) The variation of the norm for fixed ages at younger ages (5 yr, 15 yr, 25 yr, and 40 yr). c) The variation of the norm for fixed ages at elder ages (45yr, 60yr, 80yr, and 95yr age). d) Changes of the norm with age at specific frequencies (2Hz,8Hz,10Hz, 15Hz), with vertical lines at 7yr, 16yr, and 50yr.

**Figure S5.**
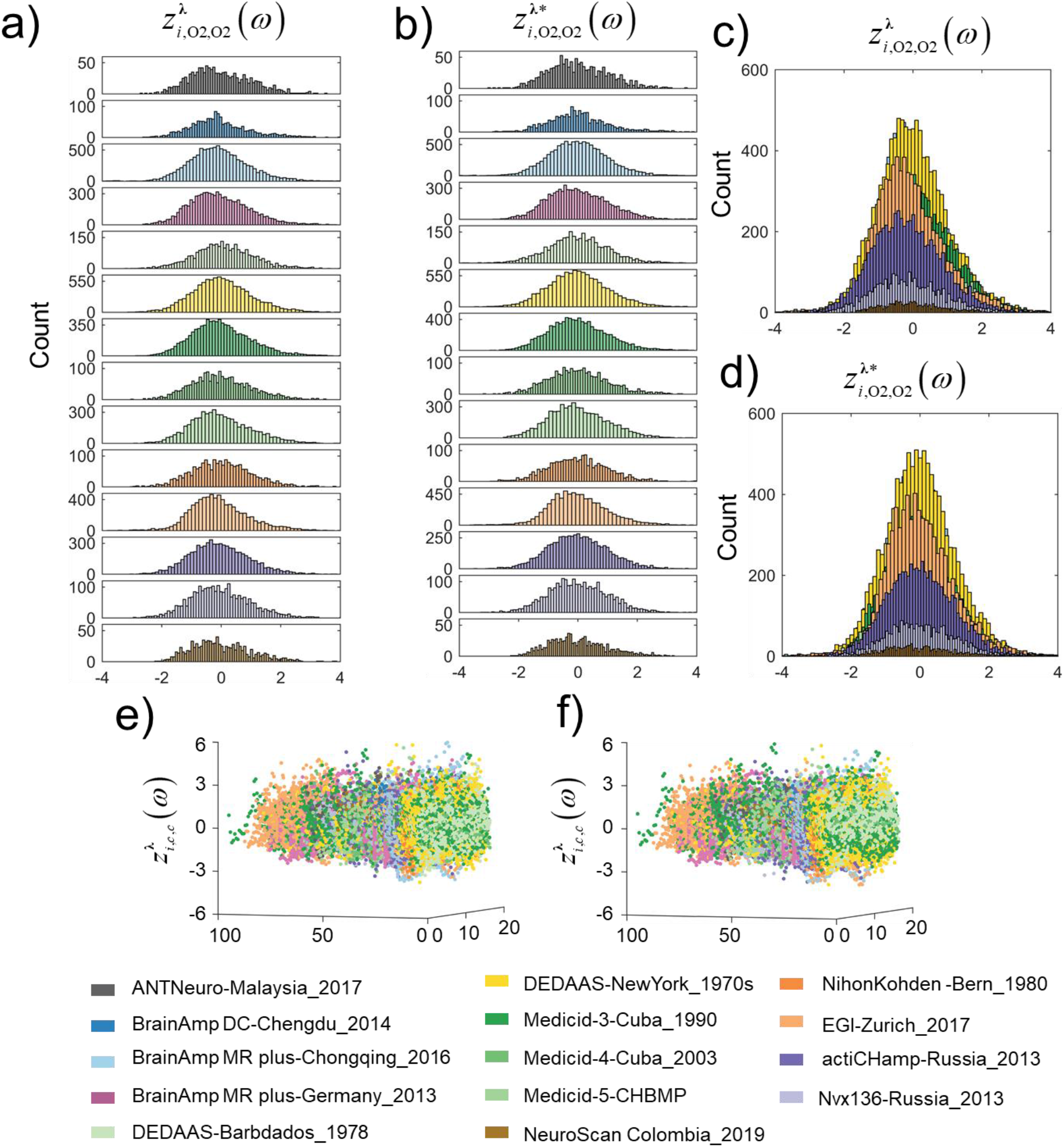
Histograms for 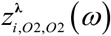 before and after harmonization. Histograms and scatter plot for 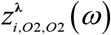 before and after harmonization. Each batch (study) is coded with a different color. a) Histograms of z-score 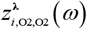 for each batch separately, and c) for all batches superimposed; b) Histograms of batch harmonized traditional log-spectrum DP z-score 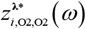 - for each batch separately, and d) with all batches superimposed. e) Scatter plot of 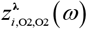 as a function of frequency and age; f) Scatter plot of 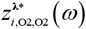 as a function of frequency and age, after harmonization

**Figure S6.**
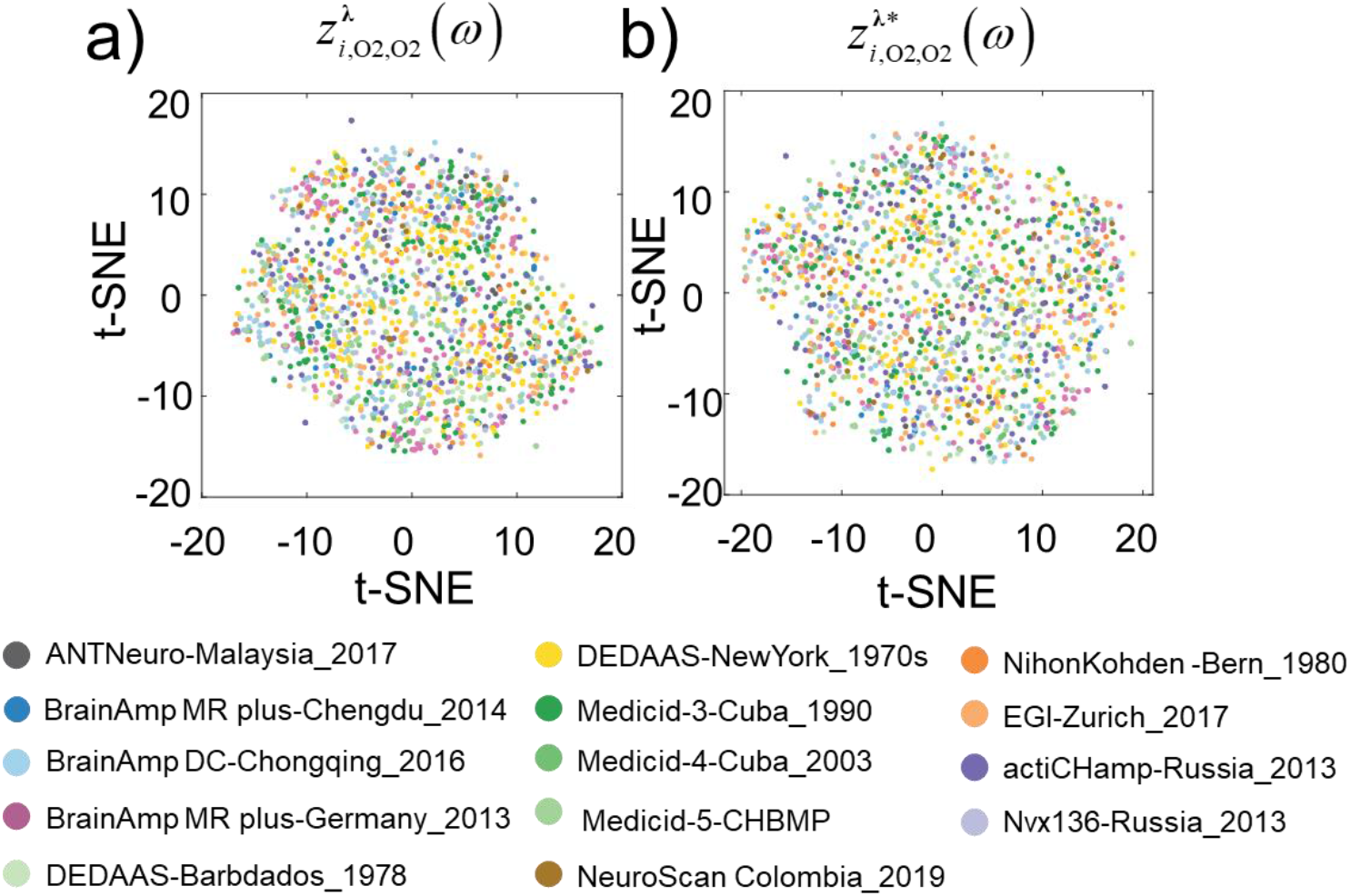
Low dimensional scatter plots of traditional log-spectrum DPs z-scores 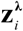, before and after harmonization. Low dimensional scatter plots of log-spectrum z-scores 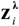, before and after harmonization. The low dimensional representation is a nonlinear mapping (via t-SNE) of z-scores of all traditional log-spectrum DPs onto two dimensions. Each axis is log-transformed. Points represent subjects, colored-coded by batch (study). a) Unharmonized 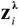 form clusters; b) Harmonized 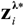 lack cluster structure.

## Reference

Ahn, H., Prichep, L., John, E., Baird, H., Trepetin, M., Kaye, H., 1980. Developmental equations reflect brain dysfunctions. Science 210, 1259–1262. https://doi.org/10.1126/science.7434027

Akaike, H., 1973. Maximum likelihood identification of Gaussian autoregressive moving average models. Biometrika 60, 255–265. https://doi.org/10.1093/biomet/60.2.255

Amador, A.A., Sosa, P.V., Marqui, R.P., Garcia, L.G., Lirio, R.B., Bayard, J.B., 1989. On the structure of EEG development. Electroencephalography and clinical Neurophysiology 73, 10–19.

Babayan, A., Erbey, M., Kumral, D., Reinelt, J.D., Reiter, A.M.F., Röbbig, J., Lina Schaare, H., Uhlig, M., Anwander, A., Bazin, P.L., Horstmann, A., Lampe, L., Nikulin, V. V., Okon-Singer, H., Preusser, S., Pampel, A., Rohr, C.S., Sacher, J., Thöne-Otto, A., Trapp, S., Nierhaus, T., Altmann, D., Arelin, K., Blöchl, M., Bongartz, E., Breig, P., Cesnaite, E., Chen, S., Cozatl, R., Czerwonatis, S., Dambrauskaite, G., Dreyer, M., Enders, J., Engelhardt, M., Fischer, M.M., Forschack, N., Golchert, J., Golz, L., Guran, C.A., Hedrich, S., Hentschel, N., Hoffmann, D.I., Huntenburg, J.M., Jost, R., Kosatschek, A., Kunzendorf, S., Lammers, H., Lauckner, M.E., Mahjoory, K., Kanaan, A.S., Mendes, N., Menger, R., Morino, E., Näthe, K., Neubauer, J., Noyan, H., Oligschläger, S., Panczyszyn-Trzewik, P., Poehlchen, D., Putzke, N., Roski, S., Schaller, M.C., Schieferbein, A., Schlaak, B., Schmidt, R., Gorgolewski, K.J., Schmidt, H.M., Schrimpf, A., Stasch, S., Voss, M., Wiedemann, A., Margulies, D.S., Gaebler, M., Villringer, A., 2019. Data descriptor: A mind-brain-body dataset of MRI, EEG, cognition, emotion, and peripheral physiology in young and old adults. Scientific Data 6, 1–21. https://doi.org/10.1038/sdata.2018.308

Barachant, A., Bonnet, S., Congedo, M., Jutten, C., 2013. Classification of covariance matrices using a Riemannian-based kernel for BCI applications. Neurocomputing, Advances in artificial neural networks, machine learning, and computational intelligence 112, 172–178. https://doi.org/10.1016/j.neucom.2012.12.039

Barachant, A., Bonnet, S., Congedo, M., Jutten, C., 2012. Multiclass Brain–Computer Interface Classification by Riemannian Geometry. IEEE Transactions on Biomedical Engineering 59, 920–928. https://doi.org/10.1109/TBME.2011.2172210

Bethlehem, R. a. I., Seidlitz, J., White, S.R., Vogel, J.W., Anderson, K.M., Adamson, C., Adler, S., Alexopoulos, G.S., Anagnostou, E., Areces-Gonzalez, A., Astle, D.E., Auyeung, B., Ayub, M., Ball, G., Baron-Cohen, S., Beare, R., Bedford, S.A., Benegal, V., Beyer, F., Bae, J.B., Blangero, J., Cábez, M.B., Boardman, J.P., Borzage, M., Bosch-Bayard, J.F., Bourke, N., Calhoun, V.D., Chakravarty, M.M., Chen, C., Chertavian, C., Chetelat, G., Chong, Y.S., Cole, J.H., Corvin, A., Courchesne, E., Crivello, F., Cropley, V.L., Crosbie, J., Crossley, N., Delarue, M., Desrivieres, S., Devenyi, G., Biase, M.A.D., Dolan, R., Donald, K.A., Donohoe, G., Dunlop, K., Edwards, A.D., Elison, J.T., Ellis, C.T., Elman, J.A., Eyler, L., Fair, D.A., Fletcher, P.C., Fonagy, P., Franz, C.E., Galan-Garcia, L., Gholipour, A., Giedd, J., Gilmore, J.H., Glahn, D.C., Goodyer, I., Grant, P.E., Groenewold, N.A., Gunning, F.M., Gur, R.E., Gur, R.C., Hammill, C.F., Hansson, O., Hedden, T., Heinz, A., Henson, R., Heuer, K., Hoare, J., Holla, B., Holmes, A.J., Holt, R., Huang, H., Im, K., Ipser, J., Jack, C.R., Jackowski, A.P., Jia, T., Johnson, K.A., Jones, P.B., Jones, D.T., Kahn, R., Karlsson, H., Karlsson, L., Kawashima, R., Kelley, E.A., Kern, S., Kim, K., Kitzbichler, M.G., Kremen, W.S., Lalonde, F., Landeau, B., Lee, S., Lerch, J., Lewis, J.D., Li, J., Liao, W., Linares, D.P., Liston, C., Lombardo, M.V., Lv, J., Lynch, C., Mallard, T.T., Marcelis, M., Markello, R.D., Mazoyer, B., McGuire, P., Meaney, M.J., Mechelli, A., Medic, N., Misic, B., Morgan, S.E., Mothersill, D., Nigg, J., Ong, M.Q.W., Ortinau, C., Ossenkoppele, R., Ouyang, M., Palaniyappan, L., Paly, L., Pan, P.M., Pantelis, C., Park, M.M., Paus, T., Pausova, Z., Binette, A.P., Pierce, K., Qian, X., Qiu, J., Qiu, A., Raznahan, A., Rittman, T., Rollins, C.K., Romero-Garcia, R., Ronan, L., Rosenberg, M.D., Rowitch, D.H., Salum, G.A., Satterthwaite, T.D., Schaare, H.L., Schachar, R.J., Schultz, A.P., Schumann, G., Schöll, M., Sharp, D., Shinohara, R.T., Skoog, I., Smyser, C.D., Sperling, R.A., Stein, D.J., Stolicyn, A., Suckling, J., Sullivan, G., Taki, Y., Thyreau, B., Toro, R., Tsvetanov, K.A., Turk-Browne, N.B., Tuulari, J.J., Tzourio, C., Vachon-Presseau, É., Valdes-Sosa, M.J., Valdes-Sosa, P.A., Valk, S.L., Amelsvoort, T. van Vandekar, S.N., Vasung, L., Victoria, L.W., Villeneuve, S., Villringer, A., Vértes, P.E., Wagstyl, K., Wang, Y.S., Warfield, S.K., Warrier, V., Westman, E., Westwater, M.L., Whalley, H.C., Witte, A.V., Yang, N., Yeo, B.T.T., Yun, H.J., Zalesky, A., Zar, H.J., Zettergren, A., Zhou, J.H., Ziauddeen, H., Zugman, A., Zuo, X.N., Aibl, Initiative, A.D.N., Investigators, A.D.R.W.B., Asrb, Team, C., Cam-CAN, Ccnp, 3r-Brain, Cobre, Group, E.D.B.A. working, FinnBrain, Study, H.A.B., Imagen, K., Nspn, Oasis-3, Project, O., Pond, The PREVENT-AD Research Group, V., Alexander-Bloch, A.F., 2021. Brain charts for the human lifespan. https://doi.org/10.1101/2021.06.08.447489

Bhatia, R., Holbrook, J., 2006. Riemannian geometry and matrix geometric means. Linear Algebra and its Applications 413, 594–618. https://doi.org/10.1016/j.laa.2005.08.025

Billings, S.A., Tsang, K.M., 1989a. Spectral analysis for non-linear systems, Part I: Parametric non-linear spectral analysis. Mechanical Systems and Signal Processing 3, 319–339. https://doi.org/10.1016/0888-3270(89)90041-1

Billings, S.A., Tsang, K.M., 1989b. Spectral analysis for non-linear systems, Part II: Interpretation of non-linear frequency response functions. Mechanical Systems and Signal Processing 3, 341–359. https://doi.org/10.1016/0888-3270(89)90042-3

Biscay Lirio, R., Valdés Sosa, P.A., Pascual Marqui, R.D., Jiménez-Sobrino, J.C., Alvarez Amador, A., Galán Garcia, L., 1989. Multivariate Box-Cox transformations with applications to neurometric data. Comput Biol Med 19, 263–267. https://doi.org/10.1016/0010-4825(89)90013-9

Bosch-Bayard, J., Galan, L., Aubert Vazquez, E., Virues Alba, T., Valdes-Sosa, P.A., 2020. Resting State Healthy EEG: The First Wave of the Cuban Normative Database. Front. Neurosci. 14. https://doi.org/10.3389/fnins.2020.555119

Bosch-Bayard, J., Galán-García, L., Fernandez, T., Lirio, R.B., Bringas-Vega, M.L., Roca-Stappung, M., Ricardo-Garcell, J., Harmony, T., Valdes-Sosa, P.A., 2018. Stable Sparse Classifiers Identify qEEG Signatures that Predict Learning Disabilities (NOS) Severity. Frontiers in Neuroscience 11, 749. https://doi.org/10.3389/fnins.2017.00749

Bosch-Bayard, J., Valdés-Sosa, P., Virues-Alba, T., Aubert-Vázquez, E., John, E.R., Harmony, T., Riera-Díaz, J., Trujillo-Barreto, N., 2001. 3D Statistical Parametric Mapping of EEG Source Spectra by Means of Variable Resolution Electromagnetic Tomography (VARETA). Clinical Electroencephalography 32, 47–61. https://doi.org/10.1177/155005940103200203

Brillinger, D.R., 1981. Time Series: Data Analysis and Theory. SIAM.

Bringas Vega, M.L., Guo, Y., Tang, Q., Razzaq, F.A., Calzada Reyes, A., Ren, P., Paz Linares, D., Galan Garcia, L., Rabinowitz, A.G., Galler, J.R., Bosch-Bayard, J., Valdes Sosa, P.A., 2019. An Age-Adjusted EEG Source Classifier Accurately Detects School-Aged Barbadian Children That Had Protein Energy Malnutrition in the First Year of Life. Front. Neurosci. 13, 1222. https://doi.org/10.3389/fnins.2019.01222

Chen, J., Chen, Z., 2012. Extended BIC for small-n-large-P sparse GLM. STAT SINICA 22. https://doi.org/10.5705/ss.2010.216

Chen, J., Chen, Z., 2008. Extended Bayesian information criteria for model selection with large model spaces. Biometrika 95, 759–771. https://doi.org/10.1093/biomet/asn034

Chen, L.-H., Cheng, M.-Y., Peng, L., 2009. Conditional variance estimation in heteroscedastic regression models. Journal of Statistical Planning and Inference 139, 236–245. https://doi.org/10.1016/j.jspi.2008.04.020

Craven, P., Wahba, G., 1978. Smoothing noisy data with spline functions. Numer. Math. 31, 377–403. https://doi.org/10.1007/BF01404567

Deza, M.M., Deza, E., 2013. Riemannian and Hermitian Metrics, in: Encyclopedia of Distances. Springer Berlin Heidelberg, Berlin, Heidelberg, pp. 125–155. https://doi.org/10.1007/978-3-642-30958-8_7

Dinga, R., Fraza, C.J., Bayer, J.M.M., Kia, S.M., Beckmann, C.F., Marquand, A.F., 2021. Normative modeling of neuroimaging data using generalized additive models of location scale and shape (preprint). Neuroscience. https://doi.org/10.1101/2021.06.14.448106

Donoghue, T., 2020. Parameterizing neural power spectra into periodic and aperiodic components. Nature Neuroscience 23, 24.

Engemann, D.A., Mellot, A., Höchenberger, R., Banville, H., Sabbagh, D., Gemein, L., Ball, T., Gramfort, A., 2021. A reusable benchmark of brain-age prediction from M/EEG resting-state signals (preprint). Neuroscience. https://doi.org/10.1101/2021.12.14.472691

Fisher, R.A., 1922. On the Interpretation of χ 2 from Contingency Tables, and the Calculation of P. Journal of the Royal Statistical Society 85, 87. https://doi.org/10.2307/2340521

Fortin, J.-P., Cullen, N., Sheline, Y.I., Taylor, W.D., Aselcioglu, I., Cook, P.A., Adams, P., Cooper, C., Fava, M., McGrath, P.J., McInnis, M., Phillips, M.L., Trivedi, M.H., Weissman, M.M., Shinohara, R.T., 2018. Harmonization of cortical thickness measurements across scanners and sites. NeuroImage 167, 104–120. https://doi.org/10.1016/j.neuroimage.2017.11.024

Fritsch, F.N., Carlson, R.E., 2006. Monotone Piecewise Cubic Interpolation. http://dx.doi.org/10.1137/0717021 17, 238–246. https://doi.org/10.1137/0717021

Galler, J.R., Bringas-Vega, M.L., Tang, Q., Rabinowitz, A.G., Musa, K.I., Chai, W.J., Omar, H., Abdul Rahman, M.R., Abd Hamid, A.I., Abdullah, J.M., Valdés-Sosa, P.A., 2021. Neurodevelopmental effects of childhood malnutrition: A neuroimaging perspective. NeuroImage 231, 117828. https://doi.org/10.1016/j.neuroimage.2021.117828

Galler, J.R., Ramsey, F., Solimano, G., Lowell, W.E., 1983a. The Influence of Early Malnutrition on Subsequent Behavioral Development: II. Classroom Behavior. Journal of the American Academy of Child Psychiatry 22, 16–22. https://doi.org/10.1097/00004583-198301000-00003

Galler, J.R., Ramsey, F., Solimano, G., Lowell, W.E., Mason, E., 1983b. The Influence of Early Malnutrition on Subsequent Behavioral Development: I. Degree of Impairment in Intellectual Performance. Journal of the American Academy of Child Psychiatry 22, 8–15. https://doi.org/10.1097/00004583-198301000-00002

Gazula, H., Kelly, R., Romero, J., Verner, E., Baker, B., Silva, R., Imtiaz, H., Saha, D., Raja, R., Turner, J., Sarwate, A., Plis, S., Calhoun, V., 2020. COINSTAC: Collaborative Informatics and Neuroimaging Suite Toolkit for Anonymous Computation. JOSS 5, 2166. https://doi.org/10.21105/joss.02166

Girard, A., 1989. A fast “Monte-Carlo cross-validation” procedure for large least squares problems with noisy data. Numer. Math. 56, 1–23. https://doi.org/10.1007/BF01395775

Gordon, E., Cooper, N., Rennie, C., Hermens, D., Williams, L.M., 2005. Integrative Neuroscience: The Role of a Standardized Database. Clin EEG Neurosci 36, 64–75. https://doi.org/10.1177/155005940503600205

Harmony, T., 2021. Neurometric Assessment of Brain Dysfunction in Neurological Patients. Routledge.

Harmony, T., Alvarez, A., Pascual, R., Ramos, A., Marosi, E., Díaz De León, A.E., Valdés, P., Becker, J., 1988. EEG Maturation on Children with Different Economic and Psychosocial Characteristics. International Journal of Neuroscience 41, 103–113. https://doi.org/10.3109/00207458808985747

Hernández, J.L., Valdés, P., Biscay, R., Virues, T., Szava, S., Bosch, J., Riquenes, A., Clark, I., 1994. A Global Scale Factor in Brain Topography. International Journal of Neuroscience 76, 267–278. https://doi.org/10.3109/00207459408986009

Hernandez-Gonzalez, G., Bringas-Vega, M.L., Galán-Garcia, L., Bosch-Bayard, J., Lorenzo-Ceballos, Y., Melie-Garcia, L., Valdes-Urrutia, L., Cobas-Ruiz, M., Valdes-Sosa, P.A., Cuban Human Brain Mapping Project (CHBMP), 2011. Multimodal Quantitative Neuroimaging Databases and Methods: The Cuban Human Brain Mapping Project. Clin EEG Neurosci 42, 149–159. https://doi.org/10.1177/155005941104200303

Hu, S., Valdes-Sosa, P.A., 2019. Xi rhythms: decoding neural oscillations to create full-brain high-resolution spectra parametric mapping. bioRxiv.

Hu, S., Yao, D., Bringas-Vega, M.L., Qin, Y., Valdes-Sosa, P.A., 2019. The Statistics of EEG Unipolar References: Derivations and Properties. Brain Topogr 32, 696–703. https://doi.org/10.1007/s10548-019-00706-y

John, E.R., 1987. Normative data bank and neurometrics. basic concepts, methods and results of norm constructions. Methods od analysis of brain electrical and magnetic signals. EEG handbook 1, 449–498.

John, E.R., Karmel, B.Z., Corning, W.C., Easton, P., Brown, D., Ahn, H., John, M., Harmony, T., Prichep, L., Toro, A., Gerson, I., Bartlett, F., Thatcher, R., Kaye, H., Valdes, P., Schwartz, E., 1977. Numerical taxonomy identifies different profiles ofbrain functions within groups of behaviorally similar people. 196, 18.

John, E.R., Prichep, L.S., Fridman, J., Easton, P., 1988. Neurometrics: Computer-Assisted Differential Diagnosis of Brain Dysfunctions. Science 239, 162–169. https://doi.org/10.1126/science.3336779

Johnson, W.E., Li, C., Rabinovic, A., 2007. Adjusting batch effects in microarray expression data using empirical Bayes methods. Biostatistics 8, 118–127. https://doi.org/10.1093/biostatistics/kxj037

Kahaner, David., Moler, C.B., Nash, Stephen., 1989. Numerical methods and software 495.

Karcher, H., 1977. Riemannian center of mass and mollifier smoothing. Communications on pure and applied mathematics 30, 509–541.

Koenig, T., Prichep, L., Lehmann, D., Sosa, P.V., Braeker, E., Kleinlogel, H., Isenhart, R., John, E.R., 2002a. Millisecond by Millisecond, Year by Year: Normative EEG Microstates and Developmental Stages. NeuroImage 16, 41–48. https://doi.org/10.1006/nimg.2002.1070

Koenig, T., Prichep, L., Lehmann, D., Sosa, P.V., Braeker, E., Kleinlogel, H., Isenhart, R., John, E.R., 2002b. Millisecond by Millisecond, Year by Year: Normative EEG Microstates and Developmental Stages. NeuroImage 16, 41–48. https://doi.org/10.1006/nimg.2002.1070

Langer, N., von Bastian, C.C., Wirz, H., Oberauer, K., Jäncke, L., 2013. The effects of working memory training on functional brain network efficiency. Cortex 49, 2424–2438. https://doi.org/10.1016/j.cortex.2013.01.008

Leonard, T., Hsu, J.S.J., 2001. Bayesian Methods: An Analysis for Statisticians and Interdisciplinary Researchers. Cambridge University Press.

Leroy, A.M., Rousseeuw, P.J., 1987. Robust regression and outlier detection, Wiley Series in Probability and Mathematical Statistics.

Li, F., Liu, T., Wang, F., Li, H., Gong, D., Zhang, R., Jiang, Y., Tian, Y., Guo, D., Yao, D., Xu, P., 2015. Relationships between the resting-state network and the P3: Evidence from a scalp EEG study. Scientific Reports 5, 15129. https://doi.org/10.1038/srep15129

Lin, X., Zhang, D., 1999. Inference in generalized additive mixed modelsby using smoothing splines. J Royal Statistical Soc B 61, 381–400. https://doi.org/10.1111/1467-9868.00183

Lorensen, T.D., Dickson, P., 2003. Quantitative EEG Normative Databases: A Comparative Investigation. Journal of Neurotherapy 7, 53–68. https://doi.org/10.1300/J184v07n03_03

Makeig, S., Debener, S., Onton, J., Delorme, A., 2004. Mining event-related brain dynamics. Trends Cogn Sci 8, 204–210. https://doi.org/10.1016/j.tics.2004.03.008

Matoušek, M., Petersén, I., 1973a. Automatic evaluation of EEG background activity by means of age-dependent EEG quotients. Electroencephalography and Clinical Neurophysiology 35, 603–612. https://doi.org/10.1016/0013-4694(73)90213-7

Matoušek, M., Petersén, I., 1973b. Automatic evaluation of EEG background activity by means of age-dependent EEG quotients. Electroencephalography and Clinical Neurophysiology 35, 603–612. https://doi.org/10.1016/0013-4694(73)90213-7

McClish, D.K., 1989. Analyzing a Portion of the ROC Curve. Med Decis Making 9, 190–195. https://doi.org/10.1177/0272989X8900900307

Nadaraya, E.A., 1964. On Estimating Regression. Theory Probab. Appl. 9, 141–142. https://doi.org/10.1137/1109020

Olive, D.J., 2004. A resistant estimator of multivariate location and dispersion. Computational Statistics & Data Analysis 46, 93–102. https://doi.org/10.1016/S0167-9473(03)00119-1

Pascual-marqui, R.D., Valdes-sosa, P.A., Alvarez-amador, A., 1988. A parametric model for multichannel EEG spectra. International Journal of Neuroscience 40, 89–99.

Pavlov, Y.G., Adamian, N., Appelhoff, S., Arvaneh, M., Benwell, C.S.Y., Beste, C., Bland, A.R., Bradford, D.E., Bublatzky, F., Busch, N.A., Clayson, P.E., Cruse, D., Czeszumski, A., Dreber, A., Dumas, G., Ehinger, B., Ganis, G., He, X., Hinojosa, J.A., Huber-Huber, C., Inzlicht, M., Jack, B.N., Johannesson, M., Jones, R., Kalenkovich, E., Kaltwasser, L., Karimi-Rouzbahani, H., Keil, A., König, P., Kouara, L., Kulke, L., Ladouceur, C.D., Langer, N., Liesefeld, H.R., Luque, D., MacNamara, A., Mudrik, L., Muthuraman, M., Neal, L.B., Nilsonne, G., Niso, G., Ocklenburg, S., Oostenveld, R., Pernet, C.R., Pourtois, G., Ruzzoli, M., Sass, S.M., Schaefer, A., Senderecka, M., Snyder, J.S., Tamnes, C.K., Tognoli, E., van Vugt, M.K., Verona, E., Vloeberghs, R., Welke, D., Wessel, J.R., Zakharov, I., Mushtaq, F., 2021. #EEGManyLabs: Investigating the replicability of influential EEG experiments. Cortex S0010945221001106. https://doi.org/10.1016/j.cortex.2021.03.013

Pennec, X., 2004. Probabilities and Statistics on Riemannian Manifolds: A Geometric approach (report). INRIA.

Pennec, X., 1999. Intrinsic statistics on Riemannian manifolds: Basic tools for geometric measurements.

Pomponio, R., Erus, G., Habes, M., Doshi, J., Srinivasan, D., Mamourian, E., Bashyam, V., Nasrallah, I.M., Satterthwaite, T.D., Fan, Y., Launer, L.J., Masters, C.L., Maruff, P., Zhuo, C., Völzke, H., Johnson, S.C., Fripp, J., Koutsouleris, N., Wolf, D.H., Gur, Raquel, Gur, Ruben, Morris, J., Albert, M.S., Grabe, H.J., Resnick, S.M., Bryan, R.N., Wolk, D.A., Shinohara, R.T., Shou, H., Davatzikos, C., 2020. Harmonization of large MRI datasets for the analysis of brain imaging patterns throughout the lifespan. NeuroImage 208, 116450. https://doi.org/10.1016/j.neuroimage.2019.116450

Rigby, R.A., Stasinopoulos, D.M., 2005. Generalized additive models for location, scale and shape. Journal of the Royal Statistical Society: Series C (Applied Statistics) 54, 507–554. https://doi.org/10.1111/j.1467-9876.2005.00510.x

Rousseeuw, P.J., Driessen, K.V., 1998. A Fast Algorithm for the Minimum Covariance Determinant Estimator. Technometrics 41, 212–223.

Rutherford, S., Kia, S.M., Wolfers, T., Fraza, C., Zabihi, M., Dinga, R., Berthet, P., Worker, A., Verdi, S., Ruhe, H.G., Beckmann, C.F., Marquand, A.F., 2021a. The Normative Modeling Framework for Computational Psychiatry (preprint). Neuroscience. https://doi.org/10.1101/2021.08.08.455583

Rutherford, S., Kia, S.M., Wolfers, T., Fraza, C., Zabihi, M., Dinga, R., Berthet, P., Worker, A., Verdi, S., Ruhe, H.G., Beckmann, C.F., Marquand, A.F., 2021b. The Normative Modeling Framework for Computational Psychiatry (preprint). Neuroscience. https://doi.org/10.1101/2021.08.08.455583

Sabbagh, D., Ablin, P., Varoquaux, G., Gramfort, A., Engemann, D.A., 2020. Predictive regression modeling with MEG/EEG: from source power to signals and cognitive states. NeuroImage 222, 116893. https://doi.org/10.1016/j.neuroimage.2020.116893

Sabbagh, D., Ablin, P., Varoquaux, G., Gramfort, A., Engemann, D.A., 2019. Predictive regression modeling with MEG/EEG: from source power to signals and cognitive states (preprint). Neuroscience. https://doi.org/10.1101/845016

Schneider-Luftman, D., Walden, A.T., 2016. Partial Coherence Estimation via Spectral Matrix Shrinkage under Quadratic Loss. IEEE Trans. Signal Process. 64, 5767–5777. https://doi.org/10.1109/TSP.2016.2582464

Schwarz, G., 1978. Estimating the Dimension of a Model. The Annals of Statistics 6, 461–464.

Simeon, G., Piella, G., Camara, O., Pareto, D., 2021. Riemannian geometry of functional connectivity matrices for multi-site attention-deficit/hyperactivity disorder data harmonization (preprint). Neuroscience. https://doi.org/10.1101/2021.09.01.458579

Stone, M., 1974. Cross-validation and multinomial prediction. Biometrika 61, 509–515. https://doi.org/10.1093/biomet/61.3.509

Szava, S., Valdes, P., Biscay, R., Galan, L., Bosch, J., Clark, I., Jimenez, J.C., 1994. High resolution quantitative EEG analysis. Brain Topogr 6, 211–219. https://doi.org/10.1007/BF01187711

Taboada-Crispi, A., Bringas-Vega, M.L., Bosch-Bayard, J., Galán-García, L., Bryce, C., Rabinowitz, A.G., Prichep, L.S., Isenhart, R., Calzada-Reyes, A., VIrues-Alba, T., Guo, Y., Galler, J.R., Valdés-Sosa, P.A., 2018. Quantitative EEG Tomography of Early Childhood Malnutrition. Front. Neurosci. 12. https://doi.org/10.3389/fnins.2018.00595

Thatcher, R.W., Walker, R.A., Biver, C.J., North, D.N., Curtin, R., 2003. Quantitative EEG Normative Databases: Validation and Clinical Correlation. Journal of Neurotherapy 7, 87–121. https://doi.org/10.1300/J184v07n03_05

Thompson, P.M., Stein, J.L., Medland, S.E., Hibar, D.P., Vasquez, A.A., Renteria, M.E., Toro, R., Jahanshad, N., the Alzheimer’s Disease Neuroimaging Initiative, EPIGEN Consortium, IMAGEN Consortium, Saguenay Youth Study (SYS) Group, Schumann, G., Franke, B., Wright, M.J., Martin, N.G., Agartz, I., Alda, M., Alhusaini, S., Almasy, L., Almeida, J., Alpert, K., Andreasen, N.C., Andreassen, O.A., Apostolova, L.G., Appel, K., Armstrong, N.J., Aribisala, B., Bastin, M.E., Bauer, M., Bearden, C.E., Bergmann, Ø., Binder, E.B., Blangero, J., Bockholt, H.J., Bøen, E., Bois, C., Boomsma, D.I., Booth, T., Bowman, I.J., Bralten, J., Brouwer, R.M., Brunner, H.G., Brohawn, D.G., Buckner, R.L., Buitelaar, J., Bulayeva, K., Bustillo, J.R., Calhoun, V.D., Cannon, D.M., Cantor, R.M., Carless, M.A., Caseras, X., Cavalleri, G.L., Chakravarty, M.M., Chang, K.D., Ching, C.R.K., Christoforou, A., Cichon, S., Clark, V.P., Conrod, P., Coppola, G., Crespo-Facorro, B., Curran, J.E., Czisch, M., Deary, I. J., de Geus, E.J.C., den Braber, A., Delvecchio, G., Depondt, C., de Haan, L., de Zubicaray, G.I., Dima, D., Dimitrova, R., Djurovic, S., Dong, H., Donohoe, G., Duggirala, R., Dyer, T.D., Ehrlich, S., Ekman, C.J., Elvsåshagen, T., Emsell, L., Erk, S., Espeseth, T., Fagerness, J., Fears, S., Fedko, I., Fernández, G., Fisher, S.E., Foroud, T., Fox, P.T., Francks, C., Frangou, S., Frey, E.M., Frodl, T., Frouin, V., Garavan, H., Giddaluru, S., Glahn, D.C., Godlewska, B., Goldstein, R.Z., Gollub, R. L., Grabe, H.J., Grimm, O., Gruber, O., Guadalupe, T., Gur, R.E., Gur, R.C., Göring, H.H.H., Hagenaars, S., Hajek, T., Hall, G.B., Hall, J., Hardy, J., Hartman, C. A., Hass, J., Hatton, S.N., Haukvik, U.K., Hegenscheid, K., Heinz, A., Hickie, I.B., Ho, B.-C., Hoehn, D., Hoekstra, P.J., Hollinshead, M., Holmes, A.J., Homuth, G., Hoogman, M., Hong, L.E., Hosten, N., Hottenga, J.-J., Hulshoff Pol, H.E., Hwang, K.S., Jack, C.R., Jenkinson, M., Johnston, C., Jönsson, E.G., Kahn, R.S., Kasperaviciute, D., Kelly, S., Kim, S., Kochunov, P., Koenders, L., Krämer, B., Kwok, J. B.J., Lagopoulos, J., Laje, G., Landen, M., Landman, B.A., Lauriello, J., Lawrie, S. M., Lee, P.H., Le Hellard, S., Lemaître, H., Leonardo, C.D., Li, C., Liberg, B., Liewald, D.C., Liu, X., Lopez, L.M., Loth, E., Lourdusamy, A., Luciano, M., Macciardi, F., Machielsen, M.W.J., MacQueen, G.M., Malt, U.F., Mandl, R., Manoach, D.S., Martinot, J.-L., Matarin, M., Mather, K.A., Mattheisen, M., Mattingsdal, M., Meyer-Lindenberg, A., McDonald, C., McIntosh, A.M., McMahon, F.J., McMahon, K.L., Meisenzahl, E., Melle, I., Milaneschi, Y., Mohnke, S., Montgomery, G.W., Morris, D. W., Moses, E.K., Mueller, B.A., Muñoz Maniega, S., Mühleisen, T.W., Müller-Myhsok, B., Mwangi, B., Nauck, M., Nho, K., Nichols, T.E., Nilsson, L.-G., Nugent, A.C., Nyberg, L., Olvera, R.L., Oosterlaan, J., Ophoff, R.A., Pandolfo, M., Papalampropoulou-Tsiridou, M., Papmeyer, M., Paus, T., Pausova, Z., Pearlson, G.D., Penninx, B.W., Peterson, C.P., Pfennig, A., Phillips, M., Pike, G.B., Poline, J.-B., Potkin, S.G., Pütz, B., Ramasamy, A., Rasmussen, J., Rietschel, M., Rijpkema, M., Risacher, S.L., Roffman, J.L., Roiz-Santiañez, R., Romanczuk-Seiferth, N., Rose, E.J., Royle, N.A., Rujescu, D., Ryten, M., Sachdev, P.S., Salami, A., Satterthwaite, T.D., Savitz, J., Saykin, A.J., Scanlon, C., Schmaal, L., Schnack, H.G., Schork, A.J., Schulz, S.C., Schür, R., Seidman, L., Shen, L., Shoemaker, J.M., Simmons, A., Sisodiya, S.M., Smith, C., Smoller, J.W., Soares, J.C., Sponheim, S.R., Sprooten, E., Starr, J.M., Steen, V.M., Strakowski, S., Strike, L., Sussmann, J., Sämann, P.G., Teumer, A., Toga, A.W., Tordesillas-Gutierrez, D., Trabzuni, D., Trost, S., Turner, J., Van den Heuvel, M., van der Wee, N.J., van Eijk, K., van Erp, T.G.M., van Haren, N. E.M., van ‘t Ent, D., van Tol, M.-J., Valdés Hernández, M.C., Veltman, D.J., Versace, A., Völzke, H., Walker, R., Walter, H., Wang, L., Wardlaw, J.M., Weale, M.E., Weiner, M.W., Wen, W., Westlye, L.T., Whalley, H.C., Whelan, C.D., White, T., Winkler, A.M., Wittfeld, K., Woldehawariat, G., Wolf, C., Zilles, D., Zwiers, M.P., Thalamuthu, A., Schofield, P.R., Freimer, N.B., Lawrence, N.S., Drevets, W., 2014. The ENIGMA Consortium: large-scale collaborative analyses of neuroimaging and genetic data. Brain Imaging and Behavior 8, 153–182. https://doi.org/10.1007/s11682-013-9269-5

Tröndle, M., Popov, T., Langer, N., 2020. Decomposing the role of alpha oscillations during brain maturation. https://doi.org/10.1101/2020.11.06.370882

Tröndle, M., Popov, T., Pedroni, A., Pfeiffer, C., Barańczuk-Turska, Z., Langer, N., 2021. Decomposing age effects in EEG alpha power (preprint). Neuroscience. https://doi.org/10.1101/2021.05.26.445765

Turlach, B.A., Wand, M.P., 1996. Fast computation of auxiliary quantities in local polynomial regression. Journal of Computational and Graphical Statistics 5, 337–350.

Valdés, P., Bosch, J., Grave, R., Hernandez, J., Riera, J., Pascual, R., Biscay, R., 1992. Frequency domain models of the EEG. Brain topography 4, 309–319.

Valdés-Sosa, P.A., Evans, A.C., Valdes-Sosa, M., Mu-ming, P., 2021. A Call for International Research on COVID Induced Brain Disorders. National Science Review. https://doi.org/10.1093/nsr/nwab190

Valdes-Sosa, P.A., Galan-Garcia, L., Bosch-Bayard, J., Bringas-Vega, M.L., Aubert-Vazquez, E., Rodriguez-Gil, I., Das, S., Madjar, C., Virues-Alba, T., Mohades, Z., MacIntyre, L.C., Rogers, C., Brown, S., Valdes-Urrutia, L., Evans, A.C., Valdes-Sosa, M.J., 2021. The Cuban Human Brain Mapping Project, a young and middle age population-based EEG, MRI, and cognition dataset. Scientific Data 8, 45. https://doi.org/10.1038/s41597-021-00829-7

Van der Maaten, L., Hinton, G., 2008. Visualizing data using t-SNE. Journal of machine learning research 9.

Voytek, B., Kramer, M.A., Case, J., Lepage, K.Q., Tempesta, Z.R., Knight, R.T., Gazzaley, A., 2015. Age-Related Changes in 1/f Neural Electrophysiological Noise. Journal of Neuroscience 35, 13257–13265. https://doi.org/10.1523/JNEUROSCI.2332-14.2015

Wand, M.P., 1994. Fast Computation of Multivariate Kernel Estimators. null 3, 433–445. https://doi.org/10.1080/10618600.1994.10474656

Yger, F., Berar, M., Lotte, F., 2017. Riemannian Approaches in Brain-Computer Interfaces: A Review. IEEE Trans. Neural Syst. Rehabil. Eng. 25, 1753–1762. https://doi.org/10.1109/TNSRE.2016.2627016

Zetterberg, L.H., 1969. Estimation of parameters for a linear difference equation with application to EEG analysis. Mathematical Biosciences 5, 227–275. https://doi.org/10.1016/0025-5564(69)90044-3

## Reference of Appendix

Koenig, T., Prichep, L., Lehmann, D., Sosa, P.V., Braeker, E., Kleinlogel, H., Isenhart, R., John, E.R., 2002. Millisecond by Millisecond, Year by Year: Normative EEG Microstates and Developmental Stages. NeuroImage 16, 41–48. https://doi.org/10.1006/nimg.2002.1070

